# Increased SGK1 activity potentiates mineralocorticoid/NaCl-induced hypertension and kidney injury

**DOI:** 10.1101/2020.07.08.191874

**Authors:** Catalina Sierra-Ramos, Silvia Velazquez-Garcia, Ayse G. Keskus, Arianna Vastola-Mascolo, Ana E. Rodríguez-Rodríguez, Sergio Luis-Lima, Guadalberto Hernández, Juan F. Navarro-González, Esteban Porrini, Ozlen Konu, Diego Alvarez de la Rosa

## Abstract

The serum and glucocorticoid-induced kinase 1 (SGK1) stimulates aldosterone-dependent renal Na^+^ reabsorption and modulates blood pressure. In addition, genetic ablation or pharmacological inhibition of SGK1 limits the development of kidney inflammation and fibrosis in response to excess mineralocorticoid signalling. In this work we tested the hypothesis that a systemic increase in SGK1 activity would potentiate mineralocorticoid/salt-induced hypertension and kidney injury. To that end, we used a transgenic mouse model with increased SGK1 activity. Mineralocorticoid/salt-induced hypertension and kidney damage was induced by unilateral nephrectomy and treatment with deoxycorticosterone acetate and NaCl in drinking water for six weeks. Our results demonstrate higher systolic blood pressure in treated transgenic mice when compared to wild type counterparts. Transgenic mice showed a mild increase in glomerular filtration rate, increased albuminuria, exacerbated glomerular hypertrophy and fibrosis. Transcriptomic analysis showed that extracellular matrix and immune response related terms were enriched in the downregulated and upregulated genes, respectively, in transgenic mice. In conclusion, we propose that systemically increased SGK1 activity is a risk factor for the development of mineralocorticoid-dependent hypertension and incipient kidney injury in the context of low renal mass.

## INTRODUCTION

Inappropriately increased mineralocorticoid receptor (MR) signalling plays an important role in the development of chronic kidney disease (CKD) [1]. It has long been known that patients with primary aldosteronism develop hypertension and renal damage. Initially, renal damage was ascribed to increased blood pressure, but it has later been shown that these patients develop kidney injury to a larger extent than comparable individuals with primary hypertension [2]. Extensive work on animal models support a role for aldosterone excess and MR signalling in hypertension-dependent and - independent renal damage and progression to CKD [1, 3, 4]. Importantly, preclinical studies conclusively show a major role for MR antagonists in preventing or limiting kidney injury in a large variety of kidney disease models, including hypertension or diabetic nephropathy, glomerulonephritis or transition from acute kidney injury to CKD [3]. These observations have been translated to humans, with data from clinical trials supporting the use of MR inhibition in limiting proteinuria and eliciting renoprotection [3].

The pathophysiological mechanisms involved in the deleterious effects of aldosterone/MR on the kidney expand beyond the role of this system in regulating tubular Na^+^ reabsorption. MR expression has been detected not only in the distal nephron epithelium, but also in podocytes, mesangial cells, endothelial and smooth muscle cells, interstitial fibroblasts and macrophages [3]. Activation of MR in these cell types have been implicated in alterations in renal circulation, oxidative stress, inflammation, glomerular and interstitial fibrosis and alteration of glomerular filtration barrier [1, 3, 4].

From a molecular perspective, aldosterone/MR-mediated kidney injury likely involves a multiplicity of direct and indirect mechanisms [3]. MR is a nuclear receptor and it is generally assumed that its main long-term effects are mediated by modulation of gene transcription. However, there is still an important knowledge gap about MR gene targets mediating the effects. One interesting candidate to mediate at least part of the processes leading to MR-induced kidney injury and progression to CKD is the serum and glucocorticoid-induced kinase 1 (SGK1), a member of the AGC family of S/T kinases. SGK1 is activated by insulin and several growth factors through the phosphatidylinositol 3-kinase (PI3K) pathway, involving downstream kinases 3-phosphoinositide-dependent kinase 1 (PDK1) and mammalian target of rapamycin complex 2 (mTORC2) [5]. SGK1 transcription and translation are tightly regulated and its expression is enhanced by various stimuli including glucocorticoids and mineralocorticoids, being a direct target gene of both MR and the glucocorticoid receptor (GR). The kinase regulates subcellular trafficking of many ion channels and transporters [6], increasing renal Na^+^ reabsorption [7, 8] and K^+^ excretion [9], thus participating in the control of blood pressure. *Sgk1* knockout protects mice against salt-induced hypertension in a high-fat diet context [10]. Polymorphisms in the human *SGK1* gene has been linked to increased blood pressure [11], particularly associated to hyperinsulinemia [12] and the effects of dietary salt intake on blood pressure [13, 14]. In addition, variants of SGK1 associate to insulin secretion [15] and type 2 diabetes [16]. Consistently, SGK1 inhibition reduces blood pressure in hyperinsulinemic mice [17] and transgenic SGK1 activation potently induces hypertension in mice treated with a high-fat diet [18].

In addition to regulating transepithelial ion transport, SGK1 has additional cellular functions through direct phosphorylation of signalling molecules such as glycogen synthase kinase-3β (GSK-3β) or transcription factors such as members of the FOXO family [19], a pathway that promotes cell survival and proliferation. SGK1 has been implicated in the development of inflammatory and fibrotic processes [20]. Increased SGK1 expression has been described in kidney biopsies from patients with heavy proteinuria [21]. Mice lacking SGK1 show decreased proteinuria and renal fibrosis in response to deoxycorticosterone acetate (DOCA)/NaCl treatment [22, 23], suggesting that this gene is key to develop kidney injury in response to excess mineralocorticoids. SGK1 has also been associated to aldosterone-induced oxidative stress and podocyte injury [24]. SGK1 is also expressed in antigen-presenting cells [25] and is important during differentiation of T cells [26], playing a key role in salt-sensitive hypertension and renal inflammation [20, 25–27], which further supports a pleiotropic role for SGK1 in kidney disease.

Altogether, the available evidence suggests that increased systemic SGK1 activity could be a common factor underlying blood pressure-sensitive and insensitive pathways leading to renal damage and thus accelerate progression to CKD. To test this hypothesis we took advantage of a transgenic mouse model (Tg.sgk1) previously developed in our laboratory using a modified mouse bacterial artificial chromosome (BAC) that drives the expression of a constitutively active mutant (SGK1-S422D) under its own promoter. This transgenic strain provides a model of generalized increase in SGK1 activity without overexpression of the kinase [18, 28, 29]. We used a well-established model of mineralocorticoid excess and high NaCl intake in the context of uninephrectomy to induce hypertension and renal damage [30, 31]. Our results show that increased SGK1 activity induced higher systolic blood pressure after treatment. In addition, transgenic mice displayed exacerbated glomerular hypertrophy, proteinuria and fibrosis with conserved glomerular filtration rate. Finally, we used transcriptomic analysis to define SGK1-modulated pathways in this model. In conclusion, we propose that systemically increased SGK1 activity could be a risk factor for the development of mineralocorticoid-dependent hypertension and kidney injury in the context of low renal mass.

## MATERIAL AND METHODS

### Transgenic expression of constitutively active SGK1 in mice

The generation of transgenic mice expressing a constitutively active mutant of SGK1 (S422D) has been previously described [18, 28, 29]. Briefly, a BAC containing 180 kb of mouse genomic DNA that includes the full SGK1 gene but no other known or predicted genes was obtained from the BACPAC Resources Center (Children’s Hospital Oakland Research Institute, Oakland, CA) and modified by homologous recombination in *Escherichia coli* to add a constitutively activating point mutation (S422D). Founder animals harbouring the transgene were backcrossed with C57BL/6 mice for eight generations. Mice homozygous for the transgene were obtained by crossing heterozygous animals.

### Animal procedures

All experiments involving mice followed protocols previously approved by the University of La Laguna Ethics Committee on Research and Animal Welfare and were performed in accordance with the European Community guidelines for the use of experimental animals and with Spanish law for the protection of animals. Mice were kept in a 12-hour light/dark cycle at 22°C with *ad libitum* access to food and water. Unilateral nephrectomy (NPX) was performed under isoflurane anaesthesia administered with a vaporizer (5% for induction, 2.5% for maintenance during surgery).

Buprenorphine (75 μg/kg body weight [BW]) was administered subcutaneously for peri-operative pain relief. After a two-week recovery period, NPX animals were divided into two groups and treated or not with 1% NaCl in drinking water and 75 mg/kg BW deoxycorticosterone acetate (DOCA) dissolved in sunflower oil and administered subcutaneously three times a week for six weeks [32] (NDS group). Untreated animals were injected three times weekly with vehicle. Animals from the four experimental groups were housed in metabolic cages. After a two-day adaptation period samples were collected for two consecutive days. Food and water intake and urinary and faecal output were measured daily. Urine samples were centrifuged for 10 min at 13000 xg and stored at −80°C for further analysis. Urinary Na^+^, K^+^ and Cl^−^ content was measured using ion-selective electrodes (Cobas c501, Roche Diagnostics). Urinary creatinine and albumin concentration were measured using commercially available kits (Creatinine Assay Kit and Mouse Albumin ELISA, Abcam). Systolic blood pressure (SBP) was measured by tail cuff plethysmography in trained conscious mice using a non-invasive blood pressure measuring system for rodents (model LE5007, Harvard Apparatus Panlab). Data was acquired with SeDaCom 2.0 software. Measurements were collected for three consecutive days after five days of training. SBP was measured at 20 min intervals for 1 h between 10 and 11 a.m. GFR measurements used a simplified method of iohexol plasma clearance that we have previously validated in mice [33]. Briefly, a single dose of iohexol (6,47 mg Omnipaque 300, GE Healthcare) was injected intravenously into the tail vein in mice lightly sedated with isoflurane (2.5%), administered by facemask while the tail vein was dilated using a heating blanket. Blood samples (≤10 μl) were collected at 15, 35, 55, and 75 min after injection from a small cut in the tail using a heparinized capillary tube. Plasma iohexol concentrations were measured by HPLC. Internal calibration curves of iohexol were prepared for each set of samples. Plasma concentrations of iohexol were recalculated from blood levels using the formula: C_plasma_= C_blood_/1-Hct where Hct is the haematocrit. Measured iohexol concentrations were fitted by a slope-intercept method according to a one-compartment model simplified method [33]. Fractional excretion of Na^+^ (FE_Na+_) was expressed as the percentage of the filtered Na^+^ load (calculated from plasma Na^+^ concentration and GFR) excreted in the urine [34].

### Tissue sampling, plasma measurements, histological analysis and cell culture

After treatments mice were euthanized by carbon dioxide inhalation, and blood was collected by cardiac puncture. Serum corticosterone and aldosterone levels were determined with commercial ELISA kits (DRG Diagnostics). Plasma Na^+^, K^+^ and Cl^−^ concentration was measured using ion-selective electrodes (Cobas c501, Roche Diagnostics). Tissues were collected from control and treatment groups at the same time and under the same conditions and processed and analysed in parallel. Kidneys were removed, weighed and frozen in liquid nitrogen and stored at −80°C for molecular analysis or fixed by immersion in 4% formaldehyde, paraffin embedded and processed for histology. Kidney 3 μm-thick microtome sections were stained with 0.1% Sirius red in saturated picric acid. Sections were codified to conduct a blinded quantification of fibrosis area. To that end, images from 5 different fields in each section were collected at 40x magnification and fibrotic area was quantified using Fiji software (ImageJ, National Institutes of Health [35]). Two independent experiments were analysed (first experiment, N=7 animals in each group; second experiment, N=5 animals in each group). Primary mouse embryonic fibroblasts (MEFs) were derived from WT and Tg.sgk1 embryos and cultured as described [36].

### Microarray analysis and quantitative RT-PCR (qPCR)

Total RNA was obtained from tissues disrupted with a Tissue Tearor (BioSpec Products) or from cultured cells using a commercial kit (Total RNA Spin Plus, REAL, Valencia, Spain), which includes an in-column treatment with DNase I. Purified total RNA was quantified using a Nanodrop 1000 (Thermo Scientific). Whole-transcriptome analysis was performed using RNA from kidney cortex of nephrectomized, DOCA/NaCl-treated WT and Tg.sgk1 mice (N=4 in each group). RNA quality analysis was performed using an Agilent 2100 Bioanalyzer. cDNA was synthesized using the GeneChip® WT PLUS Reagent Kit (Affymetrix) and analysed by hybridization to a GeneChip® Mouse Gene 2.0 ST Array (Affymetrix). MIAME-compliant microarray data has been deposited in the Gene Expression Omnibus repository (GEO, National Center for Biotechnology Information, National Institutes of Health) with accession number GSE148880. Raw files were normalized with rma using oligo package in R [37], while differential expression analysis was performed using limma package [38]. A cut-off level of 0.15 (FDR) was used for obtaining the volcano plot (https://github.com/kevinblighe/EnhancedVolcano) and STRING protein-protein interaction network [39], which were visualized in Cytoscape [40]. Gene Ontology (GO) [41] enrichment analysis was performed with an FDR = 0.3 cut-off using clusterProfiler package in R [42]. Relative mRNA abundance was examined by quantitative RT-PCR (qPCR). Equal amounts of total RNA were processed with the iScript cDNA synthesis kit (Bio-Rad). Gene expression was assessed using commercially available Taqman primer/probe sets (Thermo Fischer) or specific oligonucleotide pairs (Table 1) and SYBR® Green supermix (Bio-Rad). Relative expression of the target genes was calculated with the comparative C_t_ method [43] using glyceraldehyde-3-phosphate dehydrogenase (GAPDH) as normalizer.

**Table 1.**
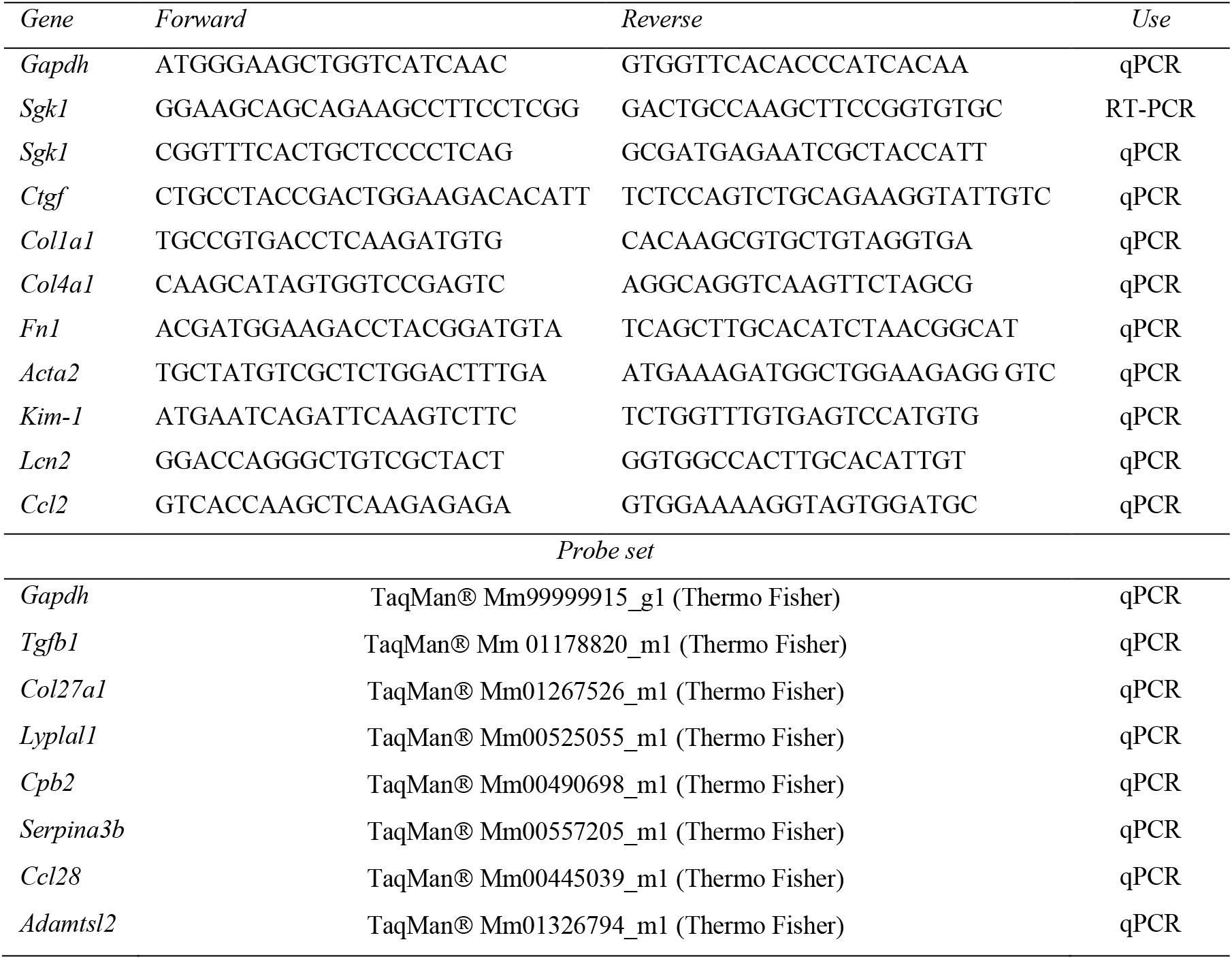
Primers and quantitative RT-PCR probesets used in this study. In the case of TaqMan® probe sets, catalog numbers are indicated. *Gapdh*, glyceraldehyde-3-phosphate dehydrogenase; *Acta2*, smooth muscle actin, alpha 2; *Ctgf*, connective tissue growth factor; *Col1a1*, collagen type I, alpha 1; *Col4a1*, collagen type IV, alpha 1; *Fn1*, fibronectin 1; *Kim-1*, kidney injury molecule 1 (*Havcr1*); *Lcn2*, lipocalin-2 (NGAL); *Ccl2*, chemokine (C-C motif) ligand 2 (MCP-1); *Tgfb1*, transforming growth factor, beta 1; *Col27a1*, collagen type XXVII, alpha 1; *Lyplal1*, lysophospholipase-like 1; *Cpb2*, carboxypeptidase B2; *Serpina3b*, serine (or cysteine) peptidase inhibitor, clade A, member 3B; *Ccl28*, chemokine (C-C motif) ligand 28; *Adamtsl2*, ADAMTS-like 2.

### Western blot

Protein extracts were obtained from frozen tissue and quantified using the bicinchoninic acid procedure. Equal amounts of protein were resolved on SDS-PAGE (10% Mini-PROTEAN® TGX Stain-Free™ Gels, Bio-Rad) and transferred to polyvinyl difluoride membranes (Bio-Rad). Western blots were performed as previously described [44]. Signals were acquired with a chemiluminescent detector (ChemiDoc™ Imaging System, Bio-Rad) and quantified using software provided by the manufacturer. Phosphorylation of GSK-3α/β at residue Ser21/Ser9 was analysed with a phospho-specific antibody (Cell Signaling, #9331). Signals were normalized to total expression of GSK-3β, obtained with rabbit monoclonal antibody (Cell Signaling #9315).

### Statistical analysis

Statistical analysis was performed with Prism 8 software (Graphpad, San Francisco, CA). Tests are indicated in each figure or table legend.

## RESULTS

### Transgenic expression of constitutively active SGK1 in the kidney increases phosphorylation of a downstream target

Total expression of SGK1 mRNA was analysed in renal cortex samples of wild type (WT) and Tg.sgk1 mice. We confirmed the expression of the transgene in kidney by RT-PCR using primers flanking the triple HA tag inserted in the COOH-terminus of Tg.sgk1 (Fig. 1A). As we have previously shown in other tissues[28, 29], Tg.sgk1 showed two bands corresponding to the endogenous SGK1 and the HA-labelled transgenic SGK1, respectively, while control animals showed only the lower band corresponding to the endogenous gene (Fig. 1B). The abundance of SGK1 transcript in renal cortex of control and Tg.sgk1 mice was examined by qPCR. Tg.sgk1 showed approximately a two-fold increased expression of SGK1, consistent with the use of a BAC containing the physiological SGK1 promoter (Fig.1C). Treatment with the mineralocorticoid deoxycorticosterone acetate (DOCA) in the context of high NaCl intake significantly increased SGK1 expression in transgenic, but not in WT animals (Fig. 1C). This suggests that constitutively active SGK1 may potentiate its own expression under certain conditions and is similar to what we have previously found in liver of animals treated with a high-fat diet [18]. In order to check whether transgenic expression of gain-of-function SGK1 mutant produces increased downstream signalling we tested the phosphorylation of GSK-3α/β at residues Ser21/Ser9, a well-known target of SGK1 [45]. Comparison between WT and Tg.sgk1 shows approximately a 50% increase in pGSK-3α/β abundance (Fig. 1D and 1E), in agreement with data obtained from other tissues [18].

**Figure 1.**
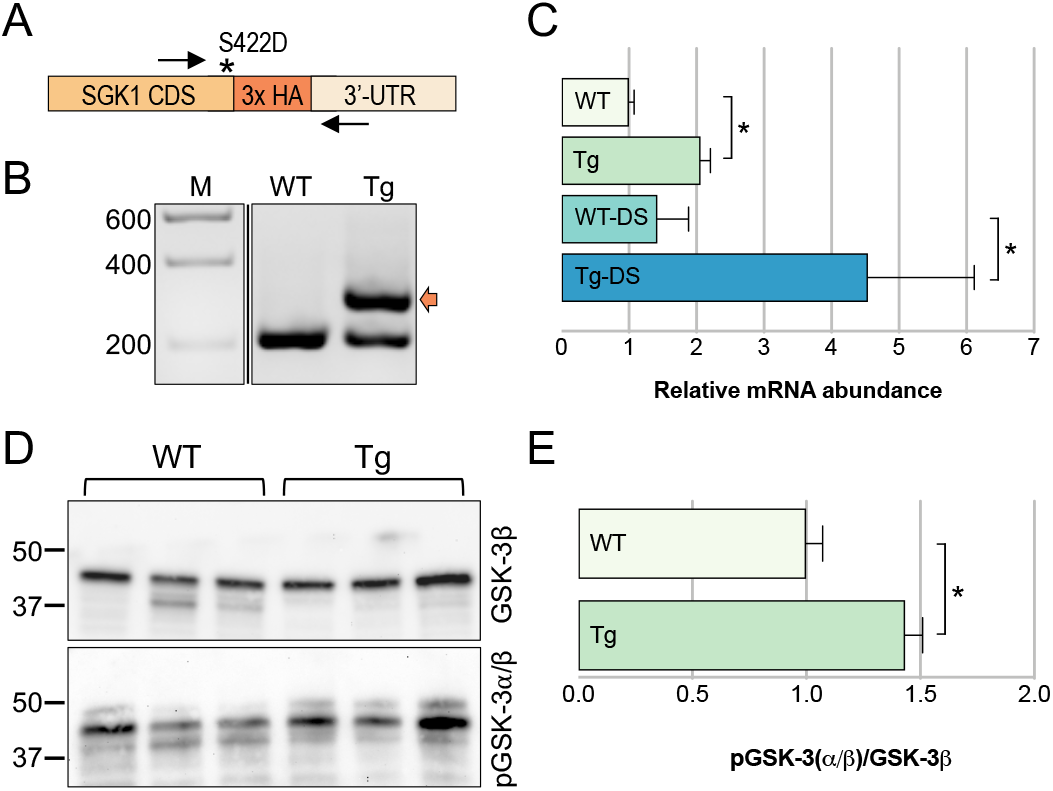
Transgenic expression of constitutively active SGK1 in the kidney and increased phosphorylation of SGK1 downstream target GSK-3. **A,** Schematic representation of the 3’ end of the modified *Sgk1* gene, including three copies of an HA tag (3x HA) and point mutation S422D, which renders the kinase constitutively active (asterisk). Arrows indicate oligonucleotides used to simultaneously detect endogenous and transgenic SGK1 by RT-PCR. **B**, Agarose gel electrophoresis analysis of RT-PCR products obtained from wild type (WT) or transgenic (Tg) kidney cortex cDNA using the oligonucleotide pair shown in panel A. M, molecular mass makers (values in base pairs). Arrow indicates the additional PCR product due to the presence of the transgene. **C,** Relative SGK1 mRNA abundance in the indicated WT and Tg.sgk1 mice experimental groups (DS, mice treated with DOCA/NaCl). Bars represent average ± SE (N=6-8) normalized to GAPDH expression. One-way ANOVA followed by Tukey’s multiple comparison test (*, p < 0.05). **D**, Representative western blots of kidney lysates from WT and Tg mice probed with anti-GSK-3β or anti-phospho-GSK-3α/β (pGSK-3α/β) antibodies. Lines indicate molecular mass marker migration (values in kDa). **E**, Relative abundance of pGSK-3α/β in WT and Tg.sgk1 kidney lysates. Bars represent average ± SE (N=6) pGSK3β/GSK3β normalized to WT. *, p < 0.05, unpaired t test.

### Blood pressure increase is more pronounced in uninephrectomized Tg.sgk1 mice treated with DOCA/NaCl

Two-kidney untreated Tg.sgk1 mice did not show any change in systemic blood pressure when compared to WT animals, even when animals where treated with DOCA/NaCl (Fig. 2A). Unilateral nephrectomy (NPX) did not alter blood pressure in Tg.sgk1 or WT mice when compared to two-kidney animals (Fig. 2A). Blood pressure increased after 4 weeks DOCA/NaCl treatment of NPX animals (NDS challenge), with a larger change in Tg.sgk1 mice (Fig. 2A). NDS lowered aldosterone circulating levels in WT animals, as expected, but no significant changes were detected in NDS Tg.sgk1 animals when compared to NPX (Table 2). There were no differences in water intake, diuresis and plasma or urine electrolytes between genotypes (Table 2).

**Figure 2.**
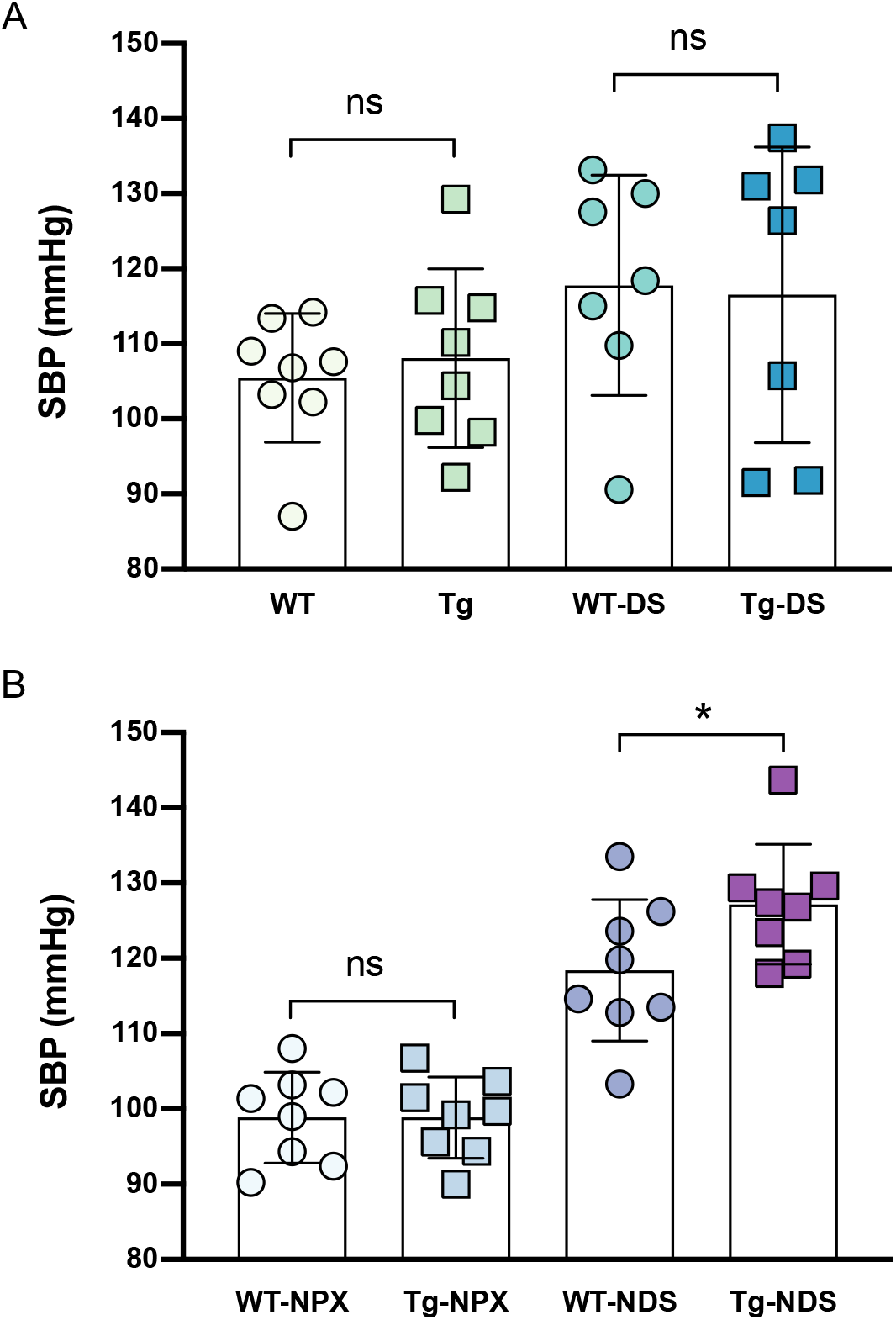
DOCA/NaCl-induced increase in systolic blood pressure is more pronounced in NPX Tg.sgk1 mice. **A,** Two-kidney animals left untreated or treated with DOCA/NaCl (DS) for 6 weeks. **B**, NPX animals left untreated or treated with DOCA/NaCl (NDS) for 6 weeks. Points represent average systolic blood pressure (SBP) in individual animals. Bars represent mean ± SD (N=6-8). One-way ANOVA followed by Sidak’s multiple comparison test for the indicated pairs of columns. *, p < 0.05; ns, no significant difference.

**Table 2.** General characteristics of nephrectomized (NPX) WT and Tg.sgk1 mice treated or not with DOCA/NaCl (NDS groups). Results are expressed as average ± SD (n, number of animals). Statistical significance was analysed using one-way ANOVA followed by Tukey’s multiple comparison test. There were no significant differences between genotypes in NPX or NDS groups. ^*^, p<0.05 compared to the same genotype.

| *Variable (units)* | NPX |  | NDS |  |
| --- | --- | --- | --- | --- |
|  | WT | Tg.sgk1 | WT | Tg.sgk1 |
| Kidney weight<sup>†</sup> | 7.5 ± 0.1; n=10 | 7.9 ± 0.2; n=10 | 9.8 ± 0.2*; n=12 | 10.0 ± 0.3*; n=10 |
| Aldosterone (nM) | 1.51 ± 0.59; n=4 | 0.77 ± 0.36; n=4 | 1.15 ± 0.21*; n=5 | 0.66 ± 0.25; n=5 |
| Corticosterone (nM) | 110 ± 17; n=4 | 55 ± 16; n=4 | 141 ± 52; n=5 | 85 ± 30; n=5 |
| Plasma Na<sup>+</sup> (mM) | 151.5 ± 1.9; n=5 | 150.0 ± 0.9; n=5 | 152.0 ± 1.3; n=6 | 150.8 ± 2.4; n=5 |
| Plasma K<sup>+</sup> (mM) | 7.4 ± 0.1; n=5 | 6.6 ± 0.3; n=5 | 6.8 ± 0.3; n=6 | 7.0 ± 0.4; n=5 |
| Plasma Cl<sup>-</sup> (mM) | 105.0 ± 1.1; n=5 | 102.3 ± 0.6; n=5 | 105.0 ± 0.8; n=6 | 102.8 ± 1.7; n=5 |
| Urine Na<sup>+</sup> (mM) | 71.2 ± 4.4; n=5 | 74.0 ± 13.3; n=5 | 273.0 ± 15.7*; n=6 | 260.0 ± 27.7*; n=5 |
| Urine K<sup>+</sup> (mM) | 235.1 ± 17.2; n=5 | 209.7 ± 30.7; n=5 | 27.8 ± 5.0*; n=6 | 33.0 ± 7.5*; n=5 |
| Urine Cl<sup>-</sup> (mM) | 160.6 ± 17.2; n=5 | 162.6 ± 26.7; n=5 | 292.3 ± 17.0*; n=6 | 276.3 ± 33.3*; n=5 |
| Diuresis (ml/24h) | 1.33 ± 0.22; n=5 | 1.76 ± 0.46; n=5 | 5.31 ± 0.22*; n=6 | 5.74 ± 0.19*; n=5 |
| Water intake (ml/24h) | 5.74 ± 0.51; n=5 | 8.42 ± 1.34; n=5 | 11.4 ± 0.56*; n=6 | 13.5 ± 2.01*; n=5 |

<sup>†</sup>Normalized to body weight x1000

### Excess SGK1 activity exacerbates albuminuria and induces hyperfiltration after nephrectomy and DOCA/NaCl treatment

Basal urinary albumin/creatinine ratio (uACR) was equal in WT and Tg.sgk1 and was enhanced by DOCA/NaCl treatment (Fig. 3A). Most importantly, uACR was significantly higher in Tg.sgk1 (Fig. 3A), suggesting damage in the glomerular filtration barrier. To evaluate the effect of SGK1 on glomerular function during the NDS challenge we measured GFR and urinary albumin excretion. Iohexol clearance measurements did not reveal any significant difference between untreated WT and Tg.sgk1 mice (Fig. 3B), with GFR values consistent with our previously reported results for male mice from the same strain [33]. NPX induced a decrease in GFR, as expected, with no significant differences between genotypes (Fig. 3B). When NPX animals were treated with DOCA/NaCl, GFR increased significantly only in Tg.sgk1, but not in WT animals (Fig. 3B).

**Figure 3.**
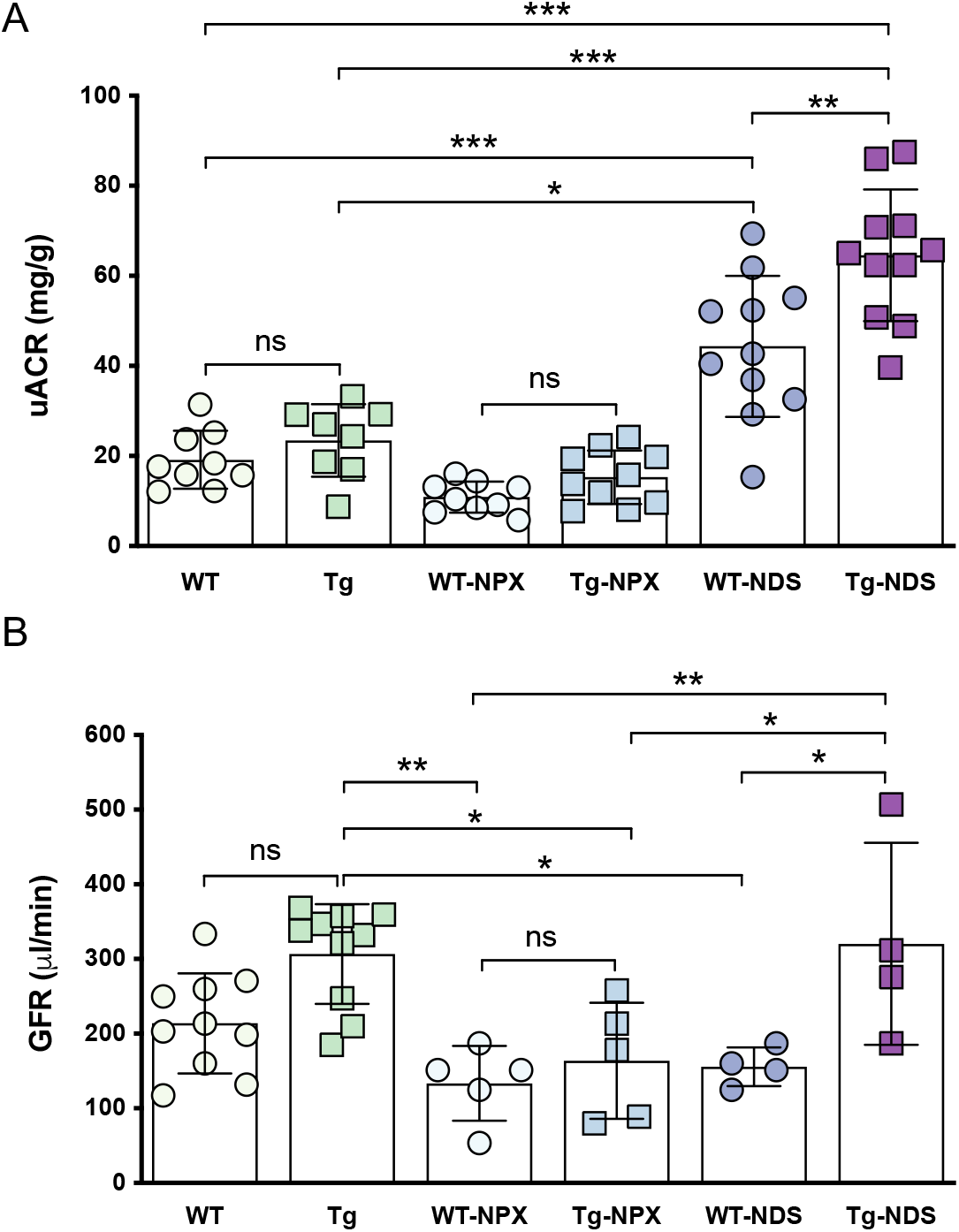
Increased SGK1 activity exacerbates albuminuria after NPX and DOCA/NaCl challenge. **A,** Urinary albumin/creatinine ratio (uACR). Data points represent measurements from individual mice. Bars represent experimental group average ± SD (N=8-12). Measurements were compared using one-way ANOVA followed by Tukey’s multiple comparison test (*, p<0.05; **, p<0.01; ***, p<0.001; ns, no significant difference). **B**, Glomerular filtration rate (GFR) was measured using the iohexol plasma clearance method in control wild type or transgenic animals (WT and Tg, N = 10), nephrectomized animals (WT-NPX and Tg-NPX, N = 5) and in animals after 5 weeks of NDS treatment (N = 4). Bars represent experimental group average ± SD. Measurements were compared using one-way ANOVA followed by Tukey’s multiple comparison test (*, p<0.05; **, p<0.01; ns, no significant difference).

### Tg.sgk1 display aggravated glomerular hypertrophy, fibrosis and inflammation under NDS treatment

NDS treatment significantly increased kidney weight in both WT and transgenic animals, without significant differences between genotypes (Table 2). Evaluation of excess SGK1 activity on NDS-induced changes in kidney morphology and fibrosis was analysed using Sirius Red staining (Fig. 4A and 4B). Upon NPX, Tg.sgk1 showed glomerular hypertrophy, an effect that was further potentiated when animals were treated with DOCA/NaCl (Fig. 4A). Quantification of Sirius Red-positive area in kidney cortex revealed increased glomerular and interstitial fibrosis, with a tendency towards increased perivascular fibrosis, in Tg.sgk1 mice when compared to WT mice after NDS (Fig. 4C). We next examined the expression of molecular markers of fibrosis in kidney cortex. SGK1 has previously been implicated in promoting the expression of connective tissue growth factor (CTGF) [46], a gene involved in the pathogenesis of fibrosis. DOCA/NaCl treatment significantly increased CTGF expression in transgenic animals, but not in their wild-type counterparts. There was no significant induction of TGF-β1 (Fig. 4D), consistently with the previously described aldosterone-induced pro-fibrotic pathway mediated by CTGF through a TGF-β1-independent pathway [47]. Fibronectin and collagen IV were also significantly more upregulated in transgenics, while α-SMA showed a tendency towards increased expression after DOCA/NaCl in both Tg.sgk1 and WT mice. Surprisingly, collagen I was less induced in transgenics than in WT (Fig. 4D). Expression of monocyte chemoattractant protein-1 (MCP-1, Ccl2), a marker of inflammation, was significantly upregulated by NDS challenge, although induction was slightly smaller in Tg.sgk1. mice (Fig. 4D).

**Figure 4.**
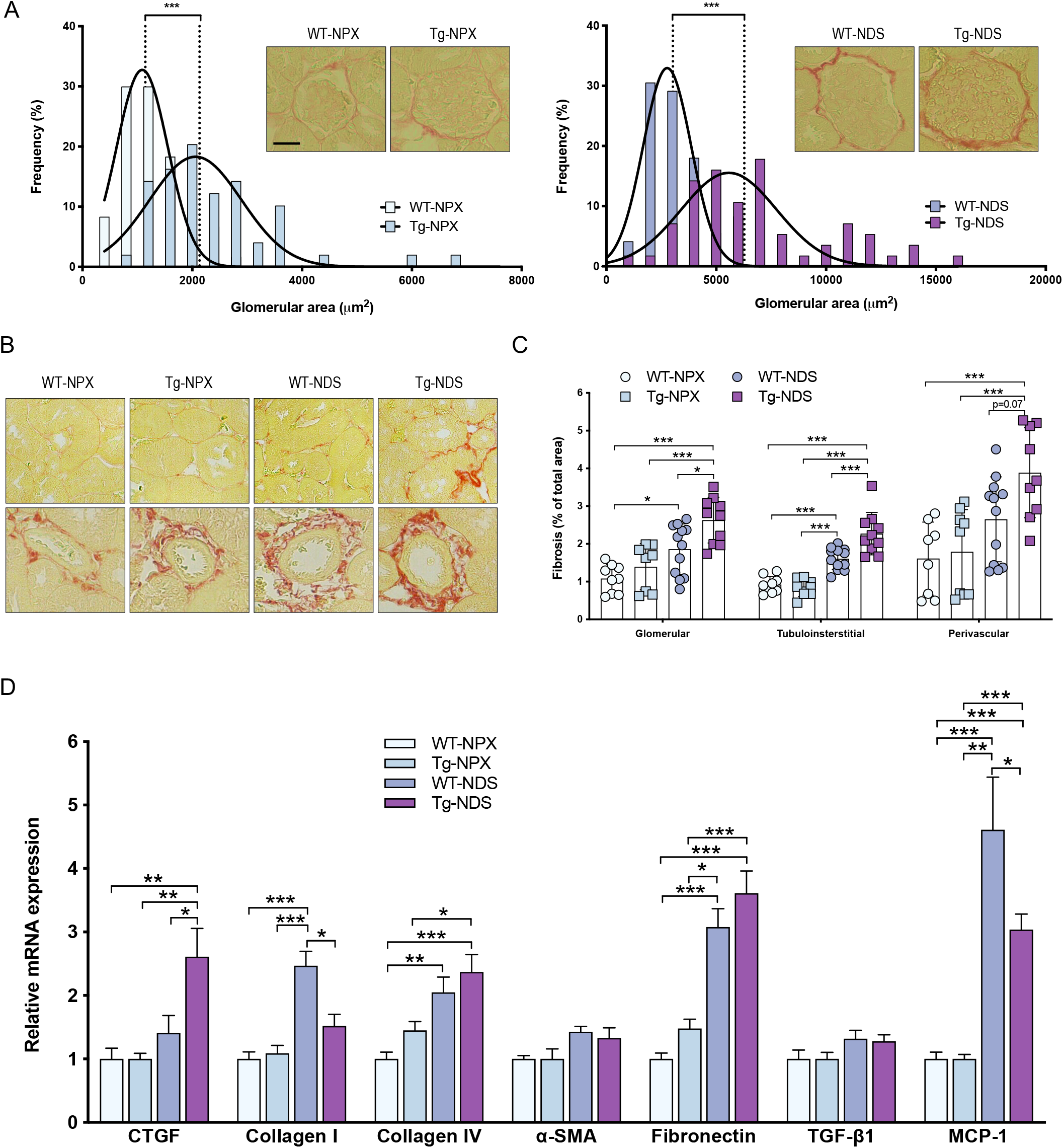
Tg.sgk1 display glomerular hypertrophy and increased fibrosis. **A,** Frequency distribution analysis of glomerular area in the indicated experimental groups. Insets show representative images of individual glomeruli from 3-μm tissue sections stained with Sirius red (scale bar = 30 μm). Plot bars represent percentage of glomeruli in each diameter class (n=46). Curves represent Gaussian fitting of the data. Medians of area distribution are indicated by dashed lines. Mann-Whitney test (***, p < 0.001). **B**, Representative images of kidney cortex obtained from the indicated experimental groups and stained with Sirius red. **C**, Quantification of Sirius red-positive area in glomeruli, interstitial and perivascular regions. Each point represents average percentage positive area obtained from five independent fields for one animal. Bars represent experimental group average ± SD (N= 8-12 animals). One-way ANOVA followed by Tukey’s multiple comparison test (*, p < 0.05; **, p < 0.01). **D**, Relative mRNA levels of the indicated fibrosis markers in WT and Tg.sgk1 mice experimental groups. Bars represent average ± SD (N=5-6) mRNA abundance values normalized to GAPDH expression. One-way ANOVA followed by Tukey’s multiple comparison test (*, p < 0.05; **, p < 0.01; ***, p < 0.001).

### Limited tubular injury in Tg.sgk1 mice

Given the increased tubulointerstitial fibrosis in Tg.sgk1 animals, we asked whether tubular function and/or markers of tubular injury are altered in transgenic mice. To functionally assess possible tubular injury, we measured fractional excretion of Na^+^ (FE_Na+_). FE_Na+_ in NPX mice was undistinguishable between WT and Tg.sgk1 (approx. 0.3%; Fig. 5A) and within the normal range [34, 48]. DOCA/NaCl increased FE_Na+_, as expected, but no significant difference was detected between WT and transgenic mice (Fig. 5A). NGAL (lipocalin 2) and KIM-1 are markers of tubular injury [49]. We detected significantly increased NGAL expression in DOCA/NaCl-treated transgenic animals (Fig. 5B). Taken together, these results suggest limited tubular injury in this model.

**Figure 5.**
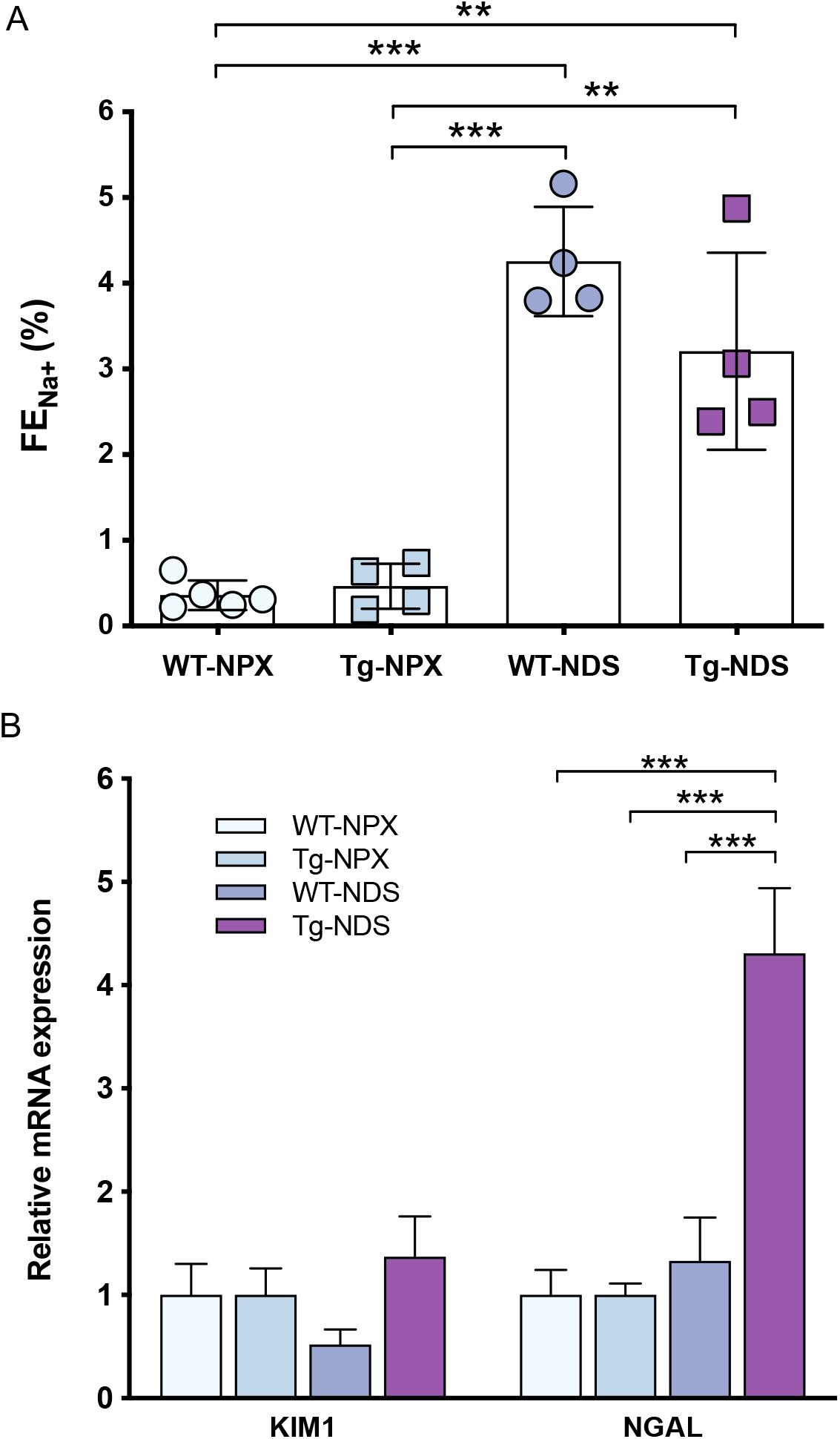
Limited tubular injury in NDS Tg.sgk1 mice. **A,** Fractional excretion of Na^+^ (FE_Na+_), expressed as %. Individual points represent values from each animal (N=4-5). Bars represent average ± SD. One-way ANOVA followed by Tukey’s multiple comparison test. **, p < 0.01; ***, p < 0.001. **B**, Relative expression of tubular injury markers in the indicated WT and Tg.sgk1 mice groups. Bars represent average ± SD (N=5-6) mRNA abundance values normalized to GAPDH expression. One-way ANOVA followed by Tukey’s multiple comparison test. *, p < 0.05; **, p < 0.01; ***, p < 0.001.

### Differentially regulated genes in Tg.sgk1 in response to DOCA/NaCl

To identify potential genes involved in the aggravated kidney injury observed in Tg.sgk1 following NDS we performed whole-transcriptome analysis using microarrays. Our analysis compared WT and Tg.sgk1 mice subject to the challenge (N=4 in each group). Differential expression (DE) analysis by limma resulted in identification of 198 probesets of which 120 were up-regulated and 78 were down-regulated at the given false discovery rate (FDR < 0.15; (on-line supplementary information, Table S1) based the volcano plot (Fig. 6A). Among these significantly modulated genes there were more up-regulated (n=38) than down-regulated (n=9) probesets exhibiting a log2 fold change greater than or equal to one (red probesets; Fig. 6A). *Cpb2* and *Serpind1a* were the most significantly modulated up-regulated and down-regulated genes, respectively (Supplementary Table S1). The hierarchical clustering of expression values of the genes obtained from the DE analysis demonstrated that there were two main clusters separating samples from WT and Tg.Sgk1 animals, respectively, as well as two clusters of genes exhibiting up-regulation and down-regulation of expression (Fig. 6B) with relatively stable within-cluster variability. The gene modules obtained for proteins with a physical- and/or context-driven interaction (String Database) between them were drawn by Cytoscape and helped identify the specific gene clusters/modules for the DE genes. There were two relatively large gene modules with 15 and 8 members, each, respectively. The 15-member gene network included the most significantly upregulated gene, *Cpb2*, along with several genes involved in collagen fibril formation, e.g., *Loxl, Col1a1* and *Col27a1* (Fig. 6C). On the other hand, 8-members gene network included mostly *Serpina* gene family in which *Serpina1* and *Serpina3* genes were down- and up-regulated, respectively (Fig. 6C). Two other smaller networks were also observed; one contained up-regulated members of cytoplasmic dynein family (*Dynlt1*) genes, known to be involved in vesicle transport, while the other gene module included down-regulated genes with relatively less well-known functions in the literature (Fig. 6C). Several other 3- or 2-member networks also emerged. We then performed a GO term enrichment analysis using ClusterProfiler package in R separately for up- and down-regulated genes, based on a less stringent FDR cut-off value (i.e., < 0.3; Supplementary Table S2). The resulting pathways for down-regulated genes included those involved in extracellular matrix (ECM) formation while up-regulated genes belonged to mostly immune response pathways, phagocytosis as well as lipid metabolism (Fig. 7; Supplementary Tables S3 and S4).

**Figure 6.**
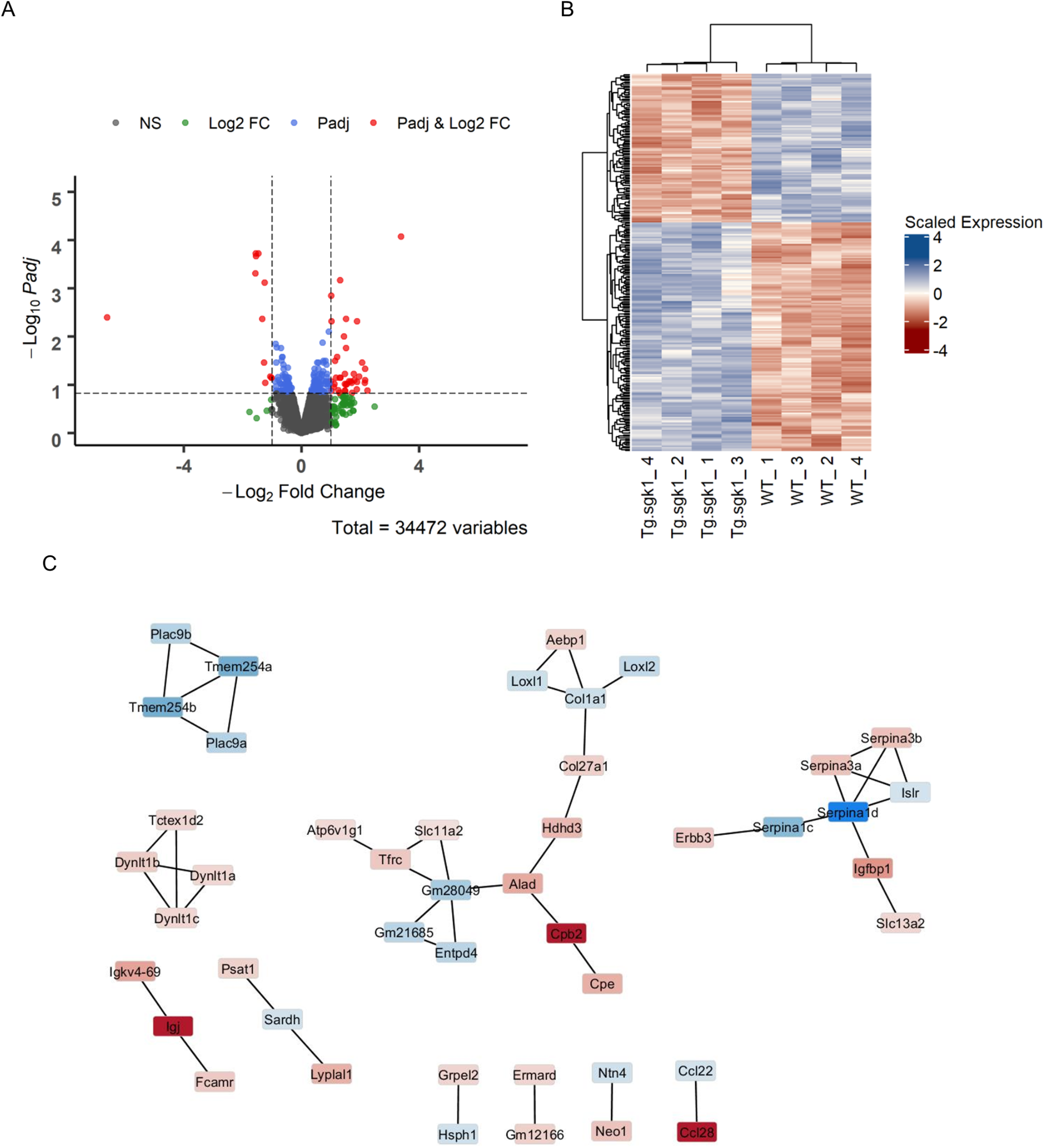
The differentially expressed genes obtained from the microarray experiment. **A,** Volcano plot showing the differentially upregulated and downregulated genes along with those not significantly changed. P stands for FDR<0.15 while the logarithmically (base 2) transformed fold change (log2FC) is greater than 1 or less than −1. Colours indicate genes that pass the thresholds given by only P, only log2FC or both. **B**, Heatmap of the differentially expressed genes with an adjusted p value threshold of FDR<0.15. Upregulation and downregulation are represented by a spectrum of red and blue, respectively. **C**, Gene network of differentially expressed genes between the groups. Nodes represent genes whose log2FC values are indicated by colours (pink-blue; up-down, respectively) while the edges that connect the nodes refer to the interactions specified by STRING database.

**Figure 7.**
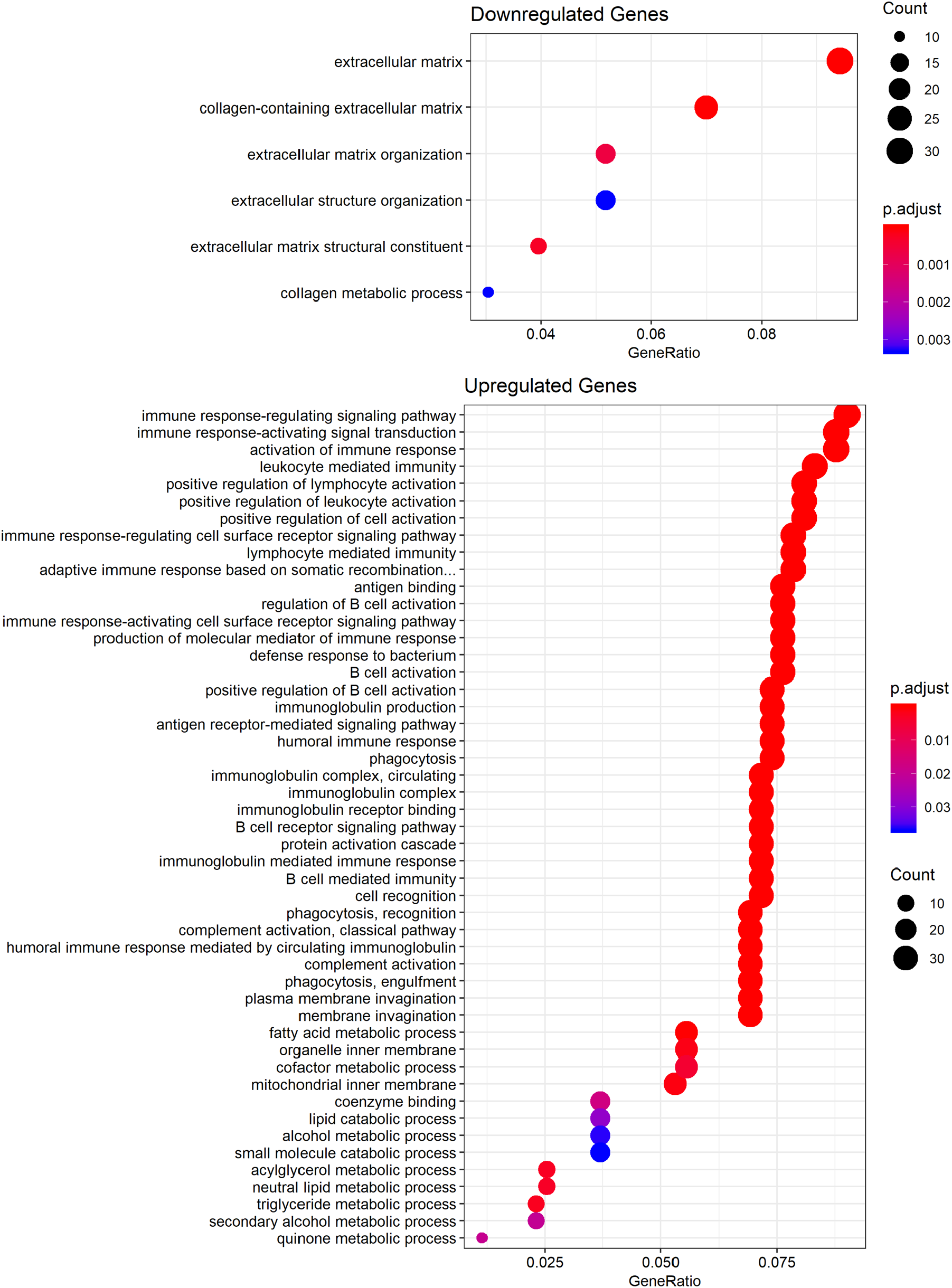
Functional annotation of differentially expressed genes in Tg.sgk1 after NDS. Gene Ontology (GO) term enrichment analysis for up- and down-regulated genes (FDR cut-off value < 0.3). Adjusted p values are indicated using the color scale shown in the figure. Circle diameter is proportional to the number of genes in each GO category.

Among differentially regulated genes detected by microarray analysis, we selected 5 up-regulated genes (*Cpb2, Lyplal1, Serpina3b, Ccl28 and Col27a1*) and 1 down-regulated gene (*Adamtsl2*) to confirm microarray data by quantitative PCR. To that end, we analysed a higher number of animals of the same cohort used to perform transcriptomics (N = 6) and expanded the analysis to the four groups of animals used throughout this study (NPX WT or Tg.sgk1, with or without DOCA/NaCl treatment). Our results were in accordance to the microarray analysis, showing that the expression of every gene tested was significantly different between WT-NDS and Tg-NDS (Fig. 8A). Interestingly, when the four experimental groups were examined in parallel, we detected three different patterns of regulation by SGK1 activity and DOCA/NaCl treatment. *Cpb2*, *Ccl28* and *Serpina3b* were upregulated by increased SGK1 activity, but DOCA/NaCl did not further increase their expression (Fig. 8A). This is also the case of the *Adamtsl2*, which was downregulated by increased SGK1 activity, but unaffected by DOCA/NaCl treatment (Fig. 8A). *Lyplal1* was also upregulated by increased SGK1 activity, but DOCA/NaCl additively increased its expression (Fig. 8A). In contrast, *Col27A1* was only upregulated by the synergism between increased SGK1 activity and DOCA/NaCl treatment (Fig. 8A). We then investigated whether these genes are direct targets of SGK1 signalling or, alternatively, whether their regulation is secondary to the deleterious effects induced by the transgene. To that end we obtained primary mouse embryonic fibroblasts (MEFs) from WT or transgenic animals (Fig. 8B). Of the six genes analysed, *Col27a1*, *Lyplal1* and *Ccl28* were detectable in both WT- and Tg.sgk1-derived MEFs (Fig. 8C). As expected from the results obtained from kidney samples, *Lyplal1* expression was significantly higher in MEFs derived from Tg.sgk1 (Fig. 8C). In contrast *Ccl28* expression was not significantly different between MEFs with the two different genotypes, suggesting that the upregulation detected *in vivo* may be secondary to other processes triggered by SGK1 in the kidney. *Col27A1* did not show any difference in expression, consistent with the results obtained from animals.

**Figure 8.**
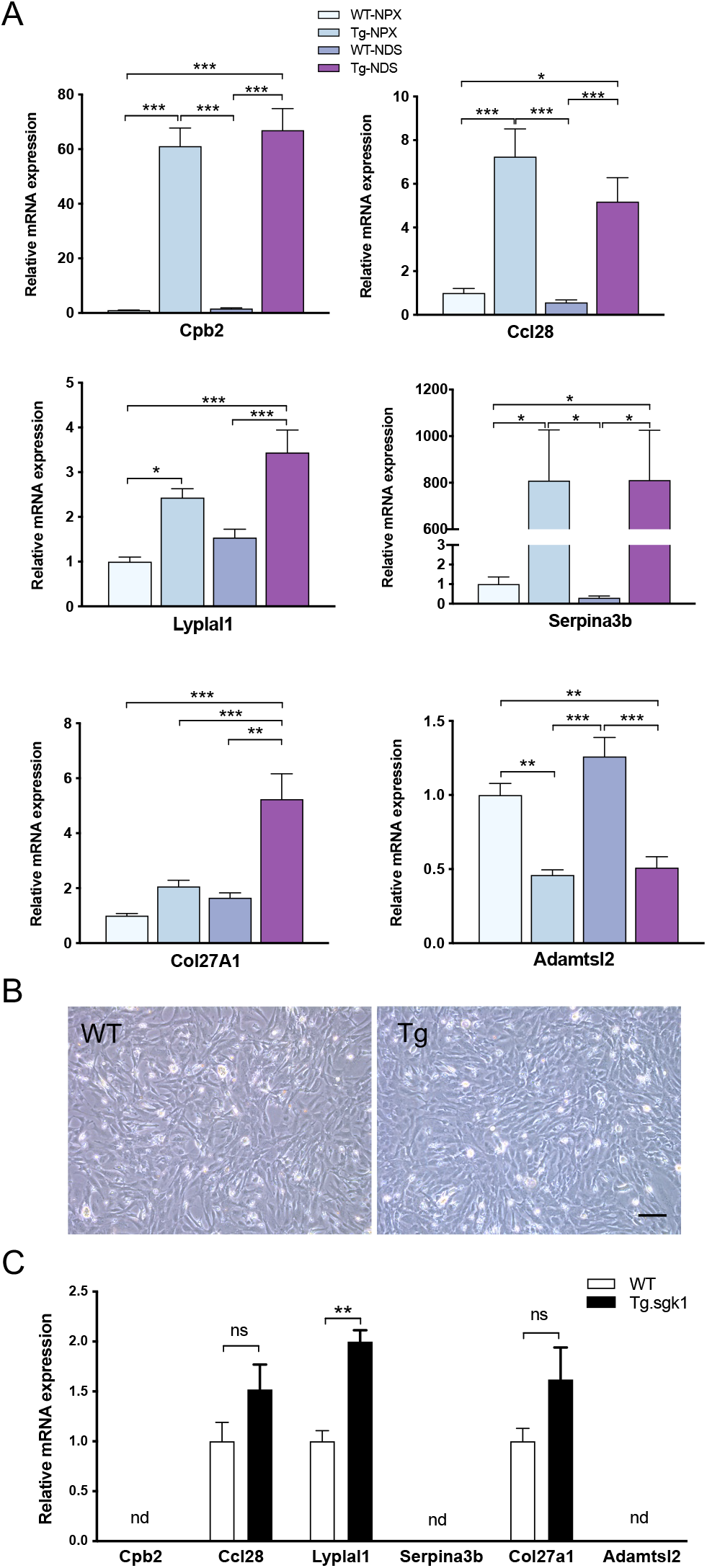
Validation of transcripts predicted by microarray analysis to be differentially regulated in NDS WT or Tg.sgk1 mice. **A,** Expression of the indicated genes was assessed by quantitative real-time PCR in the four groups used in this study (WT-NPX, Tg-NPX, WT-NDS and Tg-NDS). Bars represent average ± SE (N=6). Statistical analysis was performed by one-way ANOVA followed by Tukey’s Multiple Comparison Test (*, p<0.05; ** p<0.01; ***, p<0.001). **B**, Representative micrographs of cultured MEFs obtained from WT or Tg.sgk1 embryos. Scale bar, 200 μm. **C**, Expression of newly identified differentially expressed genes in MEFs from WT or transgenic animals. Bars represent average ± SE (N=6). Transcripts below qPCR detection level are indicated by “nd” (non-detected). Statistical analysis was performed by one-way ANOVA followed by Tukey’s Multiple Comparison Test (** p<0.01; ns, no significant difference).

## DISCUSSION

This study tested the hypothesis that increased systemic SGK1 activity constitutes a common factor underlying blood pressure-sensitive and insensitive renal damage secondary to excess mineralocorticoid signalling. To that end, we chose our previously characterized transgenic mouse model with expression of a constitutively active mutant of SGK1 expressed from a BAC [18, 28, 29]. This model has the advantage of expressing SGK1 under the control of its own promoter, avoiding overexpression of the kinase but at the same time enhancing downstream signalling. To produce mineralocorticoid-induced increases in blood pressure and development of kidney injury we used the DOCA/NaCl model, which in mice have to be performed in the context of low renal mass (uninephrectomy). Under these conditions, transgenic mice showed significantly increased blood pressure, albuminuria, hyperfiltration, glomerular hypertrophy and fibrosis when compared to their WT counterparts, consistent with accelerated kidney damage.

Our results support increased SGK1 activity as a risk factor for the development of mineralocorticoid-dependent kidney injury. In addition, the results shed light on the contribution of the kinase to hypertension-dependent and -independent pathways of renal damage. On one hand, our data showed that SGK1 enhances mineralocorticoid-induced increased blood pressure. This is consistent with previous data implicating SGK1 in the control of blood pressure in humans and mice. *SGK1* polymorphisms associate with increased blood pressure, hyperinsulinemia [12] and type 2 diabetes [16] in humans. *Sgk1* knockout mice maintain blood pressure at the expense of increased circulating aldosterone and this strain is unable to maximally activate Na^+^ reabsorption after a low Na^+^ challenge [8]. Lack of *Sgk1* precludes salt-induced hypertension in a high-fat diet context [10] and angiotensin II-induced hypertension [27]. Data from our studies expand these observations, showing that enhanced kinase activity exacerbates blood pressure increases in models with high-fat diet [18] and increased mineralocorticoid signalling.

A relevant finding of our study was that renal damage was only observed in the presence of a relevant reduction of renal mass (uninephrectomy). This indicates an interaction between a low nephron endowment and SGK1-mediated effects in the induction of renal damage in the context of hypertension. The incidence and prevalence of CKD do not match that of hypertension in humans. Recent studies indicated that not all subjects with hypertension are at risk for renal disease, only those with concomitant risk factors like metabolic syndrome or low renal mass [50]. In fact, a lower number of glomeruli has been associated with a higher risk of hypertension [51]. Also, living kidney donors, a healthy population, have an increased risk of hypertension after donation, compared to matched healthy controls that did not undergo a nephrectomy [52]. These data, together with our results may support the hypothesis that a reduction of renal mass in necessary to induce hypertension and that SGK1 may play a role in the pathogenesis of renal disease in this context.

On the other hand, our data also suggests that SGK1 may potentiate hypertension-independent kidney damage. It is important to note that blood pressure increase in the NDS model used in this study is modest when compared to the much larger effect that the kinase had in the context of a high-fat diet using the same transgenic mouse [18], with an average SBP of 170 mmHg after just 7 weeks of treatment (40% higher than WT animals). Analysis of these animal cohorts did not show differences in glomerular size or kidney fibrosis (Vastola-Mascolo, Velázquez-García and Alvarez de la Rosa, unpublished observations), indicating that during the time-frame used in this and our previously published studies (6-7 weeks) increased blood pressure alone is insufficient to produce kidney damage, at least in this genetic background. On the other hand, transgenic NPX animals (not treated with DOCA/NaCl) already demonstrated glomerular hypertrophy (Fig. 4A), even though there is no difference in SBP between both genotypes (Fig. 2B). Taken together, these results support a role for SGK1 in the early stages of hypertension-independent kidney damage. This is consistent with previously published observations showing that DOCA/NaCl-treated WT and *Sgk1*^−/−^ mice had similar blood pressures but only WT mice developed albuminuria and renal fibrosis [22].

The mechanisms involved in SGK1 contribution to development of glomerular hypertrophy and glomerular barrier damage are probably diverse. MR-dependent up-regulation of SGK1 in podocytes is linked to aldosterone-mediated oxidative stress and glomerular damage [24]. In addition, SGK1 increases ICAM-1 and CTGF expression in mesangial cells, promoting inflammatory and fibrotic responses [53]. Our results show increased CTGF expression without changes in TGF-β1 expression in Tg.sgk1, which is consistent with changes reported in aldosterone-induced fibrosis in a rat model of diabetic nephropathy [47]. Our results are also in agreement with the lack of TGF-β1 induction in the heart of DOCA/NaCl-treated mice [46] or rats [54] and the fact that CTGF can be induced independently of TGF-β1 in human mesangial cells [55]. In addition, *Sgk1* transcription is up-regulated by TGF-β1 [56] and therefore constitutive activation of the kinase appears to control downstream pro-fibrotic pathways. An additional convergence point between SGK1 and TGF-β signalling is provided by NEDD4-L, a ubiquitin ligase which modifies and targets for degradation Smad2/3, mediators of the canonical TGF-β-regulated transcriptional response. SGK1 phosphorylates NEDD4-L, preventing its interaction with Smad2/3 and therefore potentiating TGF-β signalling [57].

We can also speculate about additional pathways that may contribute to the harmful renal SGK1 effects. For instance, insulin-like growth factor-1 (IGF-1), which has been implicated in glomerular hypertrophy in the remnant kidney of NPX model [58], activates the PI3K pathway and SGK1 is one of the mediators of this pathway. Enhanced activity of MR in additional cell types, such as inflammatory cells, smooth muscle cells or the endothelium may cause damage to kidney structure and function. We have already mentioned that SGK1 expression on antigen-presenting cells contributes to renal inflammation [25]. In contrast, endothelial MR knockout prevents cardiac but not renal damage induced by DOCA/NaCl [59]. Therefore, SGK1-mediated effects detected in our study are likely independent of endothelial cell function.

In order to better understand the basic mechanism involved in SGK1-mediated kidney damage we compared whole transcriptome profiles of kidney cortex in WT and Tg.sgk1 mice in the NDS model. We detected 120 up-regulated and 78 down-regulated genes in transgenic mice (FDR < 0.15). To validate microarray results and assess whether our strategy was able to identify genes potentially involved in the exacerbation of DOCA/NaCl kidney injury we further studied five up-regulated and one down-regulated genes in an independent cohort of animals including the four experimental groups used throughout this study (NPX WT or Tg.sgk1, with or without DOCA/NaCl treatment). In every gene tested we were able to confirm the microarray analysis. Furthermore, we detected different regulation patterns by SGK1 and DOCA/NaCl treatment. Interestingly, *Col27a1* expression was not significantly affected by increased SGK1 activity in the kidney cortex or in cultured primary MEFs. However, the combination of increased SGK1 activity and DOCA/NaCl treatment produced a synergistic increase in *Col27a1*. *Col27a1*, a little known fibrillar collagen gene that plays a role during the calcification of cartilage and the transition of cartilage to bone [60, 61], has not been previously linked to kidney fibrosis. This result is highly significant, since it validates our strategy to identify genes potentially involved in the amplification of DOCA/NaCl kidney injury by SGK1 activity.

Analysis of gene modules formed by differentially expressed genes uncovered one cluster related to collagen fibril formation (including *Col27a1*, *Col1a1* and *Loxl*) and the fibrinolytic system (including *Cpb2*), which has also been implicated in the regulation of renal fibrosis [62]. GO term enrichment analysis showed an overall down-regulation of genes involved in ECM formation. While this last result may seem paradoxical, taken together the analysis strongly suggests that increased SGK1 leads to dysregulation of pathways controlling ECM remodelling [63]. This idea is supported by our analysis of classical fibrosis markers showing the expected overexpression of CTGF, fibronectin and collagen IV in Tg.sgk1 mice, but not significant changes in TGF-β1 or α-SMA, suggesting alternative pathways for fibrosis development [47]. Also, in agreement with this view are previous reports in the literature showing several downregulated ECM components associated with the development of fibrosis. For instance, decorin, one of the down-regulated genes in our gene list, is antifibrotic and decorin knockout mice show increased ECM accumulation in diabetic nephropathy [64]. Decorin sequesters TGF-β[65] and therefore its downregulation may enhance signalling through this pathway without alterations in TGF-βmRNA expression. Two membrane-bound matrix metalloproteinases (MMP14 and MMP15) also come up among the downregulated genes. There is strong evidence pointing to dysregulation of MMPs in a wide variety of kidney diseases, including acute kidney injury, diabetic nephropathy, glomerulonephritis [66]. MMP14 cleaves several ECM proteins, including collagen I, fibronectin and laminin [67]. Therefore, MMP14 has a degradative rather than synthetic role and its deficiency leads to fibrosis without altering collagen synthesis [68, 69]. In addition, both MMP14 and MMP15 activate the gelatinase MMP2, which together with MMP9 has extensive implications in kidney disease [67]. NGAL, which was found to be upregulated in our analysis, stabilizes MMP9 potentiating its function [70].

Some genes previously characterized as profibrotic showed lower expression in Tg.sgk1 when compared to WT mice. For instance, *Loxl1*, a member of the lysyl-oxidase-like family, crosslinks collagen and elastin and has been implicated in fibrogenesis [71]. Similarly, upregulation of *Loxl2* may lead to kidney fibrosis in *Col4a3* null mice [72]. On the other hand, there are reports showing that downregulation *Loxl2* may increase collagen formation via activation of TGF-β/Smad pathway in preeclamptic placenta [73]. This suggests that tissue specificity as well as genetic background may determine the effects of ECM regulators such as *Loxl2*. *Fbln5* (fibulin-5) [74] and *Serpinh1* [75] expression has been linked to development of kidney fibrosis as well. In addition to the hypothesis that these differentially expressed genes are further indication of dysregulated ECM turnover, it is conceivable that some of them may reflect negative feedback loops limiting the extent of fibrosis development. It is likely that both hypotheses are correct, depending on which individual gene or signalling pathway is being considered.

As mentioned above, the largest gene cluster obtained in our analysis included genes related not only to collagen fibril formation, but also to the fibrinolytic system, which participates in development of renal fibrosis [62]. Mice deficient in *Cpb2* (carboxypeptidase B2, also known as thrombin-activated fibrinolysis inhibitor -TAFI-), the most upregulated gene in the microarray, have better survival and less kidney injury in a sepsis model [76]. *Cpb2*/TAFI polymorphisms associate with CKD frequency in a human population [77], while its level show strong correlation with kidney function impairment. In addition, CPB2/TAFI inhibition ameliorates kidney fibrosis in an animal model of CKD [62]. Therefore, our results support a role for the fibrinolytic axis in exacerbated fibrosis observed in Tg.sgk1 mice.

The second largest cluster obtained by gene module analysis includes several members of the serine protease inhibitor clade A (*Serpina*) family. *Serpina1* (alpha-1-antitrypsin, AAT), the most downregulated gene in the microarray, has been shown to be the most abundant serine protease inhibitor with prominent anti-inflammatory and tissue-protective effects. *Serpin1a* knockout mice develop lung emphysema [78] and liver fibrosis [79]. In addition, AAT therapy protects against kidney injury secondary to ischemia-reperfusion [80]. On the other hand, serpina3, a well-known MR target [81] recently proposed as a renal fibrosis and inflammation marker during transition to CKD [82], is potently upregulated in Tg.sgk1 mice. Accordingly, our findings implicate abnormal accumulation and turnover of a network of ECM proteins in SGK1-driven fibrosis in early-stage CKD.

GO enrichment analysis for up-regulated genes uncovered mostly immune response pathways, phagocytosis and lipid metabolism. Modulation of immune response and phagocytosis pathways is in good agreement with the well-established role of SGK1 in salt-sensitive hypertension and renal injury through its expression in T cells and antigen-presenting cells [20, 25–27]. In fact, T cell-specific and dendritic cell-specific *Sgk1* knockout abrogates increased blood pressure, vascular and renal inflammation in response to DOCA/NaCl treatment [27] and the L-NAME high-salt protocol [25], respectively. *Ccl28* (C-C motif chemokine ligand 28), a prominently up-regulated gene in Tg.sgk1, has previously been characterized as lymphocyte homing factor in mucosal tissues [83] and was recently associated with the accumulation of B and T lymphocytes in the kidney during the transition from acute to chronic injury [84]. In addition, the list of differentially expressed genes includes several up-regulated genes coding for immunoglobulin chains. Taken together, our data not only supports the notion that SGK1 is a key component on T cell infiltration and inflammation in salt-sensitive kidney injury, but also supports a role for B cell infiltration in the process [85].

It is important to acknowledge several limitations in our study. First, for practical reasons we only performed experiments on male mice only. However, based on the known sexual dimorphism in blood pressure-dependent and -independent responses in mice [86, 87], additional experiments are needed to assess the role of SGK1 activity on female mice. Also, SGK1 effects on the DOCA/NaCl model are predictably the result of a synergy between pathways regulating renal ion transport, podocyte and mesangial cell function, ECM turnover and inflammatory responses. Dissecting the role of excess SGK1 activity in each of these components will require using tissue-specific, conditional SGK1 gain-of-function models.

In summary, our results support the idea that systemically increased SGK1 activity is a risk factor for the development of mineralocorticoid-dependent hypertension and incipient kidney damage, particularly in the context of low renal mass, potentially accelerating the progression to CKD. SGK1 activity alters pathways related to ECM turnover and immune system responses and may provide a therapeutic target to delay CKD development.

## CLINICAL PERSPECTIVES

1. *Background*. Excess aldosterone induces hypertension-dependent and -independent renal damage and progression to CKD. Protein kinase SGK1 is upregulated by aldosterone, has been linked to increased blood pressure and is pro-inflammatory and pro-fibrotic. Thus, increased SGK1 activity may represent a risk factor for accelerated renal damage.
2. *Summary of the results*. Increased SGK1 activity in the context of low renal mass and mineralocorticoid excess for salt status increases blood pressure and accelerates renal damage, with underlying alterations on extracellular matrix remodeling and immune response pathways.
3. *Potential significance of the results to human health and disease*. Increased SGK1 expression could be a marker for rapid progression towards CKD and a potential therapeutic target to slow down the process.

## ACKNOWLEDGEMENTS

We would like to thank Dr. Frederic Jaisser and Dr. Pablo Martín-Vasallo for critical reading of the manuscript, and Dr. Natalia Lopez-Andres and Dr. Eduardo Salido for expert advice on assessing kidney fibrosis. This study was funded by grant BFU2016-78374-R from Ministerio de Ciencia, Universidades e Innovación (MICINN, Spain) and European Union FP7 COST ADMIRE Network (BM1301). S.V.-G. is supported by Programa Agustin de Betancourt (Cabildo de Tenerife, Spain). E.P. is a researcher of the Ramón y Cajal Program (MICINN, RYC-2014-16573) and the Red de Investigación Renal (REDinREN, Spain, RD16/0009/0031).

## AUTHOR CONTRIBUTIONS

C.S.-R.: conceptualization, methodology, validation, formal analysis, investigation, writing, visualization.

S.V.-G.: methodology, validation, formal analysis, investigation, writing.

A.G.: validation, formal analysis, visualization.

A.V.-M.: validation, formal analysis, investigation.

A.E.R.-R.: methodology, investigation.

S.L.-L.: methodology, investigation, formal analysis.

G.H.: methodology, investigation, writing

J.F.N.-G.: conceptualization, formal analysis, writing.

E.P.: formal analysis, writing.

O.K.: validation, formal analysis, writing, visualization.

D.A.d.l.R.: conceptualization, formal analysis, writing, visualization, supervision, project administration, funding acquisition.

