## Supplemental Tables S1-S4 for "Increased SGK1 activity potentiates mineralocorticoid/NaCl-induced hypertension and kidney injury"

**SUPPLEMENTARY INFORMATION**

**Supplementary Table S1**. Microarray analysis of differentially regulated genes after NDS challenge (Tg.sgk1 *vs*. wild type mice). False discovery rate was set at <0.15.

| *Row.names* | *Gene.Symbol* | *logFC* | *P.Value* | *adj.P.Val* |
|---|---|---|---|---|
| 17301995 | Cpb2 | 3.382397 | 2.04E-09 | 8.42E-05 |
| 17304177 | Tmem254b; Tmem254c; Tmem254a | -1.55603 | 1.03E-08 | 0.000191 |
| 17304193 | Tmem254b; Tmem254c; Tmem254a | -1.4613 | 1.38E-08 | 0.000191 |
| 17304162 | Tmem254b; Tmem254c; Tmem254a | -1.54022 | 2.08E-08 | 0.000215 |
| 17308165 | Gm38463; Gm27177 | -1.56458 | 5.93E-08 | 0.00049 |
| 17340548 | Tmem181b-ps | 1.313552 | 9.87E-08 | 0.00068 |
| 17284981 | Akr1c21 | -1.25333 | 1.29E-07 | 0.00076 |
| 17296489 | Nnt | 1.016393 | 2.77E-07 | 0.001431 |
| 17283617 | Serpina1d | -6.6092 | 8.72E-07 | 0.004007 |
| 17330988 | Nxpe3 | -1.33576 | 1.11E-06 | 0.004326 |
| 17340435 | Pisd-ps2 | 1.515557 | 1.15E-06 | 0.004326 |
| 17426206 | Alad | 1.020603 | 1.47E-06 | 0.004837 |
| 17532694 | Pisd-ps3 | 1.893059 | 1.52E-06 | 0.004837 |
| 17230823 | Lyplal1 | 0.931049 | 2.67E-06 | 0.007883 |
| 17290155 | Gm7120 | 1.44696 | 3.62E-06 | 0.009991 |
| 17278261 | Serpina3b | 0.720256 | 5.18E-06 | 0.013393 |
| 17547837 | Gm27177; Gm21464 | -0.87928 | 5.81E-06 | 0.014129 |
| 17547846 | Gm27177 | -0.83275 | 7.35E-06 | 0.016878 |
| 17301647 | Entpd4; Gm21685 | -0.69684 | 8.29E-06 | 0.017324 |
| 17467441 | Igkv4-68 | 1.512723 | 8.38E-06 | 0.017324 |
| 17297660 | 1700112E06Rik | -0.65634 | 1.48E-05 | 0.026583 |
| 17368476 | Adamtsl2 | -0.64107 | 1.36E-05 | 0.026583 |
| 17467471 | Igkv4-50 | 1.209931 | 1.43E-05 | 0.026583 |
| 17301615 | Entpd4; Gm21685 | -0.67269 | 1.81E-05 | 0.031196 |
| 17233286 | Nepn | 0.7491 | 2.03E-05 | 0.031746 |
| 17320813 | Nell2; Gm30810 | 0.815681 | 1.93E-05 | 0.031746 |
| 17233347 | Gja1 | 0.579083 | 2.92E-05 | 0.034511 |
| 17301738 | Phyhip | -0.85746 | 2.63E-05 | 0.034511 |
| 17404534 | Lrrc31 | 0.749366 | 2.91E-05 | 0.034511 |
| 17455554 | N4bp2l1 | 0.540107 | 2.68E-05 | 0.034511 |
| 17467453 | Igkv4-59 | 2.061502 | 2.78E-05 | 0.034511 |
| 17301634 | Gm21451 | -0.61541 | 3.34E-05 | 0.038336 |
| 17426198 | Hdhd3 | 0.847678 | 3.44E-05 | 0.038342 |
| 17499796 | Gm15315 | -0.46681 | 4.16E-05 | 0.044126 |
| 17229257 | Gm23208 | -0.60724 | 4.91E-05 | 0.046932 |
| 17290163 | Ccl28 | 2.168179 | 5.39E-05 | 0.046932 |
| 17332915 | Dynlt1b | 0.573494 | 5.18E-05 | 0.046932 |
| 17340524 | Dynlt1a; Dynlt1c | 0.450981 | 5.27E-05 | 0.046932 |

Supplementary Table S1

| *Row.names* | *Gene.Symbol* | *logFC* | *P.Value* | *adj.P.Val* |
|---|---|---|---|---|
| 17220919 | A130010J15Rik | 0.738665 | 6.16E-05 | 0.052007 |
| 17519364 | Fam214a | 0.438079 | 6.89E-05 | 0.056938 |
| 17284548 | Ighv1-34 | 1.777692 | 7.69E-05 | 0.059065 |
| 17302289 | Pcdh17 | 0.869027 | 7.71E-05 | 0.059065 |
| 17246231 | Erbb3 | 0.632308 | 8.28E-05 | 0.059654 |
| 17467461 | Igkv4-57 | 1.472601 | 8.37E-05 | 0.059654 |
| 17527735 | Neo1 | 0.695312 | 8.30E-05 | 0.059654 |
| 17537901 | n-R5s12 | -0.55277 | 8.73E-05 | 0.060657 |
| 17274703 | Pxdn | 0.551222 | 9.40E-05 | 0.062684 |
| 17386697 | Gm13660 | 0.420535 | 9.32E-05 | 0.062684 |
| 17287486 | Cdhr2 | 0.687204 | 9.63E-05 | 0.063225 |
| 17231611 | Gm25682 | -1.05755 | 0.00011 | 0.067675 |
| 17459377 | Igkv6-25 | 1.962548 | 0.000108 | 0.067675 |
| 17506332 | Banp | -0.61382 | 0.000117 | 0.068966 |
| 17255258 | Gm22456 | -0.79529 | 0.000122 | 0.069104 |
| 17290010 | Esm1 | 0.440777 | 0.000125 | 0.069104 |
| 17458960 | Vmn1r20 | 0.859092 | 0.000119 | 0.069104 |
| 17478799 | Gm22494 | -0.50934 | 0.000125 | 0.069104 |
| 17236604 | Ntn4 | -0.46301 | 0.000128 | 0.069793 |
| 17247208 | Igfbp1 | 1.308051 | 0.000147 | 0.071458 |
| 17459423 | Igkv6-23; Igkv3-4; Igk-V8; Igk-V21; Igkc; Igk-V28 | 1.320944 | 0.000141 | 0.071458 |
| 17467421 | Igkv4-78 | 1.173084 | 0.000139 | 0.071458 |
| 17501584 | Gm15991 | -0.55789 | 0.000146 | 0.071458 |
| 17504122 | Ccl22 | -0.42412 | 0.000145 | 0.071458 |
| 17268010 | Acsf2 | 0.602345 | 0.000158 | 0.076116 |
| 17284530 | Ighv1-19 | 1.802558 | 0.000168 | 0.07964 |
| 17327557 | Mx2 | 0.464085 | 0.000172 | 0.080084 |
| 17347619 | Slc8a1 | 0.538424 | 0.000177 | 0.080084 |
| 17467537 | Igkv8-21 | 2.169239 | 0.000177 | 0.080084 |
| 17499686 | Gm21119 | 0.701668 | 0.000178 | 0.080084 |
| 17528249 | Slc51b | -0.79121 | 0.000171 | 0.080084 |
| 17442780 | Glt1d1 | 0.546347 | 0.000184 | 0.080804 |
| 17381622 | Gm23877 | -0.45648 | 0.000189 | 0.081195 |
| 17499637 | 4930467E23Rik; Gm15319 | 0.740597 | 0.000197 | 0.083892 |
| 17354784 | Mir143hg | -0.47427 | 0.000201 | 0.084728 |
| 17284354 | Igh-VX24; Ighm; Igh-VJ558; Ighv5-17; Ighv7-3; Ighj4; Ighg; Igh-V7183; Ighg3 | 1.620839 | 0.000221 | 0.085135 |
| 17313645 | Gm24575 | 0.839904 | 0.000229 | 0.085135 |
| 17408729 | Lrig2 | 0.457246 | 0.000213 | 0.085135 |
| 17444037 | Mafk | 0.421053 | 0.000226 | 0.085135 |
| 17459421 | Igk-V21; Igkj5 | 1.00501 | 0.000231 | 0.085135 |
| 17467435 | Igkv4-70 | 1.674561 | 0.000228 | 0.085135 |

Supplementary Table S1

| *Row.names* | *Gene.Symbol* | *logFC* | *P.Value* | *adj.P.Val* |
|---|---|---|---|---|
| 17467706 | Atoh8 | -0.51656 | 0.000225 | 0.085135 |
| 17482907 | Hs3st4 | -0.4861 | 0.000228 | 0.085135 |
| 17495258 | Gm23700 | 0.939275 | 0.000212 | 0.085135 |
| 17534213 | Rhox2c | -0.89376 | 0.00023 | 0.085135 |
| 17467389 | Igkv10-94 | 1.896121 | 0.000241 | 0.087368 |
| 17248957 | Olfr1392 | 0.595792 | 0.000274 | 0.090984 |
| 17283634 | Serpina1c | -1.23412 | 0.000274 | 0.090984 |
| 17284356 | Igh-V7183; Ighm; Igh-VJ558; Ighj3; Ighv4-2; Ighg; Ighv5-9 | 1.712063 | 0.000269 | 0.090984 |
| 17304186 | Plac9b; Plac9a; Gm9780 | -0.7294 | 0.00027 | 0.090984 |
| 17414672 | Col27a1 | 0.536162 | 0.000265 | 0.090984 |
| 17432264 | Plekhm2 | 0.474355 | 0.000266 | 0.090984 |
| 17449447 | Jchain | 2.15333 | 0.000275 | 0.090984 |
| 17487570 | Vmn1r127 | 0.528023 | 0.000272 | 0.090984 |
| 17266436 | Slc13a2 | 0.435515 | 0.000331 | 0.096093 |
| 17272461 | Cygb | -0.56091 | 0.000337 | 0.096093 |
| 17284314 | Igh-VJ558; Igha; Igh-VS107; Ighv7-1 | 1.480244 | 0.000327 | 0.096093 |
| 17286132 | Prl3d3 | -0.43628 | 0.000314 | 0.096093 |
| 17300163 | Traj44 | 0.921605 | 0.000328 | 0.096093 |
| 17304155 | Plac9b; Plac9a; Gm9780 | -0.69241 | 0.00034 | 0.096093 |
| 17304170 | Plac9b; Plac9a; Gm9780 | -0.69241 | 0.00034 | 0.096093 |
| 17383254 | Sardh | -0.43126 | 0.000342 | 0.096093 |
| 17455507 | Hsph1 | -0.45328 | 0.000309 | 0.096093 |
| 17467438 | Igkv4-69 | 1.119242 | 0.000324 | 0.096093 |
| 17467449 | Igkv4-62 | 1.549033 | 0.000295 | 0.096093 |
| 17472223 | Rerg | -0.45416 | 0.000326 | 0.096093 |
| 17509617 | Cpe | 0.952232 | 0.000324 | 0.096093 |
| 17509809 | Gm10033 | -0.41269 | 0.000298 | 0.096093 |
| 17509819 | Gm9495 | 0.769442 | 0.000327 | 0.096093 |
| 17509829 | Gm7697 | 0.837665 | 0.000301 | 0.096093 |
| 17263416 | Zfp867 | 0.563496 | 0.000356 | 0.098697 |
| 17331114 | Tbc1d23 | 0.64897 | 0.000358 | 0.098697 |
| 17354804 | Grpel2 | 0.478051 | 0.000363 | 0.098827 |
| 17467519 | Igkv8-30 | 1.517669 | 0.000363 | 0.098827 |
| 17292333 | Ptpdc1 | 0.406738 | 0.000388 | 0.103894 |
| 17463443 | Clec2h | 0.560727 | 0.000389 | 0.103894 |
| 17265672 | Spns2 | -0.47897 | 0.000395 | 0.104765 |
| 17254188 | Slfn3 | -0.52585 | 0.000398 | 0.104908 |
| 17212548 | Dnah7b | 0.575353 | 0.000411 | 0.10629 |
| 17432722 | Gm26880 | 0.903147 | 0.000416 | 0.106674 |
| 17526980 | Mir34b | -0.41066 | 0.000418 | 0.106674 |
| 17324835 | Tfrc | 0.626655 | 0.000424 | 0.107616 |

Supplementary Table S1

| *Row.names* | *Gene.Symbol* | *logFC* | *P.Value* | *adj.P.Val* |
|---|---|---|---|---|
| 17348435 | Cables1 | -0.63031 | 0.000429 | 0.108255 |
| 17331596 | Gm25038 | -0.86762 | 0.000437 | 0.108721 |
| 17467154 | Ppm1k | 0.543279 | 0.00044 | 0.108721 |
| 17520592 | Gm16010 | -0.44076 | 0.000434 | 0.108721 |
| 17315789 | Wdr70 | 0.429885 | 0.000465 | 0.111693 |
| 17459294 | Igkv1-133 | 1.107478 | 0.000463 | 0.111693 |
| 17424656 | Tpm2 | -0.36499 | 0.000473 | 0.111826 |
| 17366355 | Csprs | 0.518338 | 0.000489 | 0.1148 |
| 17357213 | Zbtb3 | -0.3329 | 0.000494 | 0.115333 |
| 17527661 | Islr | -0.40301 | 0.000497 | 0.115407 |
| 17273447 | Sectm1b | 0.623551 | 0.000531 | 0.121853 |
| 17241729 | Phyhipl | 0.546726 | 0.000539 | 0.122444 |
| 17400486 | Mtmr11 | 0.340881 | 0.000547 | 0.123672 |
| 17333966 | Zfp760 | 0.429807 | 0.000559 | 0.124819 |
| 17467466 | Igkv4-55 | 1.883184 | 0.000558 | 0.124819 |
| 17223616 | Raph1 | -0.38248 | 0.000572 | 0.126513 |
| 17396439 | Tnik | -0.41179 | 0.000576 | 0.126645 |
| 17284562 | Ighv1-43 | 1.252667 | 0.00059 | 0.129074 |
| 17322950 | Snx29 | 0.526864 | 0.000602 | 0.131015 |
| 17217639 | Lad1 | -0.49088 | 0.000625 | 0.131769 |
| 17230331 | Tfb2m | 0.603469 | 0.000616 | 0.131769 |
| 17230945 | Smyd2 | 0.577334 | 0.000619 | 0.131769 |
| 17284706 | Gm5441 | -0.42371 | 0.000623 | 0.131769 |
| 17467418 | Igkv4-79 | 0.963738 | 0.000631 | 0.131769 |
| 17491695 | Gm23962 | -0.72522 | 0.000612 | 0.131769 |
| 17501250 | Hpgd | -0.4911 | 0.00063 | 0.131769 |
| 17247241 | Abca13 | -0.8641 | 0.000669 | 0.131774 |
| 17262046 | Gm12166 | 0.530796 | 0.000664 | 0.131774 |
| 17278253 | Serpina3a | 0.717652 | 0.000646 | 0.131774 |
| 17284512 | Ighv1-9 | 2.240391 | 0.000674 | 0.131774 |
| 17311179 | Fzd6 | 0.484843 | 0.000645 | 0.131774 |
| 17395379 | Gm14412 | 0.56089 | 0.000653 | 0.131774 |
| 17406066 | Fnip2 | 0.465969 | 0.000679 | 0.131774 |
| 17411751 | Trp53inp1 | 0.369675 | 0.00065 | 0.131774 |
| 17459415 | Igkj1; Igk-V28 | 1.472622 | 0.000642 | 0.131774 |
| 17506882 | Gm22197 | -0.5992 | 0.000661 | 0.131774 |
| 17509069 | Cyp4v3 | 0.47808 | 0.000671 | 0.131774 |
| 17514961 | Olfr869 | -0.62137 | 0.000658 | 0.131774 |
| 17415432 | Gm22804 | -0.43022 | 0.00069 | 0.132672 |
| 17216949 | Fcamr | 0.614068 | 0.000724 | 0.133098 |
| 17284466 | Ighv14-4 | 1.484974 | 0.000711 | 0.133098 |

Supplementary Table S1

| *Row.names* | *Gene.Symbol* | *logFC* | *P.Value* | *adj.P.Val* |
|---|---|---|---|---|
| 17321683 | Slc11a2 | 0.342064 | 0.000716 | 0.133098 |
| 17459321 | Igkv9-120 | 1.019492 | 0.000714 | 0.133098 |
| 17527409 | Gm23407 | -0.5844 | 0.000722 | 0.133098 |
| 17527993 | LOC102638888 | -0.72984 | 0.000716 | 0.133098 |
| 17407116 | Cks1b | -0.37391 | 0.000749 | 0.135224 |
| 17437998 | Guf1 | 0.389128 | 0.000751 | 0.135224 |
| 17414760 | Atp6v1g1 | 0.390528 | 0.000764 | 0.135701 |
| 17531370 | Fbxw16 | -0.42768 | 0.000763 | 0.135701 |
| 17542711 | Gm6880; Gm15363 | -0.41089 | 0.000765 | 0.135701 |
| 17211335 | Tfap2d | -0.36908 | 0.000782 | 0.136928 |
| 17241954 | Gstt2 | 0.522335 | 0.000781 | 0.136928 |
| 17362397 | Gm24951 | -0.42811 | 0.000779 | 0.136928 |
| 17218073 | Pdc | -0.39586 | 0.00081 | 0.136958 |
| 17324794 | Tctex1d2 | 0.439465 | 0.000791 | 0.136958 |
| 17356739 | Mir194-2 | 0.593656 | 0.000791 | 0.136958 |
| 17363204 | Psat1 | 0.501733 | 0.0008 | 0.136958 |
| 17440342 | Gm15446 | -0.76563 | 0.000796 | 0.136958 |
| 17507860 | Gm15319 | 0.693711 | 0.000812 | 0.136958 |
| 17376063 | Zc3h6 | 0.772183 | 0.000822 | 0.138116 |
| 17255260 | Col1a1 | -0.4928 | 0.000839 | 0.13896 |
| 17461330 | Lrrn1 | -0.57435 | 0.00083 | 0.13896 |
| 17492175 | A830073O21Rik | -0.37745 | 0.000834 | 0.13896 |
| 17527694 | Loxl1 | -0.48711 | 0.00084 | 0.13896 |
| 17473155 | Cacng7 | -0.43167 | 0.000844 | 0.138976 |
| 17330947 | Zpld1 | -0.56702 | 0.000847 | 0.139031 |
| 17499650 | Gm15319 | 0.781234 | 0.000853 | 0.139446 |
| 17506923 | Slc35f3 | 0.61155 | 0.000869 | 0.14114 |
| 17535191 | Gm23000 | -0.82686 | 0.00087 | 0.14114 |
| 17325489 | Rabl3 | 0.42003 | 0.000876 | 0.1414 |
| 17247039 | Aebp1 | 0.507081 | 0.000891 | 0.14288 |
| 17407368 | Gm5849 | -0.46351 | 0.000892 | 0.14288 |
| 17383148 | Surf6 | 0.373019 | 0.000901 | 0.143905 |
| 17284379 | Igh-VJ558; Ighv5-4 | 1.718385 | 0.000926 | 0.146732 |
| 17308772 | Lacc1 | -0.66061 | 0.00093 | 0.146732 |
| 17333505 | Gm3435 | 0.462707 | 0.000928 | 0.146732 |
| 17459383 | Igkv6-17 | 1.29936 | 0.000937 | 0.147341 |
| 17441396 | Vsig10 | 0.453674 | 0.000943 | 0.147716 |

**Supplementary Table S2**. Microarray analysis of differentially regulated genes after NDS challenge (Tg.sgk1 *vs*. wild type mice) with a false discovery rate set at <0.3.

| *Row.names* | *Gene.Symbol* | *logFC* | *P.Value* | *adj.P.Val* |
|---|---|---|---|---|
| 17301995 | Cpb2 | 3.382397 | 2.04E-09 | 8.42E-05 |
| 17304177 | Tmem254b; Tmem254c; Tmem254a | -1.55603 | 1.03E-08 | 0.000191 |
| 17304193 | Tmem254b; Tmem254c; Tmem254a | -1.4613 | 1.38E-08 | 0.000191 |
| 17304162 | Tmem254b; Tmem254c; Tmem254a | -1.54022 | 2.08E-08 | 0.000215 |
| 17308165 | Gm38463; Gm27177 | -1.56458 | 5.93E-08 | 0.00049 |
| 17340548 | Tmem181b-ps | 1.313552 | 9.87E-08 | 0.00068 |
| 17284981 | Akr1c21 | -1.25333 | 1.29E-07 | 0.00076 |
| 17296489 | Nnt | 1.016393 | 2.77E-07 | 0.001431 |
| 17283617 | Serpina1d | -6.6092 | 8.72E-07 | 0.004007 |
| 17330988 | Nxpe3 | -1.33576 | 1.11E-06 | 0.004326 |
| 17340435 | Pisd-ps2 | 1.515557 | 1.15E-06 | 0.004326 |
| 17426206 | Alad | 1.020603 | 1.47E-06 | 0.004837 |
| 17532694 | Pisd-ps3 | 1.893059 | 1.52E-06 | 0.004837 |
| 17230823 | Lyplal1 | 0.931049 | 2.67E-06 | 0.007883 |
| 17290155 | Gm7120 | 1.44696 | 3.62E-06 | 0.009991 |
| 17278261 | Serpina3b | 0.720256 | 5.18E-06 | 0.013393 |
| 17547837 | Gm27177; Gm21464 | -0.87928 | 5.81E-06 | 0.014129 |
| 17547846 | Gm27177 | -0.83275 | 7.35E-06 | 0.016878 |
| 17301647 | Entpd4; Gm21685 | -0.69684 | 8.29E-06 | 0.017324 |
| 17467441 | Igkv4-68 | 1.512723 | 8.38E-06 | 0.017324 |
| 17297660 | 1700112E06Rik | -0.65634 | 1.48E-05 | 0.026583 |
| 17368476 | Adamtsl2 | -0.64107 | 1.36E-05 | 0.026583 |
| 17467471 | Igkv4-50 | 1.209931 | 1.43E-05 | 0.026583 |
| 17301615 | Entpd4; Gm21685 | -0.67269 | 1.81E-05 | 0.031196 |
| 17233286 | Nepn | 0.7491 | 2.03E-05 | 0.031746 |
| 17320813 | Nell2; Gm30810 | 0.815681 | 1.93E-05 | 0.031746 |
| 17233347 | Gja1 | 0.579083 | 2.92E-05 | 0.034511 |
| 17301738 | Phyhip | -0.85746 | 2.63E-05 | 0.034511 |
| 17404534 | Lrrc31 | 0.749366 | 2.91E-05 | 0.034511 |
| 17455554 | N4bp2l1 | 0.540107 | 2.68E-05 | 0.034511 |
| 17467453 | Igkv4-59 | 2.061502 | 2.78E-05 | 0.034511 |
| 17301634 | Gm21451 | -0.61541 | 3.34E-05 | 0.038336 |
| 17426198 | Hdhd3 | 0.847678 | 3.44E-05 | 0.038342 |
| 17499796 | Gm15315 | -0.46681 | 4.16E-05 | 0.044126 |
| 17229257 | Gm23208 | -0.60724 | 4.91E-05 | 0.046932 |
| 17290163 | Ccl28 | 2.168179 | 5.39E-05 | 0.046932 |
| 17332915 | Dynlt1b | 0.573494 | 5.18E-05 | 0.046932 |
| 17340524 | Dynlt1a; Dynlt1c | 0.450981 | 5.27E-05 | 0.046932 |
| 17220919 | A130010J15Rik | 0.738665 | 6.16E-05 | 0.052007 |

Supplementary Table S2

| *Row.names* | *Gene.Symbol* | *logFC* | *P.Value* | *adj.P.Val* |
|---|---|---|---|---|
| 17519364 | Fam214a | 0.438079 | 6.89E-05 | 0.056938 |
| 17284548 | Ighv1-34 | 1.777692 | 7.69E-05 | 0.059065 |
| 17302289 | Pcdh17 | 0.869027 | 7.71E-05 | 0.059065 |
| 17246231 | Erbb3 | 0.632308 | 8.28E-05 | 0.059654 |
| 17467461 | Igkv4-57 | 1.472601 | 8.37E-05 | 0.059654 |
| 17527735 | Neo1 | 0.695312 | 8.30E-05 | 0.059654 |
| 17537901 | n-R5s12 | -0.55277 | 8.73E-05 | 0.060657 |
| 17274703 | Pxdn | 0.551222 | 9.40E-05 | 0.062684 |
| 17386697 | Gm13660 | 0.420535 | 9.32E-05 | 0.062684 |
| 17287486 | Cdhr2 | 0.687204 | 9.63E-05 | 0.063225 |
| 17231611 | Gm25682 | -1.05755 | 0.00011 | 0.067675 |
| 17459377 | Igkv6-25 | 1.962548 | 0.000108 | 0.067675 |
| 17506332 | Banp | -0.61382 | 0.000117 | 0.068966 |
| 17255258 | Gm22456 | -0.79529 | 0.000122 | 0.069104 |
| 17290010 | Esm1 | 0.440777 | 0.000125 | 0.069104 |
| 17458960 | Vmn1r20 | 0.859092 | 0.000119 | 0.069104 |
| 17478799 | Gm22494 | -0.50934 | 0.000125 | 0.069104 |
| 17236604 | Ntn4 | -0.46301 | 0.000128 | 0.069793 |
| 17247208 | Igfbp1 | 1.308051 | 0.000147 | 0.071458 |
| 17459423 | Igkv6-23; Igkv3-4; Igk-V8; Igk-V21; Igkc; Igk-V28 | 1.320944 | 0.000141 | 0.071458 |
| 17467421 | Igkv4-78 | 1.173084 | 0.000139 | 0.071458 |
| 17501584 | Gm15991 | -0.55789 | 0.000146 | 0.071458 |
| 17504122 | Ccl22 | -0.42412 | 0.000145 | 0.071458 |
| 17268010 | Acsf2 | 0.602345 | 0.000158 | 0.076116 |
| 17284530 | Ighv1-19 | 1.802558 | 0.000168 | 0.07964 |
| 17327557 | Mx2 | 0.464085 | 0.000172 | 0.080084 |
| 17347619 | Slc8a1 | 0.538424 | 0.000177 | 0.080084 |
| 17467537 | Igkv8-21 | 2.169239 | 0.000177 | 0.080084 |
| 17499686 | Gm21119 | 0.701668 | 0.000178 | 0.080084 |
| 17528249 | Slc51b | -0.79121 | 0.000171 | 0.080084 |
| 17442780 | Glt1d1 | 0.546347 | 0.000184 | 0.080804 |
| 17381622 | Gm23877 | -0.45648 | 0.000189 | 0.081195 |
| 17499637 | 4930467E23Rik; Gm15319 | 0.740597 | 0.000197 | 0.083892 |
| 17354784 | Mir143hg | -0.47427 | 0.000201 | 0.084728 |
| 17284354 | Igh-VX24; Ighm; Igh-VJ558; Ighv5-17; Ighv7-3; Ighj4; Ighg; Igh-V7183; Ighg3 | 1.620839 | 0.000221 | 0.085135 |
| 17313645 | Gm24575 | 0.839904 | 0.000229 | 0.085135 |
| 17408729 | Lrig2 | 0.457246 | 0.000213 | 0.085135 |
| 17444037 | Mafk | 0.421053 | 0.000226 | 0.085135 |
| 17459421 | Igk-V21; Igkj5 | 1.00501 | 0.000231 | 0.085135 |
| 17467435 | Igkv4-70 | 1.674561 | 0.000228 | 0.085135 |
| 17467706 | Atoh8 | -0.51656 | 0.000225 | 0.085135 |

Supplementary Table S2

| *Row.names* | *Gene.Symbol* | *logFC* | *P.Value* | *adj.P.Val* |
|---|---|---|---|---|
| 17482907 | Hs3st4 | -0.4861 | 0.000228 | 0.085135 |
| 17495258 | Gm23700 | 0.939275 | 0.000212 | 0.085135 |
| 17534213 | Rhox2c | -0.89376 | 0.00023 | 0.085135 |
| 17467389 | Igkv10-94 | 1.896121 | 0.000241 | 0.087368 |
| 17248957 | Olfr1392 | 0.595792 | 0.000274 | 0.090984 |
| 17283634 | Serpina1c | -1.23412 | 0.000274 | 0.090984 |
| 17284356 | Igh-V7183; Ighm; Igh-VJ558; Ighj3; Ighv4-2; Ighg; Ighv5-9 | 1.712063 | 0.000269 | 0.090984 |
| 17304186 | Plac9b; Plac9a; Gm9780 | -0.7294 | 0.00027 | 0.090984 |
| 17414672 | Col27a1 | 0.536162 | 0.000265 | 0.090984 |
| 17432264 | Plekhm2 | 0.474355 | 0.000266 | 0.090984 |
| 17449447 | Jchain | 2.15333 | 0.000275 | 0.090984 |
| 17487570 | Vmn1r127 | 0.528023 | 0.000272 | 0.090984 |
| 17266436 | Slc13a2 | 0.435515 | 0.000331 | 0.096093 |
| 17272461 | Cygb | -0.56091 | 0.000337 | 0.096093 |
| 17284314 | Igh-VJ558; Igha; Igh-VS107; Ighv7-1 | 1.480244 | 0.000327 | 0.096093 |
| 17286132 | Prl3d3 | -0.43628 | 0.000314 | 0.096093 |
| 17300163 | Traj44 | 0.921605 | 0.000328 | 0.096093 |
| 17304155 | Plac9b; Plac9a; Gm9780 | -0.69241 | 0.00034 | 0.096093 |
| 17304170 | Plac9b; Plac9a; Gm9780 | -0.69241 | 0.00034 | 0.096093 |
| 17383254 | Sardh | -0.43126 | 0.000342 | 0.096093 |
| 17455507 | Hsph1 | -0.45328 | 0.000309 | 0.096093 |
| 17467438 | Igkv4-69 | 1.119242 | 0.000324 | 0.096093 |
| 17467449 | Igkv4-62 | 1.549033 | 0.000295 | 0.096093 |
| 17472223 | Rerg | -0.45416 | 0.000326 | 0.096093 |
| 17509617 | Cpe | 0.952232 | 0.000324 | 0.096093 |
| 17509809 | Gm10033 | -0.41269 | 0.000298 | 0.096093 |
| 17509819 | Gm9495 | 0.769442 | 0.000327 | 0.096093 |
| 17509829 | Gm7697 | 0.837665 | 0.000301 | 0.096093 |
| 17263416 | Zfp867 | 0.563496 | 0.000356 | 0.098697 |
| 17331114 | Tbc1d23 | 0.64897 | 0.000358 | 0.098697 |
| 17354804 | Grpel2 | 0.478051 | 0.000363 | 0.098827 |
| 17467519 | Igkv8-30 | 1.517669 | 0.000363 | 0.098827 |
| 17292333 | Ptpdc1 | 0.406738 | 0.000388 | 0.103894 |
| 17463443 | Clec2h | 0.560727 | 0.000389 | 0.103894 |
| 17265672 | Spns2 | -0.47897 | 0.000395 | 0.104765 |
| 17254188 | Slfn3 | -0.52585 | 0.000398 | 0.104908 |
| 17212548 | Dnah7b | 0.575353 | 0.000411 | 0.10629 |
| 17432722 | Gm26880 | 0.903147 | 0.000416 | 0.106674 |
| 17526980 | Mir34b | -0.41066 | 0.000418 | 0.106674 |
| 17324835 | Tfrc | 0.626655 | 0.000424 | 0.107616 |
| 17348435 | Cables1 | -0.63031 | 0.000429 | 0.108255 |

Supplementary Table S2

| *Row.names* | *Gene.Symbol* | *logFC* | *P.Value* | *adj.P.Val* |
|---|---|---|---|---|
| 17331596 | Gm25038 | -0.86762 | 0.000437 | 0.108721 |
| 17467154 | Ppm1k | 0.543279 | 0.00044 | 0.108721 |
| 17520592 | Gm16010 | -0.44076 | 0.000434 | 0.108721 |
| 17315789 | Wdr70 | 0.429885 | 0.000465 | 0.111693 |
| 17459294 | Igkv1-133 | 1.107478 | 0.000463 | 0.111693 |
| 17424656 | Tpm2 | -0.36499 | 0.000473 | 0.111826 |
| 17366355 | Csprs | 0.518338 | 0.000489 | 0.1148 |
| 17357213 | Zbtb3 | -0.3329 | 0.000494 | 0.115333 |
| 17527661 | Islr | -0.40301 | 0.000497 | 0.115407 |
| 17273447 | Sectm1b | 0.623551 | 0.000531 | 0.121853 |
| 17241729 | Phyhipl | 0.546726 | 0.000539 | 0.122444 |
| 17400486 | Mtmr11 | 0.340881 | 0.000547 | 0.123672 |
| 17333966 | Zfp760 | 0.429807 | 0.000559 | 0.124819 |
| 17467466 | Igkv4-55 | 1.883184 | 0.000558 | 0.124819 |
| 17223616 | Raph1 | -0.38248 | 0.000572 | 0.126513 |
| 17396439 | Tnik | -0.41179 | 0.000576 | 0.126645 |
| 17284562 | Ighv1-43 | 1.252667 | 0.00059 | 0.129074 |
| 17322950 | Snx29 | 0.526864 | 0.000602 | 0.131015 |
| 17217639 | Lad1 | -0.49088 | 0.000625 | 0.131769 |
| 17230331 | Tfb2m | 0.603469 | 0.000616 | 0.131769 |
| 17230945 | Smyd2 | 0.577334 | 0.000619 | 0.131769 |
| 17284706 | Gm5441 | -0.42371 | 0.000623 | 0.131769 |
| 17467418 | Igkv4-79 | 0.963738 | 0.000631 | 0.131769 |
| 17491695 | Gm23962 | -0.72522 | 0.000612 | 0.131769 |
| 17501250 | Hpgd | -0.4911 | 0.00063 | 0.131769 |
| 17247241 | Abca13 | -0.8641 | 0.000669 | 0.131774 |
| 17262046 | Gm12166 | 0.530796 | 0.000664 | 0.131774 |
| 17278253 | Serpina3a | 0.717652 | 0.000646 | 0.131774 |
| 17284512 | Ighv1-9 | 2.240391 | 0.000674 | 0.131774 |
| 17311179 | Fzd6 | 0.484843 | 0.000645 | 0.131774 |
| 17395379 | Gm14412 | 0.56089 | 0.000653 | 0.131774 |
| 17406066 | Fnip2 | 0.465969 | 0.000679 | 0.131774 |
| 17411751 | Trp53inp1 | 0.369675 | 0.00065 | 0.131774 |
| 17459415 | Igkj1; Igk-V28 | 1.472622 | 0.000642 | 0.131774 |
| 17506882 | Gm22197 | -0.5992 | 0.000661 | 0.131774 |
| 17509069 | Cyp4v3 | 0.47808 | 0.000671 | 0.131774 |
| 17514961 | Olfr869 | -0.62137 | 0.000658 | 0.131774 |
| 17415432 | Gm22804 | -0.43022 | 0.00069 | 0.132672 |
| 17216949 | Fcamr | 0.614068 | 0.000724 | 0.133098 |
| 17284466 | Ighv14-4 | 1.484974 | 0.000711 | 0.133098 |
| 17321683 | Slc11a2 | 0.342064 | 0.000716 | 0.133098 |
| 17459321 | Igkv9-120 | 1.019492 | 0.000714 | 0.133098 |

Supplementary Table S2

| *Row.names* | *Gene.Symbol* | *logFC* | *P.Value* | *adj.P.Val* |
|---|---|---|---|---|
| 17527409 | Gm23407 | -0.5844 | 0.000722 | 0.133098 |
| 17527993 | LOC102638888 | -0.72984 | 0.000716 | 0.133098 |
| 17407116 | Cks1b | -0.37391 | 0.000749 | 0.135224 |
| 17437998 | Guf1 | 0.389128 | 0.000751 | 0.135224 |
| 17414760 | Atp6v1g1 | 0.390528 | 0.000764 | 0.135701 |
| 17531370 | Fbxw16 | -0.42768 | 0.000763 | 0.135701 |
| 17542711 | Gm6880; Gm15363 | -0.41089 | 0.000765 | 0.135701 |
| 17211335 | Tfap2d | -0.36908 | 0.000782 | 0.136928 |
| 17241954 | Gstt2 | 0.522335 | 0.000781 | 0.136928 |
| 17362397 | Gm24951 | -0.42811 | 0.000779 | 0.136928 |
| 17218073 | Pdc | -0.39586 | 0.00081 | 0.136958 |
| 17324794 | Tctex1d2 | 0.439465 | 0.000791 | 0.136958 |
| 17356739 | Mir194-2 | 0.593656 | 0.000791 | 0.136958 |
| 17363204 | Psat1 | 0.501733 | 0.0008 | 0.136958 |
| 17440342 | Gm15446 | -0.76563 | 0.000796 | 0.136958 |
| 17507860 | Gm15319 | 0.693711 | 0.000812 | 0.136958 |
| 17376063 | Zc3h6 | 0.772183 | 0.000822 | 0.138116 |
| 17255260 | Col1a1 | -0.4928 | 0.000839 | 0.13896 |
| 17461330 | Lrrn1 | -0.57435 | 0.00083 | 0.13896 |
| 17492175 | A830073O21Rik | -0.37745 | 0.000834 | 0.13896 |
| 17527694 | Loxl1 | -0.48711 | 0.00084 | 0.13896 |
| 17473155 | Cacng7 | -0.43167 | 0.000844 | 0.138976 |
| 17330947 | Zpld1 | -0.56702 | 0.000847 | 0.139031 |
| 17499650 | Gm15319 | 0.781234 | 0.000853 | 0.139446 |
| 17506923 | Slc35f3 | 0.61155 | 0.000869 | 0.14114 |
| 17535191 | Gm23000 | -0.82686 | 0.00087 | 0.14114 |
| 17325489 | Rabl3 | 0.42003 | 0.000876 | 0.1414 |
| 17247039 | Aebp1 | 0.507081 | 0.000891 | 0.14288 |
| 17407368 | Gm5849 | -0.46351 | 0.000892 | 0.14288 |
| 17383148 | Surf6 | 0.373019 | 0.000901 | 0.143905 |
| 17284379 | Igh-VJ558; Ighv5-4 | 1.718385 | 0.000926 | 0.146732 |
| 17308772 | Lacc1 | -0.66061 | 0.00093 | 0.146732 |
| 17333505 | Gm3435 | 0.462707 | 0.000928 | 0.146732 |
| 17459383 | Igkv6-17 | 1.29936 | 0.000937 | 0.147341 |
| 17441396 | Vsig10 | 0.453674 | 0.000943 | 0.147716 |
| 17359156 | Slc35g1 | -0.64533 | 0.000966 | 0.150763 |
| 17312191 | Them6 | 0.385068 | 0.00098 | 0.151392 |
| 17547835 | Gm20236 | -0.8463 | 0.000985 | 0.151392 |
| 17547844 | Gm20236 | -0.8463 | 0.000985 | 0.151392 |
| 17473695 | Gm6909 | -0.81612 | 0.000999 | 0.152922 |
| 17386499 | Cir1 | 0.335657 | 0.001004 | 0.153167 |
| 17397936 | Mbnl1 | 0.353849 | 0.001013 | 0.153557 |

Supplementary Table S2

| *Row.names* | *Gene.Symbol* | *logFC* | *P.Value* | *adj.P.Val* |
|---|---|---|---|---|
| 17372197 | Osbpl6 | 0.298377 | 0.001036 | 0.156345 |
| 17218233 | Colgalt2 | 0.433498 | 0.001069 | 0.157301 |
| 17241598 | Gm7075 | -0.57498 | 0.001055 | 0.157301 |
| 17284405 | Ighv5-9-1 | 0.699553 | 0.001061 | 0.157301 |
| 17329527 | Mb21d2 | 0.561272 | 0.001063 | 0.157301 |
| 17350921 | F830016B08Rik | -0.31385 | 0.001062 | 0.157301 |
| 17540935 | Gm22364 | -0.66318 | 0.001067 | 0.157301 |
| 17524336 | Olfr850 | -0.30733 | 0.001091 | 0.159418 |
| 17237132 | Lin7a | 0.326248 | 0.001127 | 0.162941 |
| 17252308 | Gm23226 | -0.38045 | 0.001165 | 0.163022 |
| 17277837 | Ttc8 | -0.48867 | 0.00116 | 0.163022 |
| 17328935 | Gm26219 | -0.56051 | 0.001159 | 0.163022 |
| 17340953 | Acat3 | 0.53746 | 0.001167 | 0.163022 |
| 17364167 | Gm25611 | -0.36823 | 0.00115 | 0.163022 |
| 17407850 | Ecm1 | -0.47466 | 0.001143 | 0.163022 |
| 17459335 | Igkv14-111 | 0.747405 | 0.001138 | 0.163022 |
| 17511550 | Tox3 | -0.46319 | 0.001164 | 0.163022 |
| 17399162 | Gm25945 | 0.365273 | 0.001172 | 0.163221 |
| 17284527 | Ighv1-18 | 1.339006 | 0.001182 | 0.163941 |
| 17239714 | Tcf21 | -0.54088 | 0.001199 | 0.164498 |
| 17347279 | Gm22562 | -0.58424 | 0.001192 | 0.164498 |
| 17423207 | Mir3471-1 | -0.57021 | 0.001202 | 0.164498 |
| 17421215 | BC080695 | -0.44966 | 0.001215 | 0.165177 |
| 17312339 | Gm23027 | -0.32555 | 0.001237 | 0.167192 |
| 17349803 | Gm6756 | 0.300126 | 0.001257 | 0.168023 |
| 17458514 | Npy | -0.3644 | 0.001263 | 0.168023 |
| 17463930 | Pik3c2g | 0.712709 | 0.001268 | 0.168023 |
| 17472185 | Art4 | -0.32401 | 0.001264 | 0.168023 |
| 17472192 | Mgp | -0.60295 | 0.001252 | 0.168023 |
| 17255064 | Mmd | -0.30111 | 0.001283 | 0.168956 |
| 17278700 | Mir3070b | -0.55628 | 0.001282 | 0.168956 |
| 17325933 | LOC102641930 | 0.463445 | 0.001301 | 0.170246 |
| 17497076 | Chst15 | -0.41389 | 0.001301 | 0.170246 |
| 17332923 | Dynlt1c | 0.405262 | 0.001313 | 0.170747 |
| 17404269 | Pde7a | 0.373402 | 0.001335 | 0.171993 |
| 17534899 | Smim10l2a | 0.509866 | 0.001334 | 0.171993 |
| 17398430 | Ctso | 0.358068 | 0.001342 | 0.172321 |
| 17245686 | Mettl21b | 0.568026 | 0.001376 | 0.173091 |
| 17254895 | Mks1 | 0.300007 | 0.001391 | 0.173091 |
| 17277232 | Coq6 | 0.318309 | 0.001361 | 0.173091 |
| 17284507 | Ighv1-7 | 0.826651 | 0.001423 | 0.173091 |
| 17312127 | Ptp4a3 | -0.3115 | 0.001377 | 0.173091 |

Supplementary Table S2

| *Row.names* | *Gene.Symbol* | *logFC* | *P.Value* | *adj.P.Val* |
|---|---|---|---|---|
| 17314575 | Gm25668 | -0.8072 | 0.001413 | 0.173091 |
| 17322429 | Ppp1r1a | -0.53258 | 0.001405 | 0.173091 |
| 17323838 | Klhl24 | 0.33635 | 0.001379 | 0.173091 |
| 17344978 | 9130008F23Rik | -0.38661 | 0.001365 | 0.173091 |
| 17380659 | Cdh4 | -0.5243 | 0.001423 | 0.173091 |
| 17408336 | Hsd3b3 | 0.38083 | 0.001368 | 0.173091 |
| 17427389 | Cyp2j6 | 0.307367 | 0.0014 | 0.173091 |
| 17482230 | Acsm5 | 0.350258 | 0.001394 | 0.173091 |
| 17501669 | Gm15717 | 0.411812 | 0.001353 | 0.173091 |
| 17517344 | Gm22304 | -0.43428 | 0.001418 | 0.173091 |
| 17235077 | Cnn2 | -0.4639 | 0.001449 | 0.175179 |
| 17216960 | Pigr | 0.76774 | 0.001487 | 0.176876 |
| 17315686 | C7 | -0.79812 | 0.001489 | 0.176876 |
| 17464672 | Slc25a13 | 0.309578 | 0.001482 | 0.176876 |
| 17467430 | Igkv4-72 | 1.42626 | 0.001483 | 0.176876 |
| 17484925 | Cracr2b | -0.3141 | 0.001488 | 0.176876 |
| 17284605 | Ighv1-62-1 | 1.425399 | 0.0015 | 0.177374 |
| 17498502 | Ccnd1 | 0.345011 | 0.001502 | 0.177374 |
| 17493658 | Serpinh1 | -0.33893 | 0.001507 | 0.17752 |
| 17289568 | Trim23 | 0.508359 | 0.00153 | 0.179724 |
| 17308744 | Gm25517 | -0.62889 | 0.001536 | 0.179851 |
| 17231491 | Gm3213 | -0.32541 | 0.001554 | 0.180935 |
| 17408648 | Olfml3 | -0.3793 | 0.001569 | 0.182192 |
| 17284645 | Ighv1-75 | 0.840871 | 0.001583 | 0.183357 |
| 17231159 | A730013G03Rik | -0.35781 | 0.00161 | 0.184635 |
| 17488117 | Gm15883 | 0.526845 | 0.001606 | 0.184635 |
| 17284423 | Ighv5-17; Ighg; Igh-V7183 | 0.485221 | 0.001637 | 0.184889 |
| 17423461 | Gm26148 | -0.4079 | 0.001634 | 0.184889 |
| 17434290 | Gm9615 | -0.38989 | 0.001635 | 0.184889 |
| 17468665 | Rab43 | 0.43206 | 0.001636 | 0.184889 |
| 17500289 | Prosc | 0.270534 | 0.00163 | 0.184889 |
| 17502440 | Klf2 | -0.55954 | 0.001642 | 0.18496 |
| 17529842 | n-R5s87 | 0.462137 | 0.001649 | 0.185237 |
| 17335046 | Ggnbp1 | -0.30683 | 0.001657 | 0.185626 |
| 17223283 | Satb2 | -0.40106 | 0.001795 | 0.185676 |
| 17231114 | Mir3473c | -0.85976 | 0.001811 | 0.185676 |
| 17277794 | Gpr65 | -0.46354 | 0.001683 | 0.185676 |
| 17279198 | Apopt1 | 0.27111 | 0.001816 | 0.185676 |
| 17284432 | Ighv14-1; Ighg | 1.417332 | 0.00175 | 0.185676 |
| 17284554 | Ighv1-37 | 0.811878 | 0.001819 | 0.185676 |
| 17284560 | Ighv1-42 | 0.969353 | 0.001674 | 0.185676 |
| 17298631 | Vstm4 | -0.37946 | 0.001697 | 0.185676 |

Supplementary Table S2

| *Row.names* | *Gene.Symbol* | *logFC* | *P.Value* | *adj.P.Val* |
|---|---|---|---|---|
| 17309099 | Klf12 | 0.466031 | 0.001719 | 0.185676 |
| 17318428 | Oplah | 0.299925 | 0.00176 | 0.185676 |
| 17333521 | Ermard | 0.319099 | 0.001682 | 0.185676 |
| 17343170 | Tff2 | -0.45464 | 0.001749 | 0.185676 |
| 17385363 | Gm23315 | -0.71401 | 0.001669 | 0.185676 |
| 17415728 | Angptl3 | 0.467026 | 0.001749 | 0.185676 |
| 17432387 | Tmem51 | 0.302151 | 0.001708 | 0.185676 |
| 17441490 | Fbxo21 | 0.359312 | 0.001756 | 0.185676 |
| 17459338 | Igk-V1; Igkv1-110; Igk-V5 | 1.63494 | 0.001695 | 0.185676 |
| 17467427 | Igkv4-73 | 1.479266 | 0.001728 | 0.185676 |
| 17474621 | Ckm | -0.27728 | 0.001786 | 0.185676 |
| 17506947 | Gm22730 | -0.47093 | 0.001736 | 0.185676 |
| 17531834 | Fbxl2 | 0.393341 | 0.001778 | 0.185676 |
| 17534528 | Xpnpep2 | 0.461854 | 0.001809 | 0.185676 |
| 17459409 | Igkv3-2 | 0.531489 | 0.00183 | 0.18639 |
| 17276386 | Rhoj | -0.48698 | 0.001854 | 0.187933 |
| 17211303 | Gm25166 | -0.50784 | 0.001883 | 0.188048 |
| 17281312 | Slc25a21 | 0.58693 | 0.00188 | 0.188048 |
| 17335467 | Cdkn1a | -0.39534 | 0.001874 | 0.188048 |
| 17467451 | Igkv4-61; Igkv4-68 | 1.540773 | 0.001882 | 0.188048 |
| 17480044 | Prss23os | 0.44467 | 0.001883 | 0.188048 |
| 17545917 | Rnf138rt1 | -0.31877 | 0.001876 | 0.188048 |
| 17232235 | Ctgf | -0.50168 | 0.001922 | 0.190918 |
| 17265690 | Spns3 | -0.26334 | 0.00192 | 0.190918 |
| 17357920 | Olfr1474 | -0.38771 | 0.001926 | 0.190918 |
| 17215605 | Ackr3 | -0.43707 | 0.001992 | 0.191003 |
| 17220907 | Irf6; A130010J15Rik | 0.528786 | 0.001997 | 0.191003 |
| 17261122 | Papolg | 0.301328 | 0.001996 | 0.191003 |
| 17284551 | Ighv1-36 | 0.853068 | 0.001998 | 0.191003 |
| 17285680 | Hist1h3h | -0.39866 | 0.001962 | 0.191003 |
| 17301147 | Fam124a | -0.29352 | 0.001939 | 0.191003 |
| 17302305 | Tdrd3 | -0.42543 | 0.00195 | 0.191003 |
| 17334956 | Crebrf | 0.549812 | 0.001983 | 0.191003 |
| 17335998 | Cyp4f40 | -0.44933 | 0.001982 | 0.191003 |
| 17337122 | H2-Q8; H2-Q6 | -0.495 | 0.002027 | 0.191003 |
| 17348492 | Lama3 | 0.416259 | 0.002019 | 0.191003 |
| 17389111 | Gm24885 | -0.27412 | 0.002027 | 0.191003 |
| 17392968 | Defb19 | -0.37334 | 0.002018 | 0.191003 |
| 17406492 | Kirrel | -0.32463 | 0.002035 | 0.191003 |
| 17473292 | Lilra5 | -0.31763 | 0.002018 | 0.191003 |
| 17499538 | Gm25278 | -0.45218 | 0.002022 | 0.191003 |
| 17544149 | Hmgn5 | 0.340538 | 0.001963 | 0.191003 |

Supplementary Table S2

| *Row.names* | *Gene.Symbol* | *logFC* | *P.Value* | *adj.P.Val* |
|---|---|---|---|---|
| 17289271 | Zbed3 | -0.35544 | 0.002059 | 0.191736 |
| 17283380 | Fbln5 | -0.44674 | 0.002065 | 0.191842 |
| 17272941 | LOC102640451; Gm11767 | -0.39059 | 0.002086 | 0.193361 |
| 17220533 | Bpnt1 | -0.31402 | 0.002104 | 0.193602 |
| 17341963 | Slc9a3r2 | -0.28463 | 0.002129 | 0.193602 |
| 17395398 | C330013J21Rik | 0.712139 | 0.002127 | 0.193602 |
| 17459260 | E230016M11Rik | 0.412811 | 0.002119 | 0.193602 |
| 17490578 | Rcn3 | -0.3474 | 0.00213 | 0.193602 |
| 17547632 | Gm36385 | -0.2818 | 0.002122 | 0.193602 |
| 17348604 | Impact | 0.437694 | 0.002138 | 0.193841 |
| 17284662 | Ighv1-83 | 1.714853 | 0.002163 | 0.194872 |
| 17490328 | Myh14 | 0.3752 | 0.002158 | 0.194872 |
| 17509985 | Armc6 | -0.26296 | 0.002162 | 0.194872 |
| 17225693 | Mterf4 | 0.381321 | 0.002172 | 0.194919 |
| 17306975 | Ctsg | -0.34055 | 0.002173 | 0.194919 |
| 17499696 | Gm20946 | 0.817169 | 0.0022 | 0.196849 |
| 17393764 | Gm23134 | 0.821152 | 0.002213 | 0.197228 |
| 17244697 | Gm25342 | -0.50056 | 0.002257 | 0.198531 |
| 17284633 | Ighv1-72 | 1.144827 | 0.002243 | 0.198531 |
| 17352016 | Gm26014 | -0.92925 | 0.002255 | 0.198531 |
| 17366201 | LOC101055672 | 0.33603 | 0.002247 | 0.198531 |
| 17240621 | Aim1 | 0.345005 | 0.002272 | 0.198979 |
| 17342642 | Dusp1 | 0.593841 | 0.002292 | 0.200138 |
| 17370124 | Cutal | 0.326956 | 0.002295 | 0.200138 |
| 17450947 | 1010001B22Rik | -0.49553 | 0.002299 | 0.200138 |
| 17539110 | Smpx | -0.26277 | 0.002326 | 0.202055 |
| 17289287 | Gm25213 | -0.32067 | 0.002337 | 0.202604 |
| 17313647 | Bik | 0.299184 | 0.002372 | 0.203478 |
| 17407721 | Gabpb2 | 0.371383 | 0.002379 | 0.203478 |
| 17425606 | Gm12526 | 0.303053 | 0.002365 | 0.203478 |
| 17475182 | Zfp526 | 0.30597 | 0.002371 | 0.203478 |
| 17540378 | Maob | 0.366508 | 0.002382 | 0.203478 |
| 17328780 | Car15 | -0.50832 | 0.002394 | 0.204115 |
| 17211305 | Pi15 | -0.39229 | 0.002407 | 0.204714 |
| 17214823 | n-R5s213 | -1.02568 | 0.00242 | 0.204905 |
| 17390810 | Gatm | 0.329324 | 0.002436 | 0.205157 |
| 17489682 | Kctd15 | -0.28989 | 0.002435 | 0.205157 |
| 17241162 | Sgpl1 | 0.348609 | 0.002467 | 0.205664 |
| 17245272 | Nup107 | 0.406836 | 0.002461 | 0.205664 |
| 17288836 | Gm26088 | -0.53601 | 0.002494 | 0.205778 |
| 17439842 | Pkd2 | 0.300974 | 0.002482 | 0.205778 |
| 17400862 | Hmgcs2 | 0.729569 | 0.002499 | 0.205783 |

Supplementary Table S2

| *Row.names* | *Gene.Symbol* | *logFC* | *P.Value* | *adj.P.Val* |
|---|---|---|---|---|
| 17269615 | Kat2a | -0.31206 | 0.002511 | 0.206392 |
| 17246284 | Suox | 0.306604 | 0.002549 | 0.20668 |
| 17255947 | Mir5119 | 0.588502 | 0.002543 | 0.20668 |
| 17284442 | Ighv14-2 | 0.804723 | 0.002536 | 0.20668 |
| 17490628 | Flt3l; Rpl13a | -0.44937 | 0.002524 | 0.20668 |
| 17508904 | Gm26584 | -0.25473 | 0.002533 | 0.20668 |
| 17429800 | D830031N03Rik | 0.369321 | 0.002567 | 0.206712 |
| 17456934 | Mest | -0.28306 | 0.00257 | 0.206712 |
| 17483802 | Wdr11 | 0.261002 | 0.002579 | 0.206712 |
| 17248474 | Mir218-2 | -0.36689 | 0.002614 | 0.206972 |
| 17252495 | Gm26121 | 0.684248 | 0.002613 | 0.206972 |
| 17274401 | 6030426L16Rik | 0.353358 | 0.002622 | 0.206972 |
| 17298223 | Sfmbt1 | 0.356353 | 0.002657 | 0.206972 |
| 17312581 | Mfsd3 | -0.30225 | 0.002604 | 0.206972 |
| 17329093 | Cyp2ab1 | -0.32218 | 0.002634 | 0.206972 |
| 17485192 | LOC105243090; Gm7579 | -0.37274 | 0.002653 | 0.206972 |
| 17506206 | Crispld2 | -0.38315 | 0.002608 | 0.206972 |
| 17532472 | Cdcp1 | 0.303904 | 0.002639 | 0.206972 |
| 17396001 | Gm22074 | -0.38232 | 0.002709 | 0.209539 |
| 17539818 | Mir500 | -0.56066 | 0.002711 | 0.209539 |
| 17488148 | Mir3101; Rab4b | 0.530819 | 0.002717 | 0.209554 |
| 17328610 | 2610318N02Rik | -0.3433 | 0.002725 | 0.209791 |
| 17260474 | Igfbp3 | -0.32022 | 0.002745 | 0.209882 |
| 17432976 | Angptl7 | 0.644189 | 0.002764 | 0.209882 |
| 17535558 | Bgn | -0.60511 | 0.002737 | 0.209882 |
| 17272047 | Recql5 | 0.336392 | 0.002789 | 0.210354 |
| 17299830 | Trav13d-1 | -0.5573 | 0.002792 | 0.210354 |
| 17279131 | Tnfaip2 | -0.30275 | 0.002839 | 0.21246 |
| 17467525 | Igkv8-28 | 0.778823 | 0.002847 | 0.21246 |
| 17246860 | Nipsnap1 | 0.261241 | 0.002885 | 0.213496 |
| 17362883 | Gm336 | 0.39817 | 0.002894 | 0.213496 |
| 17465728 | Gm13861 | -0.28612 | 0.002902 | 0.213496 |
| 17485943 | Shisa7 | 0.303061 | 0.002885 | 0.213496 |
| 17511450 | Siah1a | 0.281784 | 0.002897 | 0.213496 |
| 17522110 | Pfkfb4 | 0.290869 | 0.002899 | 0.213496 |
| 17323432 | Hic2 | 0.311474 | 0.002951 | 0.214122 |
| 17402517 | Gm23279 | -0.55415 | 0.002965 | 0.214122 |
| 17406960 | Mir92b | -0.37139 | 0.002925 | 0.214122 |
| 17421390 | Gm13238 | 0.922443 | 0.002942 | 0.214122 |
| 17465805 | Slc13a4 | 0.303203 | 0.00297 | 0.214122 |
| 17501787 | Tm6sf2 | 0.325872 | 0.002963 | 0.214122 |
| 17512716 | Terf2 | 0.291747 | 0.002939 | 0.214122 |

Supplementary Table S2

| *Row.names* | *Gene.Symbol* | *logFC* | *P.Value* | *adj.P.Val* |
|---|---|---|---|---|
| 17215557 | Platr5 | -0.37178 | 0.003031 | 0.216201 |
| 17254835 | Mir142hg | -0.31026 | 0.00303 | 0.216201 |
| 17299575 | Ang; Rnase4 | 1.501677 | 0.003033 | 0.216201 |
| 17465108 | Fezf1 | -0.36099 | 0.00302 | 0.216201 |
| 17379139 | L3mbtl1 | -0.36333 | 0.003043 | 0.216204 |
| 17430861 | Sesn2 | 0.395287 | 0.00305 | 0.216309 |
| 17278820 | Mir494 | -0.38946 | 0.003078 | 0.216827 |
| 17315305 | Igfbp6 | -0.32887 | 0.003078 | 0.216827 |
| 17475342 | Tgfb1 | -0.38943 | 0.003078 | 0.216827 |
| 17442149 | P2rx4 | 0.301883 | 0.003093 | 0.217507 |
| 17317278 | Klhl38 | 0.346052 | 0.003108 | 0.217799 |
| 17438233 | Gsx2 | -0.26865 | 0.003151 | 0.219296 |
| 17498897 | Tnfsf13b | -0.34225 | 0.003145 | 0.219296 |
| 17265129 | Acadvl | 0.27695 | 0.003169 | 0.219496 |
| 17450830 | Slc26a1 | 0.313471 | 0.003161 | 0.219496 |
| 17461448 | Grm7 | -0.30739 | 0.003169 | 0.219496 |
| 17337276 | Ppp1r18 | -0.33969 | 0.003185 | 0.220167 |
| 17413078 | Gm13304; Gm10591; Gm21541; Ccl21b; Ccl21c | -0.46186 | 0.003195 | 0.220167 |
| 17434230 | Gm13304; Gm10591; Gm21541; Ccl21b; Ccl21c | -0.46186 | 0.003195 | 0.220167 |
| 17235694 | Tbxa2r | -0.35316 | 0.003216 | 0.220426 |
| 17330499 | Zdhhc23 | -0.50076 | 0.003211 | 0.220426 |
| 17496651 | Sephs2 | 0.372592 | 0.003229 | 0.220426 |
| 17343538 | 4921501E09Rik | -0.26745 | 0.003251 | 0.221065 |
| 17395039 | Gm22773 | 0.352392 | 0.003269 | 0.221962 |
| 17363497 | C330002G04Rik | 0.546585 | 0.003292 | 0.222598 |
| 17488883 | Gm6579 | -0.3442 | 0.003295 | 0.222598 |
| 17440692 | Hps4 | 0.389729 | 0.003304 | 0.222873 |
| 17378848 | Snhg11 | 0.501759 | 0.003335 | 0.224584 |
| 17302711 | Cldn10 | -0.34915 | 0.003354 | 0.225327 |
| 17247356 | Gm12002 | 0.247776 | 0.003373 | 0.225648 |
| 17467486 | Igkv12-44 | 1.446253 | 0.003371 | 0.225648 |
| 17502954 | Dnajb1 | -0.38453 | 0.003385 | 0.225713 |
| 17216120 | Gm9994 | -0.35294 | 0.003408 | 0.226901 |
| 17212301 | Tmem182 | -0.31775 | 0.003427 | 0.227784 |
| 17257906 | Map2k6 | 0.330104 | 0.003485 | 0.228012 |
| 17306325 | Olfr1507 | -0.28058 | 0.003507 | 0.228012 |
| 17322146 | Csad | 0.428951 | 0.0035 | 0.228012 |
| 17400162 | Selenbp1 | 0.283932 | 0.003504 | 0.228012 |
| 17451356 | Sgsm1 | -0.31388 | 0.00349 | 0.228012 |
| 17455801 | Col1a2 | -0.42793 | 0.003493 | 0.228012 |
| 17465750 | Wdr91 | 0.350369 | 0.003463 | 0.228012 |
| 17467458 | Igkv4-57-1 | 1.607892 | 0.003468 | 0.228012 |

Supplementary Table S2

| *Row.names* | *Gene.Symbol* | *logFC* | *P.Value* | *adj.P.Val* |
|---|---|---|---|---|
| 17503705 | Gm3134 | 0.326868 | 0.0035 | 0.228012 |
| 17331306 | Crybg3 | 0.429179 | 0.003545 | 0.230108 |
| 17360557 | Gm17197 | -0.25023 | 0.003563 | 0.230161 |
| 17404596 | Kcnmb3 | -0.28701 | 0.003557 | 0.230161 |
| 17294907 | Zfyve16 | 0.327734 | 0.003664 | 0.233721 |
| 17314373 | Muc19 | -0.43332 | 0.00368 | 0.233721 |
| 17323603 | Scarf2 | -0.33665 | 0.003677 | 0.233721 |
| 17358454 | C030016D13Rik | -0.29861 | 0.003667 | 0.233721 |
| 17385085 | Nmi | 0.357154 | 0.00368 | 0.233721 |
| 17449578 | Gm19619 | -0.24283 | 0.003653 | 0.233721 |
| 17462373 | Cecr2 | 0.446947 | 0.003654 | 0.233721 |
| 17463298 | Gm16303 | -0.46079 | 0.003636 | 0.233721 |
| 17518780 | Car12 | 0.330362 | 0.003642 | 0.233721 |
| 17417985 | Foxo6os | -0.31394 | 0.003686 | 0.233751 |
| 17529764 | Pls1 | 0.358073 | 0.0037 | 0.233923 |
| 17375981 | Gm14010 | -0.31215 | 0.003714 | 0.234061 |
| 17284420 | Ighv5-16; Igh-VX24 | 1.795186 | 0.003725 | 0.234411 |
| 17415428 | Gm24796 | -0.38355 | 0.003742 | 0.234443 |
| 17329864 | Muc20 | -0.29505 | 0.003772 | 0.234514 |
| 17355460 | Gm20544 | -0.3387 | 0.003764 | 0.234514 |
| 17366052 | Rab11fip2 | 0.386455 | 0.003771 | 0.234514 |
| 17426126 | Fkbp15 | 0.336622 | 0.003787 | 0.234618 |
| 17241719 | D630013N20Rik; Phyhipl | 0.45287 | 0.003805 | 0.235142 |
| 17372444 | Nup35 | 0.392814 | 0.003831 | 0.236426 |
| 17215208 | Chrng | -0.30357 | 0.003859 | 0.237756 |
| 17232998 | 9030612E09Rik | -0.29963 | 0.003879 | 0.238329 |
| 17434510 | Slc25a40 | 0.322373 | 0.003876 | 0.238329 |
| 17220921 | A130010J15Rik | 0.31055 | 0.003907 | 0.238881 |
| 17244362 | Snrpf | -0.34551 | 0.003923 | 0.238881 |
| 17244657 | Gm25522 | -0.48245 | 0.003935 | 0.238881 |
| 17264448 | Arhgef15 | -0.25673 | 0.003897 | 0.238881 |
| 17288354 | BC048507 | -0.61866 | 0.003982 | 0.238881 |
| 17335788 | Gm25447 | -0.35623 | 0.003992 | 0.238881 |
| 17392787 | Gm14151; Gm14147 | -0.34506 | 0.003983 | 0.238881 |
| 17420777 | Aldh4a1 | 0.354607 | 0.003909 | 0.238881 |
| 17434310 | Fam133b | 0.34357 | 0.003938 | 0.238881 |
| 17459324 | Igkv1-117 | 1.707757 | 0.003985 | 0.238881 |
| 17463611 | Prh1 | -0.29614 | 0.003968 | 0.238881 |
| 17524760 | Kank2 | -0.38593 | 0.00398 | 0.238881 |
| 17529496 | Snx14 | 0.313954 | 0.003992 | 0.238881 |
| 17534720 | Hprt | 0.302991 | 0.003945 | 0.238881 |
| 17227077 | Prelp | -0.5488 | 0.004008 | 0.239477 |

Supplementary Table S2

| *Row.names* | *Gene.Symbol* | *logFC* | *P.Value* | *adj.P.Val* |
|---|---|---|---|---|
| 17282743 | Tgfb3 | -0.34151 | 0.004021 | 0.239784 |
| 17249256 | Olfr51 | -0.33434 | 0.004044 | 0.240577 |
| 17305487 | Gm17654 | -0.33053 | 0.004075 | 0.242084 |
| 17343117 | Gm22327 | -0.7507 | 0.004121 | 0.243775 |
| 17389481 | Arhgap11a | 0.453085 | 0.004132 | 0.244053 |
| 17284617 | Ighv1-62; Ighv1-64 | 0.796045 | 0.004153 | 0.244765 |
| 17380757 | Gm6307 | -0.34993 | 0.004162 | 0.244765 |
| 17377283 | Xrn2 | 0.333987 | 0.004193 | 0.246243 |
| 17232763 | Ddo | 0.321837 | 0.004233 | 0.246353 |
| 17257893 | Gm11684 | -0.31635 | 0.004284 | 0.246353 |
| 17274766 | Fam110c | 0.350953 | 0.004276 | 0.246353 |
| 17284607 | Ighv1-62-2; Tnpo3 | 1.398988 | 0.004244 | 0.246353 |
| 17284631 | Ighv1-71; Tnpo3 | 1.398988 | 0.004244 | 0.246353 |
| 17385073 | Rbm43 | 0.271414 | 0.004302 | 0.246353 |
| 17405355 | Tm4sf1 | -0.25322 | 0.004221 | 0.246353 |
| 17419090 | 1700003M07Rik | -0.34319 | 0.00427 | 0.246353 |
| 17447595 | Stk32b | -0.52639 | 0.004276 | 0.246353 |
| 17467929 | Gm22254 | 0.453426 | 0.004274 | 0.246353 |
| 17481770 | Adm | 0.265959 | 0.004286 | 0.246353 |
| 17489420 | Fxyd1 | -0.52622 | 0.004279 | 0.246353 |
| 17498370 | Nadsyn1 | 0.349937 | 0.004267 | 0.246353 |
| 17508396 | Ddhd2 | 0.384986 | 0.004288 | 0.246353 |
| 17527917 | B930092H01Rik | -0.30799 | 0.004264 | 0.246353 |
| 17223429 | Als2cr12 | 0.502195 | 0.00433 | 0.246463 |
| 17263925 | A530017D24Rik | 0.405234 | 0.004371 | 0.246463 |
| 17284657 | Ighv1-81 | 1.501991 | 0.00434 | 0.246463 |
| 17329877 | Rubcn | -0.28559 | 0.004336 | 0.246463 |
| 17337441 | Trim26 | 0.322754 | 0.004347 | 0.246463 |
| 17394911 | Gm26883 | -0.32829 | 0.004367 | 0.246463 |
| 17412965 | Gm13304; Gm10591; Gm21541; Ccl21b; Ccl21c | -0.46279 | 0.004381 | 0.246463 |
| 17413025 | Gm13304; Gm10591; Gm21541; Ccl21b; Ccl21c | -0.46279 | 0.004381 | 0.246463 |
| 17413123 | Gm13304; Gm10591; Gm21541; Ccl21b; Ccl21c | -0.46279 | 0.004381 | 0.246463 |
| 17419812 | Rsrp1 | 0.387375 | 0.004343 | 0.246463 |
| 17434150 | Gm13304; Gm10591; Gm21541; Ccl21b; Ccl21c | -0.46279 | 0.004381 | 0.246463 |
| 17333844 | Gm5145 | -0.38514 | 0.004409 | 0.247649 |
| 17408507 | Gm10355 | -0.55944 | 0.00442 | 0.247773 |
| 17264792 | Atp1b2 | -0.34661 | 0.00449 | 0.247884 |
| 17284919 | Akr1c14 | 0.735936 | 0.004475 | 0.247884 |
| 17290384 | Gm22750 | -0.34938 | 0.004547 | 0.247884 |
| 17292174 | Atxn1 | 0.465557 | 0.004459 | 0.247884 |
| 17317900 | Gm10362 | -0.32333 | 0.004444 | 0.247884 |
| 17325410 | Golgb1 | 0.391341 | 0.004455 | 0.247884 |

Supplementary Table S2

| *Row.names* | *Gene.Symbol* | *logFC* | *P.Value* | *adj.P.Val* |
|---|---|---|---|---|
| 17338612 | Gm26547 | -0.38594 | 0.004581 | 0.247884 |
| 17351811 | Acaa2 | 0.313533 | 0.004572 | 0.247884 |
| 17375006 | Ganc; Capn3 | 0.311952 | 0.004552 | 0.247884 |
| 17388725 | Fjx1 | -0.52372 | 0.00454 | 0.247884 |
| 17457731 | Try4 | -0.30994 | 0.004437 | 0.247884 |
| 17488073 | Gm24865; Gm4613 | 0.473607 | 0.004515 | 0.247884 |
| 17506081 | Osgin1; Mlycd | -0.42523 | 0.004498 | 0.247884 |
| 17513008 | Gm24103 | 0.396181 | 0.004548 | 0.247884 |
| 17533952 | Gm2863 | -0.4036 | 0.004525 | 0.247884 |
| 17543742 | Gm26152 | -0.32221 | 0.004529 | 0.247884 |
| 17301943 | Gm10847 | -0.26799 | 0.004605 | 0.247975 |
| 17340389 | 4930480K15Rik | -0.27143 | 0.004611 | 0.247975 |
| 17411605 | Sdcbp | 0.236693 | 0.004606 | 0.247975 |
| 17502084 | Mir1969 | -0.34405 | 0.004619 | 0.24802 |
| 17485702 | Leng9 | -0.29804 | 0.004643 | 0.2487 |
| 17396383 | Tnfsf10 | 0.423421 | 0.004659 | 0.24886 |
| 17318734 | Zfp647 | -0.42982 | 0.004678 | 0.249547 |
| 17236800 | Dcn | -0.76908 | 0.004697 | 0.250199 |
| 17380222 | Pck1 | 0.350683 | 0.004708 | 0.250199 |
| 17433055 | Kif1b | 0.261409 | 0.004708 | 0.250199 |
| 17529901 | Slc25a36 | 0.337212 | 0.004754 | 0.251764 |
| 17363156 | Gm22323 | 0.685118 | 0.004783 | 0.252246 |
| 17363158 | Gm23931 | 0.685118 | 0.004783 | 0.252246 |
| 17338660 | Shd | -0.42743 | 0.004804 | 0.252762 |
| 17425703 | Gm12580 | -0.22631 | 0.004833 | 0.252762 |
| 17432988 | Tardbp | 0.249239 | 0.004824 | 0.252762 |
| 17316276 | Gm26163 | -0.43236 | 0.004874 | 0.254102 |
| 17494111 | Olfr551 | -0.26072 | 0.004888 | 0.254512 |
| 17436438 | Uvssa | 0.445717 | 0.004897 | 0.254692 |
| 17388389 | Cry2 | 0.394595 | 0.004926 | 0.254885 |
| 17476317 | Clip3 | -0.33355 | 0.004951 | 0.255549 |
| 17463829 | Atf7ip | 0.42251 | 0.004958 | 0.255611 |
| 17296528 | 3110070M22Rik | -0.30751 | 0.005016 | 0.256362 |
| 17321918 | Gm5476 | -0.41187 | 0.005 | 0.256362 |
| 17393092 | Dnmt3bos | -0.29579 | 0.005008 | 0.256362 |
| 17527848 | Thsd4 | -0.38321 | 0.005012 | 0.256362 |
| 17250855 | 9630013K17Rik | -0.28566 | 0.005029 | 0.25669 |
| 17221079 | Gm25493 | 0.303584 | 0.005065 | 0.257867 |
| 17223852 | D630023F18Rik | 0.337376 | 0.005068 | 0.257867 |
| 17512975 | Ddx19a | -0.34214 | 0.005077 | 0.257867 |
| 17464901 | Tmem168 | 0.293932 | 0.005085 | 0.257982 |
| 17214368 | Cyp27a1 | 0.392794 | 0.005188 | 0.25808 |

Supplementary Table S2

| *Row.names* | *Gene.Symbol* | *logFC* | *P.Value* | *adj.P.Val* |
|---|---|---|---|---|
| 17218861 | F5 | 0.556077 | 0.005132 | 0.25808 |
| 17284648 | Ighv1-76 | 1.66375 | 0.005156 | 0.25808 |
| 17302102 | Ccdc122 | -0.39957 | 0.005169 | 0.25808 |
| 17306877 | Adcy4 | -0.36828 | 0.005169 | 0.25808 |
| 17310430 | Mir1898 | 0.378457 | 0.0052 | 0.25808 |
| 17363470 | Gda | 0.318907 | 0.00513 | 0.25808 |
| 17461080 | Gm22840 | -0.30499 | 0.005164 | 0.25808 |
| 17473817 | Zfp446 | 0.280848 | 0.005163 | 0.25808 |
| 17481248 | Olfr632 | -0.45964 | 0.00516 | 0.25808 |
| 17497044 | Ikzf5 | 0.337208 | 0.005116 | 0.25808 |
| 17498084 | Krtap5-1 | -0.45929 | 0.005195 | 0.25808 |
| 17503681 | Cyld | 0.30163 | 0.005102 | 0.25808 |
| 17507872 | 2610005L07Rik | 0.515948 | 0.005196 | 0.25808 |
| 17459236 | Mad2l1 | 0.253832 | 0.005235 | 0.258886 |
| 17497811 | Mir210 | -0.46245 | 0.005246 | 0.259143 |
| 17217768 | Gm26568 | -0.45504 | 0.005271 | 0.259154 |
| 17284660 | Ighm; Ighv1-82 | 1.458407 | 0.005278 | 0.259154 |
| 17288114 | Gm16133 | -0.26872 | 0.005278 | 0.259154 |
| 17325159 | Mylk | -0.27672 | 0.005293 | 0.259612 |
| 17421875 | Slc2a5 | 0.353257 | 0.005322 | 0.259773 |
| 17414482 | Snx30 | 0.274426 | 0.005338 | 0.260157 |
| 17407531 | Gm10696 | -0.27431 | 0.005373 | 0.260679 |
| 17419880 | Ifnlr1 | 0.318076 | 0.005367 | 0.260679 |
| 17481284 | Trim34b; Trim34a | 0.513568 | 0.005378 | 0.260679 |
| 17269903 | Ptges3l | -0.71018 | 0.005454 | 0.26234 |
| 17382755 | Glt6d1 | -0.24839 | 0.005438 | 0.26234 |
| 17419237 | Pum1 | 0.229318 | 0.005457 | 0.26234 |
| 17500996 | Acsl1 | 0.310599 | 0.005427 | 0.26234 |
| 17217169 | Mir135b | -0.33664 | 0.00548 | 0.262847 |
| 17483048 | Apobr | -0.36021 | 0.00548 | 0.262847 |
| 17342038 | Fahd1 | 0.259962 | 0.005498 | 0.263217 |
| 17245902 | Shmt2 | 0.29664 | 0.005525 | 0.26345 |
| 17275718 | Mia2 | 0.282949 | 0.005513 | 0.26345 |
| 17364133 | Ifit1bl2 | 0.362326 | 0.005524 | 0.26345 |
| 17282117 | Plek2 | 0.52028 | 0.00561 | 0.263739 |
| 17284609 | Ighv1-62-3 | 1.351655 | 0.00561 | 0.263739 |
| 17324859 | Tnk2 | 0.314839 | 0.005612 | 0.263739 |
| 17359411 | Zfp518a; Mir8092 | 0.227418 | 0.005633 | 0.263739 |
| 17407542 | Oaz3 | -0.25024 | 0.00555 | 0.263739 |
| 17469016 | Wnt7a | -0.27075 | 0.005601 | 0.263739 |
| 17483007 | Sbk1 | 0.278101 | 0.005565 | 0.263739 |
| 17484587 | Cyp2e1 | 0.25697 | 0.005628 | 0.263739 |

Supplementary Table S2

| *Row.names* | *Gene.Symbol* | *logFC* | *P.Value* | *adj.P.Val* |
|---|---|---|---|---|
| 17514320 | 2610044O15Rik8 | 0.318346 | 0.00563 | 0.263739 |
| 17283393 | Trip11 | 0.252541 | 0.005705 | 0.264417 |
| 17344736 | H2-M10.3 | -0.24576 | 0.005654 | 0.264417 |
| 17353042 | Klhl14 | -0.32579 | 0.005682 | 0.264417 |
| 17381277 | Gm13175 | -0.34168 | 0.005678 | 0.264417 |
| 17390186 | Tmem87a | 0.280076 | 0.005704 | 0.264417 |
| 17419265 | Gm12971 | 0.319595 | 0.00569 | 0.264417 |
| 17531923 | Cmtm7 | -0.32275 | 0.005697 | 0.264417 |
| 17542220 | Gabra3 | -0.30784 | 0.005664 | 0.264417 |
| 17220797 | Tmem206 | 0.305293 | 0.005757 | 0.264519 |
| 17230451 | Gm5069 | -0.28776 | 0.005715 | 0.264519 |
| 17233001 | Pdss2 | 0.301182 | 0.005779 | 0.264519 |
| 17273591 | Kif3c | -0.26273 | 0.00579 | 0.264519 |
| 17284571 | Ighv1-50 | 0.329904 | 0.005755 | 0.264519 |
| 17294267 | Slc6a19 | 0.486907 | 0.005815 | 0.264519 |
| 17368009 | Abca2 | 0.25426 | 0.005733 | 0.264519 |
| 17385719 | Dpp4 | 0.314983 | 0.005813 | 0.264519 |
| 17425708 | AI314180 | 0.267512 | 0.005767 | 0.264519 |
| 17450180 | Fam175a | 0.279295 | 0.005786 | 0.264519 |
| 17467469 | Igkv4-53 | 0.487698 | 0.00576 | 0.264519 |
| 17476752 | Rhpn2 | 0.371463 | 0.005797 | 0.264519 |
| 17514719 | Fam76b | 0.337753 | 0.005765 | 0.264519 |
| 17525839 | Gm16096 | -0.59774 | 0.005769 | 0.264519 |
| 17325637 | B4galt4 | -0.36272 | 0.00584 | 0.265347 |
| 17229891 | Atp1a2 | -0.4686 | 0.005851 | 0.265542 |
| 17278056 | Rin3 | -0.31059 | 0.00593 | 0.266431 |
| 17336446 | Psmb8 | -0.33983 | 0.005878 | 0.266431 |
| 17378575 | Cnbd2 | 0.312858 | 0.005922 | 0.266431 |
| 17435422 | Nub1 | 0.2826 | 0.005894 | 0.266431 |
| 17462841 | Gm23554 | -0.4098 | 0.005917 | 0.266431 |
| 17518563 | Rasl12 | -0.30304 | 0.005913 | 0.266431 |
| 17539107 | Gm6568 | 0.388423 | 0.005935 | 0.266431 |
| 17408123 | Cd160 | -0.27978 | 0.005961 | 0.266971 |
| 17494172 | Olfr605 | -0.36141 | 0.00598 | 0.266971 |
| 17518417 | Vwa9 | -0.42338 | 0.005957 | 0.266971 |
| 17222072 | Ccdc115 | 0.300004 | 0.00604 | 0.267043 |
| 17236752 | Gm24433 | -0.4642 | 0.006017 | 0.267043 |
| 17237887 | Cyp27b1 | 1.415279 | 0.00614 | 0.267043 |
| 17246392 | Olfr767 | -0.32509 | 0.006105 | 0.267043 |
| 17300279 | Mmp14 | -0.45005 | 0.006155 | 0.267043 |
| 17309802 | Oxct1 | 0.247733 | 0.006091 | 0.267043 |
| 17322507 | Dnase1 | 0.381779 | 0.005999 | 0.267043 |

Supplementary Table S2

| *Row.names* | *Gene.Symbol* | *logFC* | *P.Value* | *adj.P.Val* |
|---|---|---|---|---|
| 17330218 | Slc15a2 | 0.503971 | 0.006091 | 0.267043 |
| 17357906 | Olfr1461 | -0.2647 | 0.006159 | 0.267043 |
| 17378827 | Lbp | -0.26117 | 0.006134 | 0.267043 |
| 17398321 | Gm24544 | 0.473736 | 0.006149 | 0.267043 |
| 17398416 | Pdgfc | 0.269535 | 0.006017 | 0.267043 |
| 17437887 | Limch1 | 0.377962 | 0.006162 | 0.267043 |
| 17459291 | Igkv1-135 | 1.675663 | 0.006146 | 0.267043 |
| 17467209 | Fam13a | 0.329392 | 0.006027 | 0.267043 |
| 17509504 | Gm23329 | -0.52931 | 0.006052 | 0.267043 |
| 17511069 | Nanos3 | -0.31779 | 0.006033 | 0.267043 |
| 17535841 | Plxna3 | -0.23574 | 0.006132 | 0.267043 |
| 17287827 | Tgfbi | -0.25251 | 0.006247 | 0.267606 |
| 17288194 | Zfp369 | 0.320741 | 0.006281 | 0.267606 |
| 17324643 | Bdh1 | 0.574448 | 0.006291 | 0.267606 |
| 17330649 | Gm17783 | -0.44473 | 0.006295 | 0.267606 |
| 17408156 | Gja8 | -0.275 | 0.006253 | 0.267606 |
| 17419268 | Gm24091; Gm23711 | 0.437479 | 0.006301 | 0.267606 |
| 17419270 | Gm23711 | 0.437479 | 0.006301 | 0.267606 |
| 17451629 | Hnf1a | 0.290745 | 0.006266 | 0.267606 |
| 17455903 | Asb4 | -0.2843 | 0.006205 | 0.267606 |
| 17457802 | Trbj2-7; Tcrb-J; Trbv13-2; Trbv29 | -0.55867 | 0.006209 | 0.267606 |
| 17502690 | Gm23981 | -0.39509 | 0.006271 | 0.267606 |
| 17533959 | Eif2s3x | 0.420859 | 0.006231 | 0.267606 |
| 17542419 | L1cam | 0.404487 | 0.006218 | 0.267606 |
| 17218042 | Gm5263 | 0.412579 | 0.006365 | 0.268359 |
| 17284593 | Ighv8-8 | 1.546727 | 0.006367 | 0.268359 |
| 17456339 | Cped1 | -0.48785 | 0.006342 | 0.268359 |
| 17467624 | Kdm3a | 0.301404 | 0.006361 | 0.268359 |
| 17491999 | Lrrc28 | 0.252709 | 0.006345 | 0.268359 |
| 17539548 | Piga | 0.362212 | 0.006364 | 0.268359 |
| 17489873 | Uri1 | 0.339936 | 0.006394 | 0.268936 |
| 17261425 | Stc2 | -0.3565 | 0.006412 | 0.269019 |
| 17431717 | Rap1gapos | -0.29347 | 0.006416 | 0.269019 |
| 17411235 | Ak5 | -0.37397 | 0.00643 | 0.269051 |
| 17480986 | Numa1 | 0.246447 | 0.006436 | 0.269051 |
| 17371053 | Gm13572 | -0.25565 | 0.006466 | 0.269486 |
| 17269263 | Krtap16-1 | 0.278244 | 0.006524 | 0.269796 |
| 17283445 | Lgmn | 0.258748 | 0.006528 | 0.269796 |
| 17284387 | Ighv5-6; Igh-V7183 | 0.730703 | 0.006524 | 0.269796 |
| 17377778 | Id1 | -0.66318 | 0.006512 | 0.269796 |
| 17499660 | Gm31464 | 0.548865 | 0.006532 | 0.269796 |
| 17508735 | Tnks | 0.411036 | 0.006507 | 0.269796 |

Supplementary Table S2

| *Row.names* | *Gene.Symbol* | *logFC* | *P.Value* | *adj.P.Val* |
|---|---|---|---|---|
| 17544244 | Chm | 0.238244 | 0.006523 | 0.269796 |
| 17437565 | Gm22536 | -0.45395 | 0.006549 | 0.269971 |
| 17500441 | Purg | 0.434401 | 0.006544 | 0.269971 |
| 17491716 | Mir344e | -0.23182 | 0.006566 | 0.270056 |
| 17512153 | Cdh11 | -0.37788 | 0.00658 | 0.270056 |
| 17429441 | Guca2b | -0.28597 | 0.006604 | 0.27009 |
| 17539824 | Mir188 | -0.60346 | 0.006594 | 0.27009 |
| 17307215 | Zdhhc20 | 0.390373 | 0.006623 | 0.270585 |
| 17238097 | Tmem194 | 0.349628 | 0.00664 | 0.270759 |
| 17470580 | Apobec1 | 0.319077 | 0.006637 | 0.270759 |
| 17459449 | Foxi3 | -0.43752 | 0.006664 | 0.271448 |
| 17508779 | Cldn23 | -0.30961 | 0.006675 | 0.271629 |
| 17353260 | Gm26823 | -0.34565 | 0.006691 | 0.271767 |
| 17499831 | Defb13 | -0.27188 | 0.006689 | 0.271767 |
| 17236638 | Fgd6 | -0.29906 | 0.00671 | 0.272063 |
| 17301340 | Gm26225 | -0.33665 | 0.006712 | 0.272063 |
| 17229412 | Lrrc52 | -0.33302 | 0.006736 | 0.272254 |
| 17436907 | Afap1 | 0.291229 | 0.006724 | 0.272254 |
| 17477237 | Klk8 | -0.3336 | 0.006759 | 0.272908 |
| 17237665 | Llph | -0.35189 | 0.006782 | 0.273018 |
| 17509235 | Trappc11 | 0.2275 | 0.006782 | 0.273018 |
| 17326405 | Dcbld2 | 0.28896 | 0.006815 | 0.273195 |
| 17362467 | Tmem179b | -0.27773 | 0.006808 | 0.273195 |
| 17436839 | Acox3 | 0.296587 | 0.006805 | 0.273195 |
| 17530227 | n-R5s89 | -0.52918 | 0.006819 | 0.273195 |
| 17364139 | Slc16a12 | 0.286907 | 0.006869 | 0.274145 |
| 17482110 | Xylt1 | 0.373194 | 0.006877 | 0.274145 |
| 17510166 | Mast3 | 0.281894 | 0.006896 | 0.274145 |
| 17537373 | Gm22590 | -0.26394 | 0.006895 | 0.274145 |
| 17517284 | 4930510E17Rik | 0.276253 | 0.006907 | 0.274314 |
| 17294246 | D630045M09Rik | -0.5481 | 0.006928 | 0.274387 |
| 17337945 | Gm17080 | -0.24254 | 0.006937 | 0.274387 |
| 17378964 | Dhx35 | 0.309046 | 0.006944 | 0.274387 |
| 17514399 | Gucy1a2 | -0.38573 | 0.006935 | 0.274387 |
| 17216339 | Tnfrsf11a | 0.299509 | 0.006973 | 0.274565 |
| 17396315 | Tbl1xr1 | 0.389851 | 0.006967 | 0.274565 |
| 17403790 | Erich3 | 0.289161 | 0.006962 | 0.274565 |
| 17486474 | Zscan4b | -0.32605 | 0.007036 | 0.275275 |
| 17520662 | Prr23a1 | -0.27735 | 0.007036 | 0.275275 |
| 17521171 | Tlr9 | -0.29918 | 0.007026 | 0.275275 |
| 17221510 | Gm16070 | -0.20787 | 0.007088 | 0.2757 |
| 17266751 | Gm17268 | -0.49074 | 0.007062 | 0.2757 |

Supplementary Table S2

| *Row.names* | *Gene.Symbol* | *logFC* | *P.Value* | *adj.P.Val* |
|---|---|---|---|---|
| 17277195 | Ptgr2 | 0.31443 | 0.007078 | 0.2757 |
| 17362689 | Sdhaf2 | 0.324714 | 0.007086 | 0.2757 |
| 17375454 | Slc28a2 | -0.47922 | 0.007084 | 0.2757 |
| 17218722 | 2810442N19Rik | -0.25899 | 0.007115 | 0.275775 |
| 17222800 | Slc39a10 | 0.247414 | 0.007139 | 0.275775 |
| 17290033 | Arl15 | 0.262659 | 0.007136 | 0.275775 |
| 17298731 | Gdf10 | -0.43716 | 0.007125 | 0.275775 |
| 17406285 | D930015E06Rik | 0.38493 | 0.007139 | 0.275775 |
| 17474994 | Zfp428 | -0.35835 | 0.0071 | 0.275775 |
| 17519052 | Gm24615 | -0.2528 | 0.007133 | 0.275775 |
| 17522479 | Prss45 | -0.31799 | 0.007144 | 0.275775 |
| 17395099 | Ankrd60 | -0.23227 | 0.007182 | 0.276213 |
| 17411411 | Gm15577 | -0.26615 | 0.007178 | 0.276213 |
| 17231423 | Akap12 | -0.31073 | 0.007564 | 0.277168 |
| 17232778 | Mettl24 | 0.293078 | 0.007533 | 0.277168 |
| 17234339 | Adora2a | -0.29987 | 0.007636 | 0.277168 |
| 17236339 | Gnptab | 0.239974 | 0.007582 | 0.277168 |
| 17241257 | Col13a1 | -0.31149 | 0.007262 | 0.277168 |
| 17253450 | Mir451a | -0.2405 | 0.007452 | 0.277168 |
| 17264960 | Fgf11 | -0.37189 | 0.007619 | 0.277168 |
| 17265818 | Olfr410 | -0.29034 | 0.007446 | 0.277168 |
| 17268729 | Fbxl20 | 0.259247 | 0.007352 | 0.277168 |
| 17274875 | Nampt | 0.328409 | 0.007473 | 0.277168 |
| 17278767 | Gm23600 | -0.29065 | 0.007612 | 0.277168 |
| 17308875 | Zfp957 | -0.33898 | 0.007452 | 0.277168 |
| 17320809 | Gm25546 | -0.31594 | 0.007428 | 0.277168 |
| 17335234 | Scube3 | -0.52195 | 0.007554 | 0.277168 |
| 17352056 | Epg5 | 0.253412 | 0.007338 | 0.277168 |
| 17362342 | Lgals12 | 0.242315 | 0.007448 | 0.277168 |
| 17369076 | Pkn3 | -0.29552 | 0.007572 | 0.277168 |
| 17372753 | Olfr1009 | -0.31529 | 0.007587 | 0.277168 |
| 17372771 | Olfr1022 | -0.32798 | 0.007498 | 0.277168 |
| 17381171 | Pcmtd2 | 0.32208 | 0.007364 | 0.277168 |
| 17389132 | Platr8 | -0.23233 | 0.007616 | 0.277168 |
| 17398499 | Fga | 0.309638 | 0.007601 | 0.277168 |
| 17400510 | Sf3b4 | 0.332975 | 0.007493 | 0.277168 |
| 17400549 | Gm20634 | -0.65873 | 0.007488 | 0.277168 |
| 17402996 | Ddit4l | -0.29247 | 0.007421 | 0.277168 |
| 17406279 | Tlr2 | 0.232824 | 0.00762 | 0.277168 |
| 17418881 | A3galt2 | -0.23503 | 0.007505 | 0.277168 |
| 17425615 | Txn1 | -0.3333 | 0.007473 | 0.277168 |
| 17427441 | Gm10192 | -0.26784 | 0.007446 | 0.277168 |

Supplementary Table S2

| *Row.names* | *Gene.Symbol* | *logFC* | *P.Value* | *adj.P.Val* |
|---|---|---|---|---|
| 17429430 | AA415398 | 0.253387 | 0.007609 | 0.277168 |
| 17436629 | Add1 | 0.239722 | 0.007388 | 0.277168 |
| 17440218 | Gm10419 | -0.24429 | 0.007252 | 0.277168 |
| 17444930 | Gm15409 | -0.37459 | 0.007377 | 0.277168 |
| 17446760 | Ost4 | -0.21011 | 0.00745 | 0.277168 |
| 17448170 | Gm22273 | -0.51103 | 0.007632 | 0.277168 |
| 17458956 | Vmn1r19 | 0.882645 | 0.007229 | 0.277168 |
| 17467302 | Vmn1r36 | -0.47181 | 0.007535 | 0.277168 |
| 17470187 | Zfp9 | -0.21775 | 0.007344 | 0.277168 |
| 17479974 | Nox4 | 0.330952 | 0.007584 | 0.277168 |
| 17503507 | Lonp2 | 0.262203 | 0.007328 | 0.277168 |
| 17504374 | Gm25160 | -0.40067 | 0.007426 | 0.277168 |
| 17505283 | Wwp2 | 0.286865 | 0.007408 | 0.277168 |
| 17514926 | Gm23048 | -0.43524 | 0.007604 | 0.277168 |
| 17518238 | Calml4 | 0.388195 | 0.007359 | 0.277168 |
| 17525955 | Sc5d | 0.297267 | 0.007485 | 0.277168 |
| 17541681 | Gpc3 | -0.40786 | 0.007434 | 0.277168 |
| 17310637 | Fam134b | 0.287983 | 0.00767 | 0.278166 |
| 17250322 | 1810063I02Rik | -0.33183 | 0.007717 | 0.279616 |
| 17220059 | Cdc42bpa | 0.261127 | 0.00779 | 0.280267 |
| 17250317 | Gm12714 | -0.20971 | 0.007803 | 0.280267 |
| 17268404 | Pnpo | 0.231543 | 0.007774 | 0.280267 |
| 17284628 | Ighv1-69 | 1.087918 | 0.007823 | 0.280267 |
| 17340050 | Plekhh2 | -0.27343 | 0.007764 | 0.280267 |
| 17400291 | Fam63a | 0.219015 | 0.007817 | 0.280267 |
| 17424913 | Fbxo10 | 0.32317 | 0.00781 | 0.280267 |
| 17430288 | Azin2 | 0.271744 | 0.007745 | 0.280267 |
| 17434285 | Gm1987; Ccl21a | -0.41949 | 0.007757 | 0.280267 |
| 17515819 | Ets1 | -0.20852 | 0.007782 | 0.280267 |
| 17211572 | Rab23 | 0.311288 | 0.007855 | 0.28035 |
| 17271259 | Wipi1 | 0.256498 | 0.007864 | 0.28035 |
| 17312209 | Ly6e | -0.34582 | 0.007834 | 0.28035 |
| 17539700 | Mid1 | -0.64494 | 0.007864 | 0.28035 |
| 17339499 | Snord53 | -0.24723 | 0.007901 | 0.28112 |
| 17315002 | Mettl7a1 | 0.243719 | 0.007931 | 0.281198 |
| 17367576 | Acbd5 | 0.239228 | 0.007931 | 0.281198 |
| 17387565 | Lrrc55 | -0.25377 | 0.007938 | 0.281198 |
| 17390296 | Ttbk2; Cdan1 | 0.308542 | 0.007944 | 0.281198 |
| 17428553 | Rad54l | -0.25547 | 0.007927 | 0.281198 |
| 17535448 | Zfp185 | 0.479823 | 0.007933 | 0.281198 |
| 17314085 | Ppp6r2 | 0.233897 | 0.00798 | 0.281345 |
| 17321422 | Rhebl1 | 0.240946 | 0.008002 | 0.281345 |

Supplementary Table S2

| *Row.names* | *Gene.Symbol* | *logFC* | *P.Value* | *adj.P.Val* |
|---|---|---|---|---|
| 17413162 | 4930578G10Rik | -0.40367 | 0.007998 | 0.281345 |
| 17419741 | Slc30a2 | 0.315372 | 0.007968 | 0.281345 |
| 17542583 | Tex28 | -0.29005 | 0.007998 | 0.281345 |
| 17343461 | Zfp871 | 0.488934 | 0.00801 | 0.281359 |
| 17499949 | Dkk4 | -0.226 | 0.008028 | 0.281523 |
| 17276743 | Plekhh1 | 0.433114 | 0.008047 | 0.281873 |
| 17319719 | Tcf20 | -0.2335 | 0.008067 | 0.281953 |
| 17364218 | Cpeb3 | 0.364484 | 0.008107 | 0.282612 |
| 17481171 | Olfr584 | -0.38299 | 0.008121 | 0.282644 |
| 17421168 | Gm22039 | -0.41898 | 0.008146 | 0.282649 |
| 17457796 | Tcrb-J; Trbj2-4 | -0.42427 | 0.008149 | 0.282649 |
| 17303386 | Dnase1l3 | -0.26339 | 0.008281 | 0.28276 |
| 17325497 | Hgd | -0.23376 | 0.008262 | 0.28276 |
| 17326640 | Rbm11 | 0.323163 | 0.008221 | 0.28276 |
| 17344132 | Hspa1a | -0.55726 | 0.008294 | 0.28276 |
| 17349520 | Ctnna1 | 0.218998 | 0.008219 | 0.28276 |
| 17372564 | Gm13691; Gm13697; Gm13698; Gm13694; Gm13693; Gm13696 | 0.247629 | 0.008286 | 0.28276 |
| 17372566 | Gm13691; Gm13697; Gm13698; Gm13694; Gm13693; Gm13696 | 0.247629 | 0.008286 | 0.28276 |
| 17372568 | Gm13695 | 0.247629 | 0.008286 | 0.28276 |
| 17372570 | Gm13691; Gm13697; Gm13698; Gm13694; Gm13693; Gm13696 | 0.247629 | 0.008286 | 0.28276 |
| 17372572 | Gm13691; Gm13697; Gm13698; Gm13694; Gm13693; Gm13696 | 0.247629 | 0.008286 | 0.28276 |
| 17372574 | Gm13691; Gm13697; Gm13698; Gm13694; Gm13693; Gm13696 | 0.247629 | 0.008286 | 0.28276 |
| 17372576 | Gm13691; Gm13697; Gm13698; Gm13694; Gm13693; Gm13696 | 0.247629 | 0.008286 | 0.28276 |
| 17402468 | Gm25501 | 0.407409 | 0.008258 | 0.28276 |
| 17448373 | LOC102634968 | -0.37584 | 0.008234 | 0.28276 |
| 17475170 | Zfp574 | 0.26303 | 0.008205 | 0.28276 |
| 17491026 | Emp3 | -0.31116 | 0.008185 | 0.28276 |
| 17507945 | Defb11 | -0.63595 | 0.008296 | 0.28276 |
| 17238179 | Rdh18-ps; Rdh16 | 1.035719 | 0.008323 | 0.283316 |
| 17299582 | Eddm3b | 2.484652 | 0.008326 | 0.283316 |
| 17240880 | Man1a | 0.268245 | 0.008338 | 0.283514 |
| 17224360 | Prkag3 | 0.3426 | 0.008412 | 0.28361 |
| 17236792 | 4930556N09Rik | -0.21246 | 0.008382 | 0.28361 |
| 17245466 | Tbc1d30 | 0.272517 | 0.008395 | 0.28361 |
| 17270558 | Lyzl6 | -0.25699 | 0.008442 | 0.28361 |
| 17280577 | Hbp1 | 0.308273 | 0.00836 | 0.28361 |
| 17285446 | Gm25020 | 0.463255 | 0.008408 | 0.28361 |
| 17309397 | Tgds | 0.425058 | 0.008458 | 0.28361 |

Supplementary Table S2

| *Row.names* | *Gene.Symbol* | *logFC* | *P.Value* | *adj.P.Val* |
|---|---|---|---|---|
| 17369721 | Pomt1 | 0.316867 | 0.008421 | 0.28361 |
| 17435584 | Insig1 | 0.45141 | 0.008454 | 0.28361 |
| 17467392 | Igkv19-93 | 0.622867 | 0.008462 | 0.28361 |
| 17471231 | Prmt8 | -0.24469 | 0.008456 | 0.28361 |
| 17504293 | Mmp15 | -0.25928 | 0.008465 | 0.28361 |
| 17372892 | Olfr1153 | -0.36429 | 0.008474 | 0.283677 |
| 17224386 | Cryba2 | -0.31067 | 0.008516 | 0.284869 |
| 17377744 | Rem1 | -0.31324 | 0.008532 | 0.284944 |
| 17453454 | Eln | -0.37294 | 0.008531 | 0.284944 |
| 17315570 | Nckap1l | -0.25615 | 0.008578 | 0.285555 |
| 17531130 | Apeh | 0.311953 | 0.008578 | 0.285555 |
| 17324513 | Cldn16 | -0.40005 | 0.008608 | 0.286262 |
| 17486254 | 2810047C21Rik1 | -0.52692 | 0.008621 | 0.286262 |
| 17525673 | Panx3 | -0.23954 | 0.008627 | 0.286262 |
| 17391150 | Trpm7 | 0.247965 | 0.008636 | 0.286328 |
| 17404180 | Car1 | -0.22291 | 0.008687 | 0.287113 |
| 17227480 | Gpr25 | -0.27613 | 0.008752 | 0.287403 |
| 17234910 | Cdc34 | -0.26998 | 0.008747 | 0.287403 |
| 17350925 | Iigp1 | -0.48614 | 0.008719 | 0.287403 |
| 17364917 | Hpse2 | -0.33332 | 0.008735 | 0.287403 |
| 17385985 | Ttc21b | 0.320422 | 0.008728 | 0.287403 |
| 17387742 | Olfr259 | -0.47643 | 0.008712 | 0.287403 |
| 17467540 | Igkv6-20 | 1.121917 | 0.00875 | 0.287403 |
| 17391997 | Gpcpd1 | 0.465963 | 0.008766 | 0.287556 |
| 17409753 | Agl | 0.252803 | 0.00877 | 0.287556 |
| 17431708 | Ldlrad2 | -0.36485 | 0.008778 | 0.287571 |
| 17346317 | Lonp1 | 0.282955 | 0.008786 | 0.2876 |
| 17245481 | Rassf3 | 0.514063 | 0.008799 | 0.287804 |
| 17233037 | Gm9034 | -0.43787 | 0.008866 | 0.288143 |
| 17242150 | Col6a2 | -0.37433 | 0.008868 | 0.288143 |
| 17278470 | Vrk1 | -0.27115 | 0.008876 | 0.288143 |
| 17299518 | Parp2 | 0.386739 | 0.00887 | 0.288143 |
| 17355788 | Gm24720 | 0.33009 | 0.008872 | 0.288143 |
| 17358891 | Pcgf5 | 0.350053 | 0.008859 | 0.288143 |
| 17394267 | Tnnc2 | -0.20997 | 0.008854 | 0.288143 |
| 17486099 | Zim1 | -0.32059 | 0.008868 | 0.288143 |
| 17496532 | Pagr1a; Prrt2 | -0.25402 | 0.008879 | 0.288143 |
| 17518777 | Gm15563 | -0.33959 | 0.008835 | 0.288143 |
| 17216779 | Acmsd | -0.29598 | 0.009014 | 0.288267 |
| 17298626 | Fam170b | -0.33565 | 0.008988 | 0.288267 |
| 17344720 | H2-M10.2 | -0.34364 | 0.008957 | 0.288267 |
| 17350932 | BC023105 | -0.44439 | 0.009029 | 0.288267 |

Supplementary Table S2

| *Row.names* | *Gene.Symbol* | *logFC* | *P.Value* | *adj.P.Val* |
|---|---|---|---|---|
| 17366744 | Mir669a-3 | -0.693 | 0.008894 | 0.288267 |
| 17443985 | Cyp2w1 | -0.28647 | 0.009005 | 0.288267 |
| 17482450 | Abca16 | -0.19919 | 0.008961 | 0.288267 |
| 17492066 | Nr2f2 | -0.2608 | 0.008957 | 0.288267 |
| 17506042 | Cdh13 | -0.417 | 0.008938 | 0.288267 |
| 17507487 | Gm15875 | -0.38734 | 0.009017 | 0.288267 |
| 17519259 | Ccpg1 | 0.272399 | 0.009 | 0.288267 |
| 17539750 | Gm3701 | -0.29811 | 0.009 | 0.288267 |
| 17410636 | Mttp | 0.439384 | 0.009047 | 0.288312 |
| 17498080 | Gm10013 | -0.32702 | 0.009053 | 0.288312 |
| 17396132 | E2f5 | -0.2339 | 0.009069 | 0.288437 |
| 17471140 | Ntf3 | -0.1979 | 0.009106 | 0.289157 |
| 17219516 | Gm17224 | 0.329995 | 0.009161 | 0.290229 |
| 17247695 | Xpo1 | -0.27753 | 0.009169 | 0.290266 |
| 17297383 | Gm22350 | 0.331257 | 0.009176 | 0.290266 |
| 17424461 | Gm1987; Ccl21a | -0.42204 | 0.009184 | 0.290311 |
| 17301061 | Mipep | 0.213856 | 0.009222 | 0.291265 |
| 17308280 | Ppp3cc | -0.36977 | 0.00923 | 0.291324 |
| 17272895 | Eif4a3 | -0.39464 | 0.00925 | 0.291661 |
| 17532645 | mt-Ty | 0.811522 | 0.009255 | 0.291661 |
| 17461807 | Irak2 | 0.276974 | 0.009263 | 0.291687 |
| 17340845 | Igf2r | 0.496193 | 0.009278 | 0.29194 |
| 17234175 | Ipmk | 0.256389 | 0.009294 | 0.292005 |
| 17397750 | Tm4sf4 | -0.30137 | 0.009294 | 0.292005 |
| 17533332 | Gm4906 | -0.37729 | 0.009303 | 0.292044 |
| 17530348 | Dnajc13 | 0.273859 | 0.009313 | 0.292139 |
| 17438788 | Rufy3 | 0.21186 | 0.009321 | 0.292158 |
| 17211526 | Gm5524 | 0.372708 | 0.009347 | 0.292316 |
| 17216063 | Bok | -0.21314 | 0.009386 | 0.293276 |
| 17314115 | Adm2 | -0.37493 | 0.009392 | 0.293276 |
| 17217182 | Nuak2 | -0.63456 | 0.009399 | 0.29329 |
| 17277939 | 9030617O03Rik; Gm36572 | -0.29146 | 0.009428 | 0.293522 |
| 17290781 | Sugct | 0.265067 | 0.009442 | 0.293522 |
| 17365728 | Gpam | 0.222364 | 0.009429 | 0.293522 |
| 17237564 | Mdm1 | -0.31104 | 0.009457 | 0.293556 |
| 17234850 | Olfr8 | -0.29781 | 0.009564 | 0.293843 |
| 17236878 | Galnt4 | 0.227567 | 0.009505 | 0.293843 |
| 17265935 | Rpa1 | 0.257089 | 0.009584 | 0.293843 |
| 17281791 | L3hypdh | 0.253524 | 0.009587 | 0.293843 |
| 17305167 | Ghitm | 0.229222 | 0.009556 | 0.293843 |
| 17397297 | 3110057O12Rik | 0.354798 | 0.009573 | 0.293843 |
| 17405998 | Platr10 | -0.29306 | 0.009491 | 0.293843 |

Supplementary Table S2

| *Row.names* | *Gene.Symbol* | *logFC* | *P.Value* | *adj.P.Val* |
|---|---|---|---|---|
| 17420895 | Gm26226 | -0.52874 | 0.009516 | 0.293843 |
| 17430722 | Gm12992 | -0.2241 | 0.009544 | 0.293843 |
| 17501299 | Gm25379 | -0.4534 | 0.009533 | 0.293843 |
| 17262297 | Gm12195 | -0.25901 | 0.009633 | 0.294416 |
| 17279148 | Eif5 | 0.230603 | 0.009624 | 0.294416 |
| 17409461 | Gm9857 | -0.24159 | 0.009635 | 0.294416 |
| 17518809 | Gm19299 | -0.28145 | 0.009633 | 0.294416 |
| 17363025 | Stx3 | 0.382752 | 0.009714 | 0.296039 |
| 17421960 | Gm23209 | 0.563549 | 0.009716 | 0.296039 |
| 17311551 | Col14a1 | -0.26572 | 0.00974 | 0.296111 |
| 17211243 | Terf1 | 0.246604 | 0.009762 | 0.296552 |
| 17494469 | Olfr685 | -0.25402 | 0.009793 | 0.297058 |
| 17269595 | Dhx58 | 0.23785 | 0.009838 | 0.297322 |
| 17424215 | Gm26049 | 0.287231 | 0.009864 | 0.297322 |
| 17456051 | n-R5s160 | -0.40217 | 0.009859 | 0.297322 |
| 17465924 | Svopl | 0.394573 | 0.009872 | 0.297322 |
| 17484030 | Hmx3 | -0.28705 | 0.009817 | 0.297322 |
| 17501427 | Cbr4 | 0.243053 | 0.009835 | 0.297322 |
| 17516194 | Vwa5a | 0.333008 | 0.009817 | 0.297322 |
| 17527934 | Kif23 | -0.38361 | 0.009849 | 0.297322 |
| 17547965 | Cbr1 | -0.31921 | 0.009874 | 0.297322 |
| 17385065 | Gm13490 | -0.21808 | 0.009896 | 0.297774 |
| 17284360 | Ighm; Ighj1; Igh-VJ558; Ighg3; Igh-V7183 | 0.947036 | 0.009963 | 0.298228 |
| 17400124 | Selenbp2 | 0.236477 | 0.009941 | 0.298228 |
| 17403992 | Gm23592 | -0.24292 | 0.009946 | 0.298228 |
| 17496664 | Zfp747 | -0.40811 | 0.009955 | 0.298228 |
| 17539303 | Phka2 | 0.221149 | 0.009969 | 0.298228 |
| 17539966 | Gata1 | -0.22268 | 0.009923 | 0.298228 |
| 17266882 | LOC102635154 | -0.29729 | 0.010027 | 0.298274 |
| 17281622 | Cdkl1 | 0.359199 | 0.010045 | 0.298274 |
| 17318083 | Ly6a | -0.43557 | 0.010037 | 0.298274 |
| 17348282 | Colec12 | -0.29847 | 0.010036 | 0.298274 |
| 17353940 | Pcdh12 | -0.32383 | 0.010002 | 0.298274 |
| 17400989 | 4930406D18Rik | -0.28748 | 0.010057 | 0.298274 |
| 17412191 | Coq3 | 0.226367 | 0.010027 | 0.298274 |
| 17418128 | Trit1 | 0.336438 | 0.009986 | 0.298274 |
| 17478181 | Kcnc1 | -0.30567 | 0.010016 | 0.298274 |
| 17484151 | Fank1 | -0.25527 | 0.010026 | 0.298274 |
| 17420831 | LOC102637278; Gm13017 | 0.282504 | 0.010085 | 0.298906 |
| 17283082 | Lysmd1 | 0.236659 | 0.010138 | 0.298951 |
| 17307134 | Cryl1 | 0.236723 | 0.010122 | 0.298951 |
| 17315669 | AW549877 | 0.273977 | 0.010145 | 0.298951 |

Supplementary Table S2

| *Row.names* | *Gene.Symbol* | *logFC* | *P.Value* | *adj.P.Val* |
|---|---|---|---|---|
| 17392543 | Gm14110 | -0.40792 | 0.010139 | 0.298951 |
| 17411633 | Gm23860 | -0.24464 | 0.010118 | 0.298951 |
| 17428142 | Osbpl9 | 0.273242 | 0.010132 | 0.298951 |
| 17446093 | Gm22054 | -0.32606 | 0.01016 | 0.299006 |
| 17270410 | Plcd3 | 0.267396 | 0.010207 | 0.299006 |
| 17304145 | 1700054O19Rik | 0.587292 | 0.010221 | 0.299006 |
| 17351204 | Napg | 0.329836 | 0.010191 | 0.299006 |
| 17489151 | Psenen | -0.21508 | 0.01021 | 0.299006 |
| 17226049 | Kdsr | 0.227747 | 0.010241 | 0.299018 |
| 17291361 | Aldh5a1 | 0.231422 | 0.010236 | 0.299018 |
| 17276893 | Srsf5 | 0.299427 | 0.010259 | 0.299137 |
| 17421476 | Gm13139 | 0.582366 | 0.010257 | 0.299137 |
| 17344620 | Gm11127 | 0.646515 | 0.010273 | 0.299327 |
| 17284574 | Ighv1-52 | 0.996838 | 0.010281 | 0.299335 |
| 17513289 | Maf | 0.33536 | 0.01031 | 0.299558 |
| 17336502 | H2-Eb1 | -0.2697 | 0.010321 | 0.299664 |
| 17285938 | Hist1h2aa | -0.44505 | 0.010335 | 0.299866 |
| 17425496 | Tmem245 | 0.353599 | 0.010345 | 0.299944 |

**Supplementary Table S3**. GO term enrichment analysis for up-regulated genes, based on a false discovery rate cut-off value of <0.3.

| *ONTOLOGY* | *ID* | *Description* | *GeneRatio* | *BgRatio* | *pvalue* | *p.adjust* | *qvalue* | *Count* |
|---|---|---|---|---|---|---|---|---|
| CC | GO:0042571 | immunoglobulin complex, circulating | 31/433 | 96/21769 | 2.78E-29 | 1.19E-25 | 1.16E-25 | 31 |
| CC | GO:0019814 | immunoglobulin complex | 31/433 | 100/21769 | 1.17E-28 | 2.52E-25 | 2.45E-25 | 31 |
| MF | GO:0034987 | immunoglobulin receptor binding | 31/433 | 103/21769 | 3.29E-28 | 4.71E-25 | 4.59E-25 | 31 |
| BP | GO:0006910 | phagocytosis, recognition | 30/433 | 110/21769 | 6.52E-26 | 7.00E-23 | 6.82E-23 | 30 |
| MF | GO:0003823 | antigen binding | 33/433 | 158/21769 | 2.49E-24 | 2.14E-21 | 2.09E-21 | 33 |
| BP | GO:0006958 | complement activation, classical pathway | 30/433 | 124/21769 | 3.16E-24 | 2.26E-21 | 2.20E-21 | 30 |
| BP | GO:0050853 | B cell receptor signaling pathway | 31/433 | 141/21769 | 1.22E-23 | 7.50E-21 | 7.30E-21 | 31 |
| BP | GO:0002455 | humoral immune response mediated by circulating immunoglobulin | 30/433 | 138/21769 | 9.24E-23 | 4.96E-20 | 4.83E-20 | 30 |
| BP | GO:0006956 | complement activation | 30/433 | 146/21769 | 5.30E-22 | 2.20E-19 | 2.14E-19 | 30 |
| BP | GO:0050871 | positive regulation of B cell activation | 32/433 | 172/21769 | 5.46E-22 | 2.20E-19 | 2.14E-19 | 32 |
| BP | GO:0072376 | protein activation cascade | 31/433 | 159/21769 | 5.64E-22 | 2.20E-19 | 2.14E-19 | 31 |
| BP | GO:0006911 | phagocytosis, engulfment | 30/433 | 148/21769 | 8.05E-22 | 2.88E-19 | 2.81E-19 | 30 |
| BP | GO:0099024 | plasma membrane invagination | 30/433 | 157/21769 | 4.88E-21 | 1.61E-18 | 1.57E-18 | 30 |
| BP | GO:0010324 | membrane invagination | 30/433 | 164/21769 | 1.82E-20 | 5.58E-18 | 5.44E-18 | 30 |
| BP | GO:0050864 | regulation of B cell activation | 33/433 | 221/21769 | 1.52E-19 | 4.34E-17 | 4.23E-17 | 33 |
| BP | GO:0016064 | immunoglobulin mediated immune response | 31/433 | 212/21769 | 3.67E-18 | 9.85E-16 | 9.60E-16 | 31 |
| BP | GO:0019724 | B cell mediated immunity | 31/433 | 215/21769 | 5.56E-18 | 1.40E-15 | 1.37E-15 | 31 |
| BP | GO:0002377 | immunoglobulin production | 32/433 | 234/21769 | 7.92E-18 | 1.89E-15 | 1.84E-15 | 32 |
| BP | GO:0008037 | cell recognition | 31/433 | 225/21769 | 2.11E-17 | 4.77E-15 | 4.65E-15 | 31 |
| BP | GO:0050851 | antigen receptor-mediated signaling pathway | 32/433 | 249/21769 | 5.07E-17 | 1.09E-14 | 1.06E-14 | 32 |
| BP | GO:0002768 | immune response-regulating cell surface receptor signaling pathway | 34/433 | 292/21769 | 1.12E-16 | 2.29E-14 | 2.23E-14 | 34 |
| BP | GO:0002429 | immune response-activating cell surface receptor signaling pathway | 33/433 | 280/21769 | 2.23E-16 | 4.35E-14 | 4.23E-14 | 33 |
| BP | GO:0002764 | immune response-regulating signaling pathway | 39/433 | 411/21769 | 6.58E-16 | 1.23E-13 | 1.20E-13 | 39 |
| BP | GO:0002757 | immune response-activating signal transduction | 38/433 | 398/21769 | 1.29E-15 | 2.31E-13 | 2.25E-13 | 38 |
| BP | GO:0051251 | positive regulation of lymphocyte activation | 35/433 | 345/21769 | 2.86E-15 | 4.91E-13 | 4.78E-13 | 35 |

Supplementary Table S3

| *ONTOLOGY* | *ID* | *Description* | *GeneRatio* | *BgRatio* | *pvalue* | *p.adjust* | *qvalue* | *Count* |
|---|---|---|---|---|---|---|---|---|
| BP | GO:0006959 | humoral immune response | 32/433 | 288/21769 | 3.52E-15 | 5.81E-13 | 5.66E-13 | 32 |
| BP | GO:0006909 | phagocytosis | 32/433 | 290/21769 | 4.29E-15 | 6.82E-13 | 6.65E-13 | 32 |
| BP | GO:0002440 | production of molecular mediator of immune response | 33/433 | 325/21769 | 1.78E-14 | 2.73E-12 | 2.66E-12 | 33 |
| BP | GO:0002449 | lymphocyte mediated immunity | 34/433 | 368/21769 | 1.12E-13 | 1.66E-11 | 1.62E-11 | 34 |
| BP | GO:0002696 | positive regulation of leukocyte activation | 35/433 | 395/21769 | 1.66E-13 | 2.38E-11 | 2.32E-11 | 35 |
| BP | GO:0042742 | defense response to bacterium | 33/433 | 354/21769 | 2.04E-13 | 2.82E-11 | 2.75E-11 | 33 |
| BP | GO:0002253 | activation of immune response | 38/433 | 467/21769 | 2.13E-13 | 2.86E-11 | 2.78E-11 | 38 |
| BP | GO:0042113 | B cell activation | 33/433 | 357/21769 | 2.58E-13 | 3.36E-11 | 3.27E-11 | 33 |
| BP | GO:0002460 | adaptive immune response based on somatic recombination... | 34/433 | 385/21769 | 4.12E-13 | 5.20E-11 | 5.06E-11 | 34 |
| BP | GO:0050867 | positive regulation of cell activation | 35/433 | 409/21769 | 4.61E-13 | 5.65E-11 | 5.50E-11 | 35 |
| BP | GO:0002443 | leukocyte mediated immunity | 36/433 | 449/21769 | 1.45E-12 | 1.73E-10 | 1.69E-10 | 36 |
| BP | GO:0006631 | fatty acid metabolic process | 24/433 | 363/21769 | 3.25E-07 | 3.77E-05 | 3.67E-05 | 24 |
| CC | GO:0005743 | mitochondrial inner membrane | 23/433 | 409/21769 | 8.71E-06 | 0.00098383 | 0.0009582 | 23 |
| CC | GO:0019866 | organelle inner membrane | 24/433 | 448/21769 | 1.25E-05 | 0.00137428 | 0.00133847 | 24 |
| BP | GO:0006641 | triglyceride metabolic process | 10/433 | 96/21769 | 2.15E-05 | 0.00230401 | 0.00224397 | 10 |
| BP | GO:0006639 | acylglycerol metabolic process | 11/433 | 119/21769 | 2.61E-05 | 0.00273307 | 0.00266185 | 11 |
| BP | GO:0006638 | neutral lipid metabolic process | 11/433 | 121/21769 | 3.05E-05 | 0.00311838 | 0.00303712 | 11 |
| BP | GO:0051186 | cofactor metabolic process | 24/433 | 486/21769 | 4.63E-05 | 0.00462384 | 0.00450335 | 24 |
| MF | GO:0050662 | coenzyme binding | 16/433 | 281/21769 | 0.00017488 | 0.01706642 | 0.0166217 | 16 |
| BP | GO:1901661 | quinone metabolic process | 5/433 | 28/21769 | 0.00020494 | 0.01955539 | 0.01904581 | 5 |
| BP | GO:1902652 | secondary alcohol metabolic process | 10/433 | 126/21769 | 0.00021623 | 0.02018464 | 0.01965867 | 10 |
| BP | GO:0016042 | lipid catabolic process | 16/433 | 296/21769 | 0.00031379 | 0.02866798 | 0.02792094 | 16 |
| BP | GO:0006066 | alcohol metabolic process | 16/433 | 303/21769 | 0.00040607 | 0.03632668 | 0.03538007 | 16 |
| BP | GO:0044282 | small molecule catabolic process | 16/433 | 304/21769 | 0.000421 | 0.0368932 | 0.03593182 | 16 |

**Supplementary Table S4**. GO term enrichment analysis for down-regulated genes, based on a false discovery rate cut-off value of <0.3.

| *ONTOLOGY* | *ID* | *Description* | *GeneRatio* | *BgRatio* | *pvalue* | *p.adjust* | *qvalue* | *Count* |
|---|---|---|---|---|---|---|---|---|
| CC | GO:0031012 | extracellular matrix | 31/329 | 465/21769 | 5.39E-12 | 2.06E-08 | 1.98E-08 | 31 |
| CC | GO:0062023 | collagen-containing extracellular matrix | 23/329 | 356/21769 | 5.95E-09 | 1.14E-05 | 1.10E-05 | 23 |
| MF | GO:0005201 | extracellular matrix structural constituent | 13/329 | 140/21769 | 2.17E-07 | 0.00027676 | 0.00026683 | 13 |
| BP | GO:0030198 | extracellular matrix organization | 17/329 | 265/21769 | 6.57E-07 | 0.000627 | 0.00060451 | 17 |
| BP | GO:0043062 | extracellular structure organization | 17/329 | 308/21769 | 5.08E-06 | 0.00331782 | 0.00319881 | 17 |
| BP | GO:0032963 | collagen metabolic process | 10/329 | 107/21769 | 5.21E-06 | 0.00331782 | 0.00319881 | 10 |
